# Growth directions and stiffness across cell layers determine whether tissues stay smooth or buckle

**DOI:** 10.1101/2023.07.22.549953

**Authors:** Avilash Singh Yadav, Lilan Hong, Patrick M. Klees, Annamaria Kiss, Manuel Petit, Xi He, Iselle M. Barrios, Michelle Heeney, Anabella Maria D. Galang, Richard S. Smith, Arezki Boudaoud, Adrienne H.K. Roeder

**Affiliations:** Institute of Nuclear Agricultural Sciences, Key Laboratory of Nuclear Agricultural Sciences of Ministry of Agriculture and Zhejiang Province, College of Agriculture and Biotechnology, Zhejiang University, Hangzhou 310058, China; Weill Institute for Cell and Molecular Biology and Section of Plant Biology, School of Integrative Plant Sciences, Cornell University, Ithaca, NY 14853, USA; Laboratoire Reproduction et Développement des Plantes, Univ Lyon, ENS de Lyon, UCB Lyon, CNRS, INRAE, INRIA, F-69342 Lyon, France; The John Innes Centre, Norwich NR4 7UH, UK; LadHyX, Ecole Polytechnique, CNRS, IP Paris, 91128 Palaiseau Cedex, France

## Abstract

From smooth shapes to buckles, nature exhibits organs of various shapes and forms. How cells grow to produce smooth shaped leaves and sepals remain unclear. Here, we show that growth along the longitudinal axis during early developmental stages and comparable stiffness across both epidermal layers of Arabidopsis sepals are essential for smoothness, as seen in the wild type. We identified a mutant (*as2-7D*) with ectopic expression of *ASYMMETRIC LEAVES 2* (*AS2*) on the outer epidermis. Our analysis reveals that ectopic *AS2* expression causes the outer epidermis of *as2-7D* sepals to buckle during early stages of sepal development. We show that buckling of the outer epidermis occurs due to conflicting cell growth directions and unequal tissue stiffness across the epidermal layers. Overexpression of cyclin-dependent kinase (CDK) inhibitor Kip-related protein 1 (KRP1) in *as2-7D* restores sepal smoothness by aligning the growth directions of the outer epidermal cells along the longitudinal axis, while also increasing the overall stiffness of the outer epidermis. Furthermore, buckling is associated with the convergence of auxin efflux transporter protein PIN-FORMED 1 (PIN1) to generate outgrowth in the sepals at later stages, suggesting that buckling can initiate outgrowths. Our findings suggest that in addition to molecular cues influencing tissue mechanics, tissue mechanics can also modulate molecular signals, giving rise to well-defined shapes.

**One-Sentence Summary:** The *asymmetric leaves 2-7D* mutant sepals buckle due to discoordination of growth between the two epidermal layers.

## INTRODUCTION

During development, many organs such as the brain, intestine, and orchid flower petals form complex shapes with undulations, folds, and loops ^1–6^; conversely, other organs such as Arabidopsis leaf blades and fly wings develop smooth forms ^7–10^. These undulations, folds, and loops generally form through mechanical buckling, which is the bending of tissue that occurs when its growth patterns generate compressive forces ^11–14^. How cells within tissues regulate their growth and division to generate buckling in certain instances and maintain smoothness in others is not well understood.

Buckling plays a significant role in organ morphogenesis and development across organisms. In animals for instance, numerous studies indicate the role of buckling in organ morphogenesis ^2–4,15–17^. Differential growth of connected tissue layers gives rise to buckling. Buckling creates complex organ shapes starting from simple geometries, which is important for functional specialization. These include gyrification of the cerebral cortex to increase neuron content ^1^, formation of gut villi and looping to increase contact with food and lengthen the gastrointestinal tract ^2–4^, and initiation of new branches in lung airway epithelium to enhance gas exchange and oxygen transfer ^16^. Buckling also serves key roles in plant development. For instance, it generates helical twists in roots to enhance penetrative forces at the growing tip ^18,19^, triggers lobe deformation in Venus flytraps for rapid energy storage and release to capture prey ^20,21^, and induces nanoscale cuticular wrinkles on petals to produce iridescence ^5,6,22^. Buckling has also been proposed to drive formation and positioning of primordia on the shoot apical meristem ^23–25^.

Buckling, however, is not always beneficial. Weight of the inflorescence and external forces such as wind or rain can compress and buckle the stems of cereal crops, which causes damage and reduced yield ^26^. Similarly, slender tree stems are also vulnerable to buckling ^27^. To counter this, plants need to allocate resources to increase stem thickness and stiffness ^28,29^. While buckling generates diverse leaf shapes ^30–32^, the majority of the leaves suppress buckling, generating smooth leaf blades with a relatively flat structure that enhance light absorption and gas exchange for efficient photosynthesis ^33,34^. Similarly, sepals, which are the outermost leaf-like floral organs, also exhibit smooth surfaces, which is important for all four sepals to properly enclose and protect the developing flower bud ^35^. However, what growth mechanisms and tissue properties underlie the development of smooth organ morphology remain relatively unknown.

Using *Arabidopsis thaliana* (henceforth Arabidopsis) sepals as the model organ, we identified and characterized a gain-of-function mutant of the transcription factor *ASYMMETRIC LEAVES 2* (*AS2*), *as2-7D*, which exhibits folds and outgrowths on the outer epidermis of the sepal. We used *as2-7D* as a tool to investigate what cell growth patterns and mechanical properties of the tissue facilitate the formation of smooth sepals in wild type.

## RESULTS

### Ectopic expression of *ASYMMETRIC LEAVES 2* (*AS2*) disrupts sepal smoothness

To determine how the sepals achieve smooth morphology, we performed a forward genetic screen ^36^ and identified an Arabidopsis mutant that disrupts sepal smoothness. This mutant exhibits an irregular morphology with folds and outgrowths (Fig. 1, A to F). The folds comprise ridges and invaginations, whereas the outgrowths, which are pointed epidermal projections, are typically located at the end of the folds (Fig. 1F and fig. S1, A and B). These folds and outgrowths are visible only on the outer (abaxial) epidermis, while the inner (adaxial) epidermis is smooth like the wild type (Fig. 1, E to H, fig. S1, A and B and movie S1). Positional cloning identified a G-to-A point mutation 1,484 bps upstream of the translational start site of *ASYMMETRIC LEAVES 2* (*AS2*) (Fig. 1I). *AS2* encodes a LATERAL ORGAN BOUNDARIES (LOB) domain transcription factor that plays a critical role in leaf development, particularly in specifying the upper (adaxial) layer of the leaf ^37–40^. An identical mutation in the same locus was identified previously ^41^. However, the fold and outgrowth phenotype in the sepals was not reported ^41^, since the phenotype is very subtle in the Columbia (Col-0) background compared to what we observe in the Landsberg *erecta* (L*er*-0) background (hereafter referred to as wild type) (Fig. 1, A to D and fig. S1, C to F). Previously, Wu et. al. ^41^ demonstrated that this mutation in the *AS2* promoter disrupts the binding site of the repressor KANADI, a transcription factor involved in specifying cell fate on the lower (abaxial) side of the leaf ^42,43^. They further showed that this mutation leads to ectopic *AS2* expression on the abaxial side, while normally in the wild type, the *AS2* expression domain is restricted to the adaxial side (Fig.1I). To verify that this mutation also leads to ectopic *AS2* expression in sepals of the L*er*-0 background, we expressed GFP under the wild-type *AS2* promoter (*pAS2^WT^::GUS-GFP*) and mutant *as2-7D AS2* promoter (*pAS2^as2–7D^::GUS-GFP*), in the wild-type (L*er*-0) background. While the *AS2* promoter from wild type drives GFP specifically on the inner sepal epidermal layer (Fig. 1, J and K), the mutant *AS2* promoter drives GFP on both the inner and outer epidermal layers (Fig. 1L). The mutant *AS2* promoter drove GFP expression evenly throughout the outer epidermis, and no instance of patchy *AS2* expression patterns were observed on the outer epidermis. In addition, *AS2* is still expressed in its regular expression domain on the inner epidermis in *as2-7D* mutants (Fig. 1L). This analysis confirms that the mutation causes a gain-of-function through ectopic *AS2* expression on the outer epidermis, consistent with the dominant nature of this allele (see Materials and methods). Therefore, following the convention of previous reports ^41,44^, we named the mutant allele *as2-7D*.

**Fig. 1.**
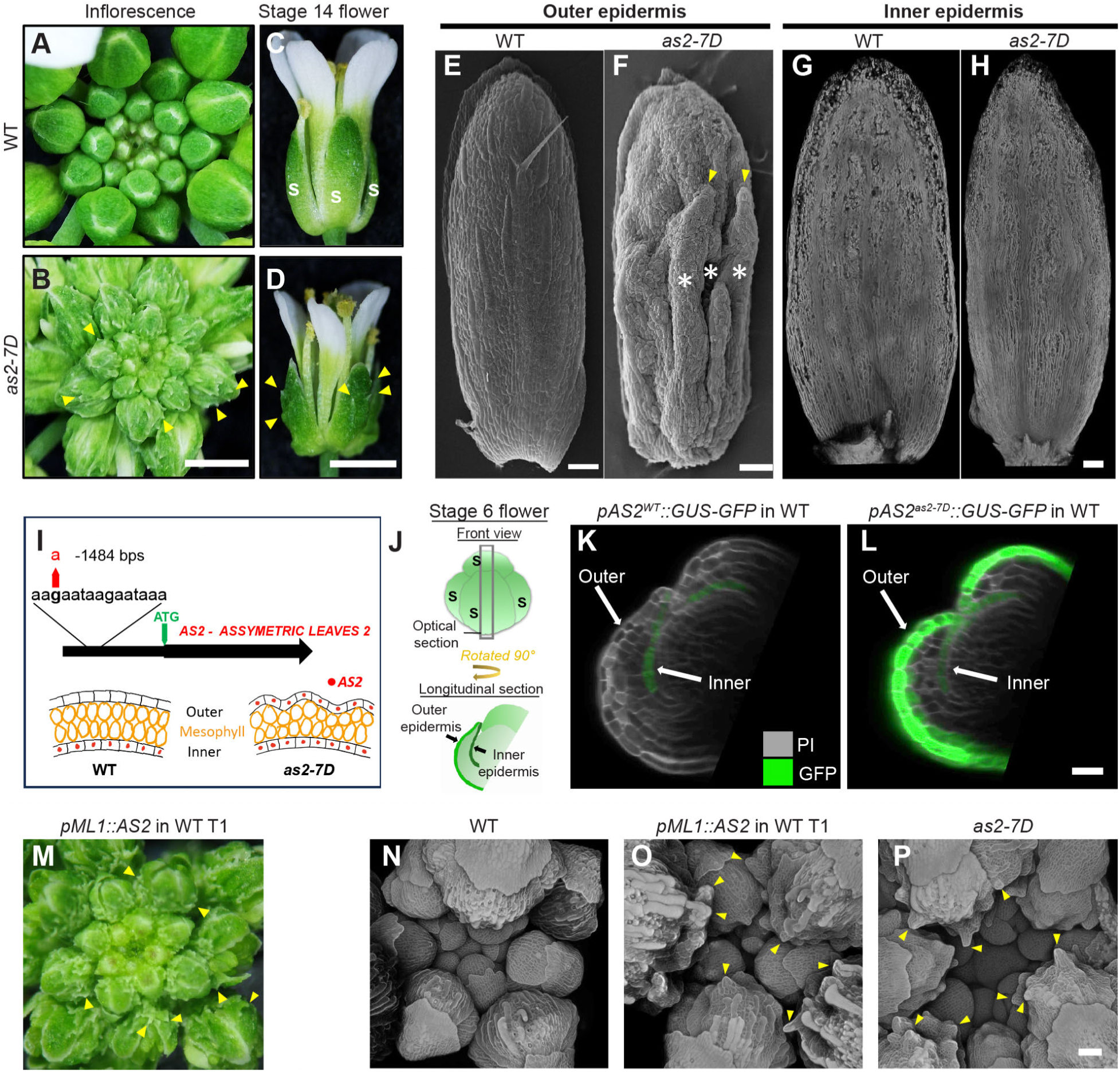
The ectopic expression of *ASYMMETRIC LEAVES 2* (*AS2*) on the outer epidermal layer disrupts sepal smoothness in *as2-7D* mutant. (**A** and **B**) Whole inflorescences of (**A**) Wild type (WT, L*er*-0) and **B**) *as2-7D* mutant (in L*er*-0 background). Yellow arrowheads indicate outgrowth. Scalebar, 1 cm; representative images of *n =* 3 inflorescences for both genotypes. (**C** and **D**) Mature flowers (stage 14 ^88^) of (**C**) WT and (**D**) *as2-7D* mutant. WT sepals (green leaf-like organs) are marked with the letter “S” in white (C). Flowers are enclosed within four sepals. Note the outgrowths (yellow arrowheads) on the outer epidermis of *as2-7D* sepals (D). Scale bar, 1 cm; representative images of *n =* 3 flowers. (**E** and **F**) Scanning electron microscopy (SEM) images of the outer epidermis of (**E**) WT and (**F**) *as2-7D* mutant sepals. Note the folds (*) and outgrowths (yellow arrowheads) on the *as2-7D* outer epidermis. Scale bar, 100 µm; representative images of *n =* 5 for WT and *n =* 10 for *as2-7D*. (**G** and **H**) Confocal microscopy images of the inner epidermis of (**G**) WT and (**H**) *as2-7D* mutant. The images are representative of T2s harboring pLH13 (*35S::mCitrine-RCI2A*), a yellow fluorescent protein-based cell membrane marker ^89^. Scale bar, 100 µm; representative images of *n =* 3 for both genotypes. (**I**) Schematic diagram representing the genetic mutation in *as2-7D*. Gene body of *AS2* showing the G-to-A point mutation (red arrow) at position 1484 bps upstream of ATG in *as2-7D* (Top half). The *AS2* expression (red dots) domain in the WT and *as2-7D* sepals is shown (Bottom half). (**J**) Anatomical diagram representing (Top) an early stage 6 flower and (Bottom) an optical longitudinal section through the flower. The letter “S” in black represents sepals. The outer and inner epidermal layers of the outer sepal are marked. (**K** and **L**) Longitudinal section of stage 6 flowers of WT. Cell walls are stained with propidium-iodide (PI, greyscale). Images show the GFP expression pattern (green) driven by (**K**) WT *AS2* promoter (*pAS2^WT^*) and (**L**) *AS2* promoter from *as2-7D* (*pAS2^as2–7D^*). Arrows indicate the outer and inner epidermal layers of the outer sepal. Scale bar = 20 µm; *n =* 3. (**M**) Whole inflorescence of a representative transgenic (T1) harboring *pML1::AS2* (*AS2* expression driven by *ATML1* epidermal specific promoter) transgene in WT. Yellow arrows mark the outgrowths on the sepal epidermis. Independent T1s assessed = 28, all of which show similar levels of outgrowth formation. Representative image of *n =* 3 inflorescences. (**N** to **P**) Confocal images of PI-stained (greyscale) inflorescences of (**N**) WT, (**O**) representative transgenic (T1) harboring *pML1::AS2* transgene in WT, and (**P**) *as2-7D* mutant. WT outer epidermis is smooth, whereas *pML1::AS2* exhibits outgrowths (yellow arrowheads) like *as2-7D*. Scale bar, 50 µm; *n =* 3 inflorescences per genotype.

To test whether ectopic *AS2* expression on the outer epidermal layer is sufficient to generate the folds and outgrowths, we drove *AS2* expression under epidermal specific *MERISTEM LAYER 1* (*ML1*) promoter (*pML1::AS2*), which ectopically expresses *AS2* on both inner and outer epidermis of the wild type. All the transgenic plants form lumps and outgrowths on the outer epidermis of the sepals, similar to *as2-7D*, which suggests that ectopic outer epidermal expression of *AS2* is sufficient to replicate the *as2-7D* phenotype (Fig. 1, M to P). Additionally, we do observe folds and outgrowths form on the epidermis of the leaves and fruits (fig. S1, G to N), consistent with previous reports ^39^, indicating a potential stronger effect of constitutive epidermal *AS2* expression in the transgenics than in the *as2-7D* gain-of-function mutant. In summary, the mutation in *as2-7D* induces ectopic *AS2* expression on the outer epidermal layer, which is sufficient to disrupt smooth sepal morphology by generating folds and outgrowths on the outer epidermal layer.

### The *as2-7D* outer epidermis appears to buckle first and then form outgrowths

To understand how ectopic *AS2* expression on the outer epidermis disrupts smooth sepal morphology, we live-imaged wild type and *as2-7D* flowers from early sepal developmental stages (Fig. 2, A and B, fig. S2, A to D, and movie S2). Initially, the outer epidermis of both the wild type and *as2-7D* sepals are smooth (Fig. 2, A and B; 0-24h). While the outer epidermis of wild-type sepals remains smooth throughout live imaging, *as2-7D* starts to develop a fold in the middle of the sepal blade at 48 hours (Fig. 2, B and C). As the flower develops, the folds intensify further, forming several inward invaginations and outward ridges throughout the outer epidermis of *as2-7D* mutant (Fig. 2, B to D and movie S2). As the folds intensify over time, we begin to see the development of phenotypically distinct pointed outgrowths. The outgrowths become apparent only in the later stages as the folds intensify, which suggests that formation of the folds precedes the emergence of the outgrowths (Fig. 2, B to D and fig. S2, C and D).

**Fig. 2.**
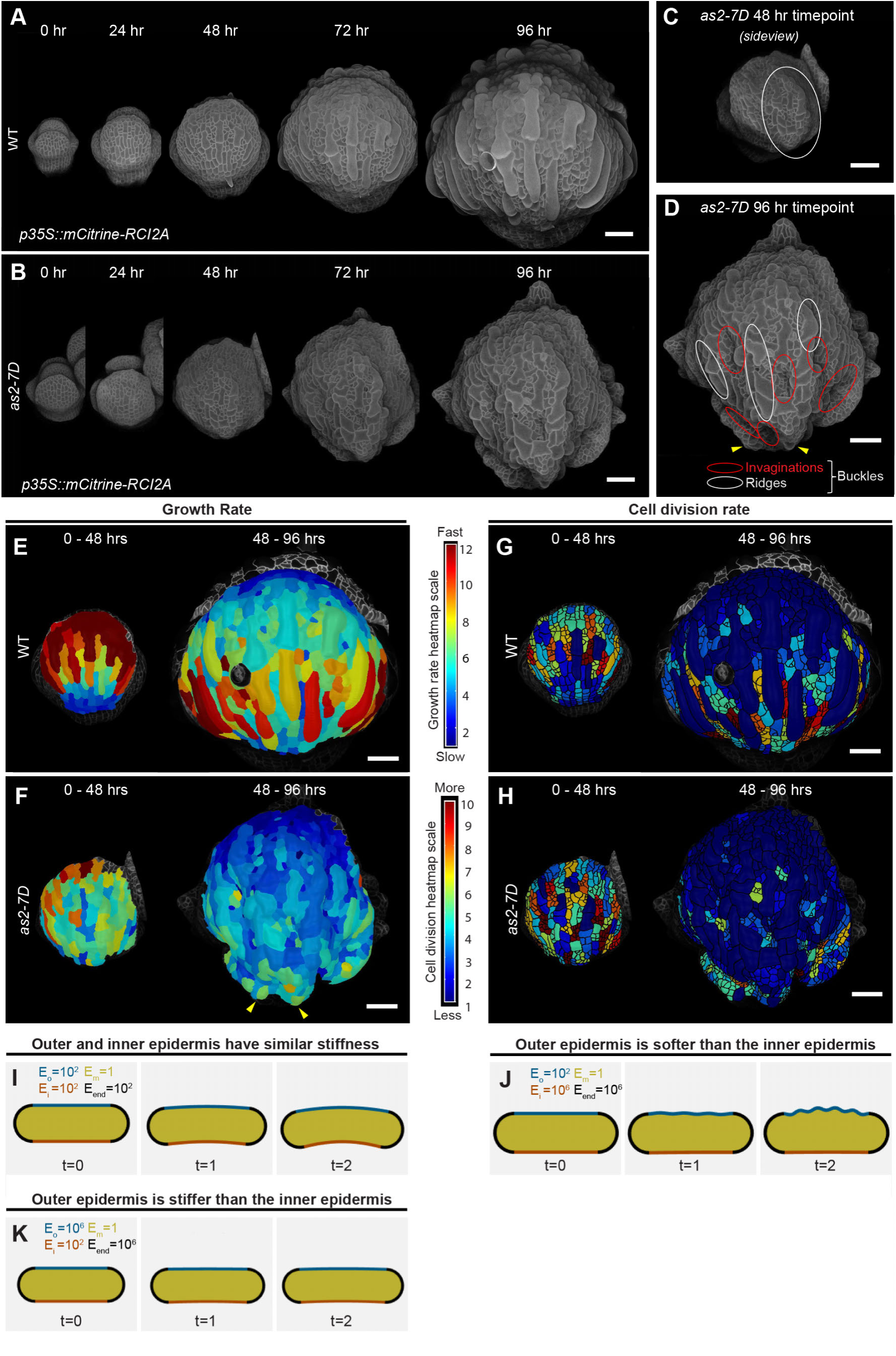
The *as2-7D* sepal outer epidermal morphology transitions from smooth to buckles and subsequently outgrowths. (**A** and **B**) Representative live-imaging series of (**A**) a wild-type (WT) flower and (**B**) an *as2-7D* mutant flower starting from developmental stage 4 over a course of 96 hours, imaged at 24-hour intervals (*n =* 3). Images capture morphological changes of the outer sepal. Both genotypes harbor the pLH13 (*35S::mCitrine-RCI2A*) cell membrane marker (greyscale). Trichomes were removed from the outer sepal epidermis by image processing to better visualization the sepal morphology. Scale bar, 50 µm. (**C**) Tilted view of *as2-7D* mutant flower at the 48-hour time point. Note the appearance of the out-of-plane protrusion (ridge) in the middle of the sepal, circled in white. Scale bar, 50 µm. (**D**) The *as2-7D* mutant flower at the 96-hour time point. Note the dramatic increase in the number and extent of ridges (red circles) and invaginations (white circles), characteristic of buckles, on the outer sepal epidermis. Yellow arrowheads indicate outgrowths. Scale bar, 50 µm. (**E** and **F**) Cell area growth rate over a 48-hour period for the (**E**) WT and (**F**) *as2-7D* mutant sepals. Growth rate is calculated as the ratio of cell lineage area at the later time point divided by that of the earlier time point. The heatmap is projected onto the later time point. Red to blue gradient shows fast to slow growth. The growth rate over 0-48 hours (left) and 48-96 hours (right) is shown. Arrowheads point to the outgrowths. Scale bar, 50 µm. (**G** and **H**) Cell division rate over a 48-hour period for the (**G**) WT and (**H**) *as2-7D*. Cell division rate is calculated based on the number of daughter cells per lineage, represented as a heatmap projected onto the later time point. Red to blue gradient shows more to less division. The cell lineages were tracked using MorphoGraphX software ^90^. Segmented cells are outlined in black. Representative heatmaps for one of the three replicates (*n =* 3) for WT and *as2-7D* are shown. Scale bar, 50 µm. (**I** to **K**) Simulations based on the analytical model (see Materials and methods), providing visual illustrations of the conditions that may favor buckling of the outer epidermis. The simulations illustrate change in tissue morphology along the cross-section when the outer epidermis was allowed to grow in width over two time points. The elastic modulus (E) of the outer epidermis (E_o_), inner epidermis (E_i_), mesophyll (E_m_) and edges (E_end_) are shown. Initial growth tensor along the width for all the four components were set to 1, with the outer epidermal growth rate set to 0.1, while the growth rates of the other components was set to 0. (**I**) Simulation outcomes when the outer and inner epidermis have similar stiffness (E_o_ and E_i_ = 10^2^), causing the tissue to curve. (**J**) When the stiffness of the outer epidermis (E_o_ = 10^2^) is less than that of the inner epidermis (E_o_ = 10^6^), the outer epidermis buckles, while the inner epidermis remains unaffected. (**K**) Simulation outcomes wherein the stiffness of the outer epidermis (E_o_ = 10^6^) is more than that of the inner epidermis (E_o_ = 10^2^), leading to a slight increase in tissue curvature over time.

We considered two hypotheses for how the folds form: (1) due to localized growth at each ridge or (2) due to mechanical buckling caused by a surface-wide growth disruption. To test the first hypothesis, we compared the cell growth rates and division rates between wild type and *as2-7D* mutant on the outer sepal epidermis at 48-hour interval over a 96-hour growth period (Fig. 2, E to H and fig. S2, E to N). We do not see any localized fast growth or cell division in the region where the first fold is formed in *as2-7D* at 48 hours (Fig. 2, C to F). In the initial stages (0-48 hours), the wild type exhibits the stereotypical basipetal growth pattern described earlier ^45^, exhibiting fast cell growth at the tip, moderate growth in the center, and slow growth towards the base. In contrast, the cells grow at relatively homogeneous rates throughout the outer epidermis of *as2-7D* (Fig. 2, E and F; 0-48 hours). Although the pattern of growth is different, we observe no difference in the average cell growth rates between the wild type and *as2-7D* (fig. S2M). This pattern is also reflected in cell division rates, where we observe a basipetal pattern in wild type and a more uniform pattern throughout the sepal epidermis in *as2-7D* (Fig. 2, G and H; 0-48 hours). Our results suggest that the folds form due to surface-wide disruption of cell growth patterns, and not because of localized fast cell growth and division at the location of the ridge. In the later time point (48-96 hrs) the growth is still relatively uniform along the places where the *as2-7D* folds form. In contrast to folds, we do observe localized growth and division later at sites where outgrowths form (Fig. 2F and fig. S2, E to L; 48-96 hrs). We therefore considered the hypothesis that the folds on *as2-7D* outer epidermis may form through mechanical buckling caused by surface-wide growth defects.

### Modeling predicts two criteria required for buckling

Based on our observations of *as2-7D* development, we formulated a simple two-dimensional (2D) analytical model to determine the basic conditions that could give rise to buckling (see Materials and methods). Our model indicates that buckling can occur if the following two conditions are simultaneously fulfilled. Firstly, if the average specified growth rate (the cell-autonomous growth rate achieved if the layers were isolated) of the outer and inner epidermis is higher than that of the mesophyll, then the epidermal layers will be under compression, which will favor buckling in the epidermal layers. Secondly, considering that the mesophyll is known to be softer than the epidermis ^46^, a significantly lower stiffness of the outer epidermis compared to the inner epidermis can cause the outer epidermis specifically to buckle, without causing any buckling of the inner epidermis.

We further tested the validity of the analytical model with simulations on a simple geometry: a cross-sectional representation of a growing elastic system, comprising of the outer epidermis, the mesophyll, the inner epidermis, and the edges (Fig. 2, I to K, fig. S3 and S4). In these simulations, only the outer epidermis was allowed to grow in width, which is equivalent to the outer epidermis having a higher specified growth rate than the other two layers. Under these conditions, when the outer and inner epidermis exhibit similar stiffness, the tissue curves while maintaining a smooth morphology for both the outer and inner epidermis (Fig. 2I). However, when the outer epidermis is softer than the inner epidermis, the outer epidermis buckles, while the inner epidermis shows no change in morphology (Fig. 2J). In contrast, increasing the stiffness of the outer epidermis relative to the inner epidermis restores the smooth morphology of the outer epidermis, while the inner epidermal morphology remains unaffected (Fig. 2K). In summary, although our simple analytical model and simulations do not account for all the complex dynamics of tissue growth in 3D. However, our model does help guide our investigations towards testing whether differences in growth and tissue stiffness properties between the outer and inner epidermal layers contribute to buckling in *as2-7D*.

### Aligned of cell growth orientation between epidermal layers is essential for the formation of smooth sepals

To test the first of our model predictions, we live-imaged the wild type and *as2-7D* sepals through the stages prior to buckling (Fig. 3, A and B), and extracted the inner and outer epidermal layers of the same growing sepals by image processing (Fig. 3, C and D). We found that cells throughout the wild-type inner epidermis grow significantly faster than the outer epidermal cells (Fig. 3E and fig. S5, A, B, and E). We also observe fast cell divisions on the inner epidermis (Fig. 3G and fig. S5, F, G, and J). The fast growth of the inner epidermis was surprising, since for the sepals to eventually grow into flat shape with smooth morphology, one would intuitively expect the cells on both epidermal layers to grow at similar rates. However, during the initial developmental stages, the sepals are highly curved along the longitudinal direction, due to the smaller surface area of the inner compared to the outer epidermis. Hence, to transition from an initially curved shape to a subsequently flat structure, the inner epidermis of the sepal grows faster compared to the outer epidermis. (Fig. 3C and D, Fig. 4, and movie S2). Like wild type, the inner epidermal cells of *as2-7D* mutant also grow faster than outer epidermal cells; however, the inner epidermal cells of *as2-7D* do not grow as fast as wild type (Fig. 3, E and F and fig. S5, A to E). In concert with the growth rate, the cell division rate is also lower on the inner epidermis of *as2-7D* compared to wild type (Fig. 3, G and H and fig. S5, F to J).

**Fig. 3.**
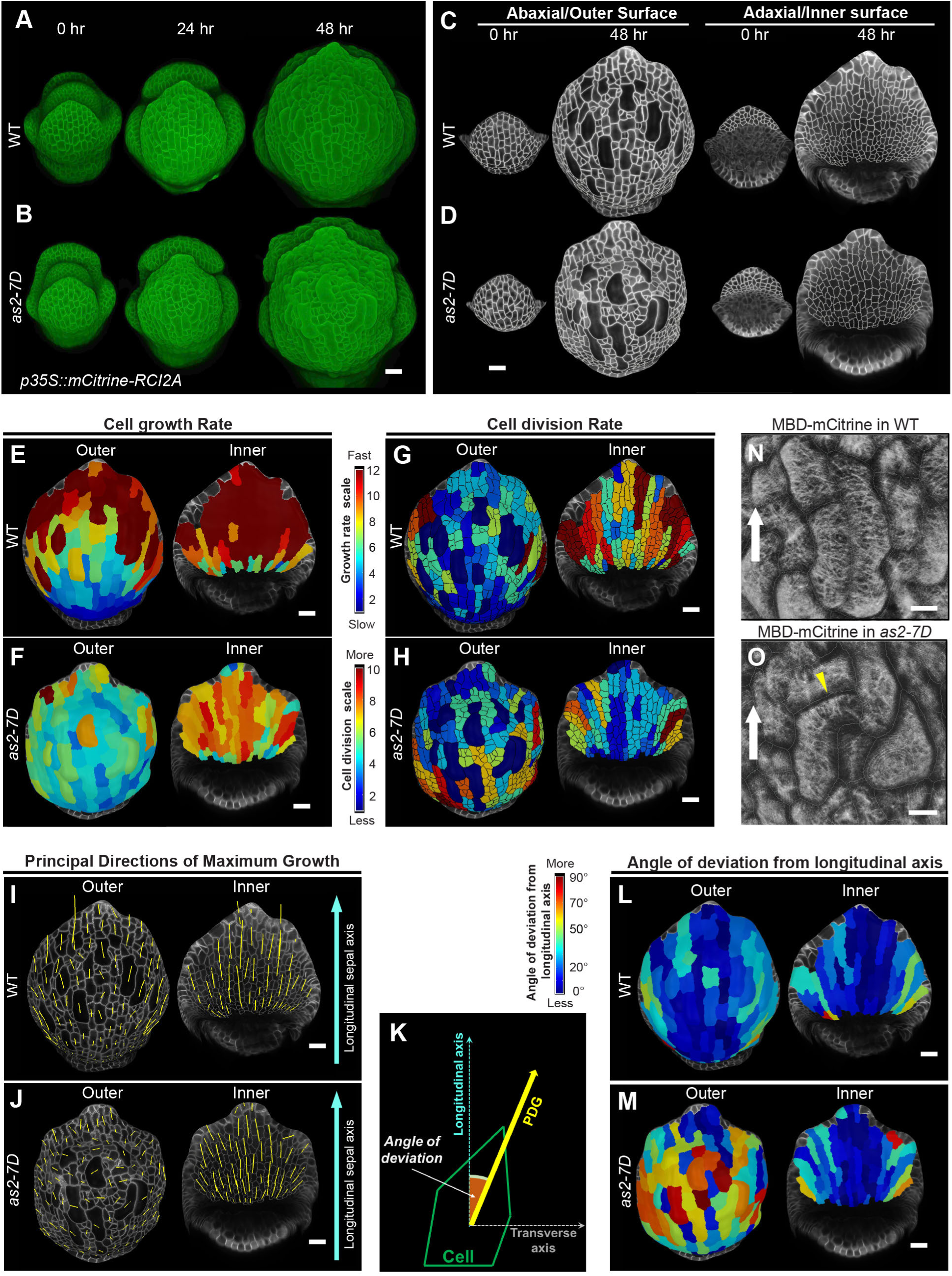
Unidirectional growth on both epidermal layers is essential for smooth sepal development. (**A** and **B**) Representative confocal images of the (**A**) Wild type (WT) and (**B**) *as2-7D* mutant live-imaged with a high laser intensity (∼16-20%) every 24 hours. Both the genotypes harbor pLH13 (*35S::mCitrine-RCI2A*) cell membrane marker (green). Scale bar, 20 µm. One of the three representative replicates (*n =* 3) is shown. (**C** and **D**) Segmented cells in grey superimposed onto the plasma membrane signal (also in grey) projected on mesh surfaces generated using MorphoGraphX ^90^. Mesh surfaces for 0 and 48-hour time points of the outer (left) and inner (right) epidermis of the (**C**) WT and (**D**) *as2-7D* mutant are shown. Note the strikingly reduced number of cells on the inner versus outer surface at the 0-hour time point. Scale bar, 20 µm; *n =* 3. (**E** and **F**) Cell growth rates over a 48-hour period for the (**E**) WT and (**F**) *as2-7D* mutants are shown. The heatmap has been projected onto the 48-hour time-point surface. In each panel, the growth rate of cells on the outer surface (left) and inner surface (right) is shown. Red to blue heatmap gradient shows fast to slow growth. Scale bar, 20 µm; *n =* 3. (**G** and **H**) Cell division rates over a 48-hour period for the (**G**) WT and (**H**) *as2-7D* mutant on the outer surface (left) and inner surface (right) are shown. Segmented cells are outlined in black. Red to blue gradient shows more to less/no cell division respectively. Scale bar, 20 µm; *n =* 3. (**I** and **J**) Principal directions of maximal growth (PDG_max_, yellow) of the cells on the outer (left) and inner (right) epidermis of (**K**) WT and (**L**) *as2-7D*. Segmented cells in grey are superimposed on the plasma membrane signal (also in grey). Length of any given PDG_max_ tensor is directly proportional to the cell growth rate in that direction. A blue arrow indicates the longitudinal sepal axis. Note the misoriented PDG_max_ tensors on the *as2-7D* outer epidermis. Scale bar, 20 µm; *n =* 3. (**K**) A schematic diagram depicting the calculation of the angle of deviation between PDG_max_ tensor and the longitudinal axis to obtain a quantitative measure of the deviation. (**L** and **M**) Heatmaps show the angle of deviation of the PDG_max_ tensor from the sepal longitudinal axis for the outer (left) and inner (right) epidermal cells of the (**L**) WT and (**M**) *as2-7D* mutant. Scale for the angle of deviation ranges from 90° (highly deviated, red) to 0° (no deviation, blue). Scale bar, 20 µm; *n =* 3. (**N** and **O**) Cortical microtubule orientation (greyscale) from comparable regions of stage 6 flower outer epidermis for the (**I**) WT and (**J**) *as2-7D* mutant is shown. These plants harbor *pPDF1::mCitrine-MBD*, a YFP based microtubule marker ^91^. . Cells segmented using meshes generated from PI-stained images are outlined in gray. White arrows point towards the longitudinal axis of the sepals. Microtubules in the WT are oriented perpendicular to the sepal longitudinal axis, whereas, in *as2-7D*, the microtubules are misoriented. Yellow arrowheads point towards the misoriented microtubules. Scale bar, 10 µm; *n =* 3.

**Fig. 4.**
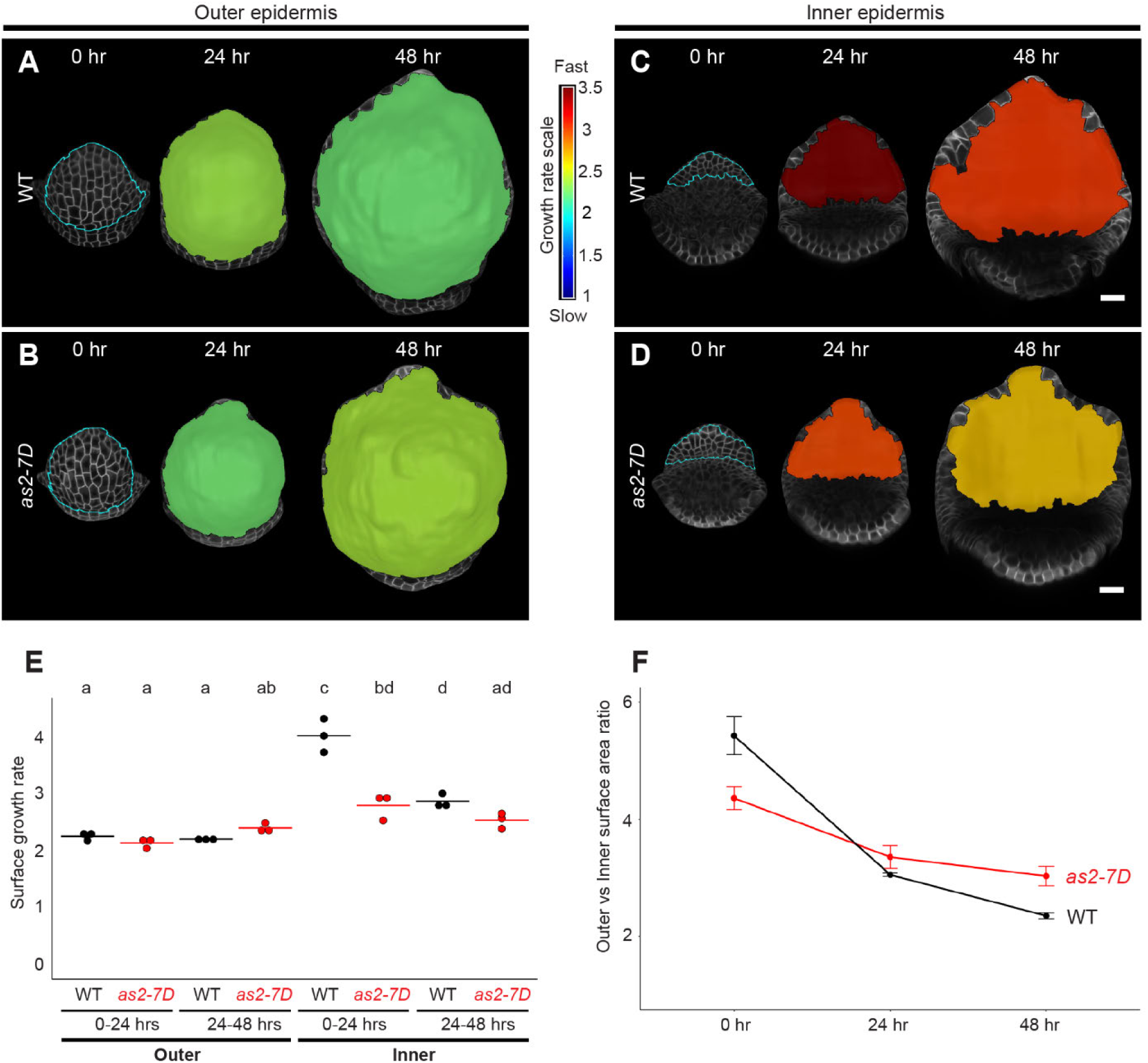
Slower growth on the inner epidermis leads to overgrowth of the outer epidermis in *as2-7D*. (**A** to **D**) Heatmaps represent growth rates of (**A**) Wild-type (WT) outer, (**B**) *as2-7D* outer, (**C**) WT inner, and (**D**) *as2-7D* inner epidermal surface at every 24-hour interval, over a 48-hour live imaging period. Growth rate is projected onto the later time point. Growth rate for the region demarcated in cyan at 0-hour time point was tracked over time. Heatmap: red, fast growth, and blue, slow growth. Representative heatmaps for *n =* 3 biological replicates are shown. Scalebar, 20 µm. (**E**) Graph shows epidermal surface growth rates every 24 hours for the WT (black) and *as2-7D* (red) is shown (*n =* 3). Lowercase letters on top of the data points represent significant differences based on one-way ANOVA with Tukey’s HSD post-hoc test (*p* < 0.05, similar letters represent no significant difference).

**Fig. 5.**
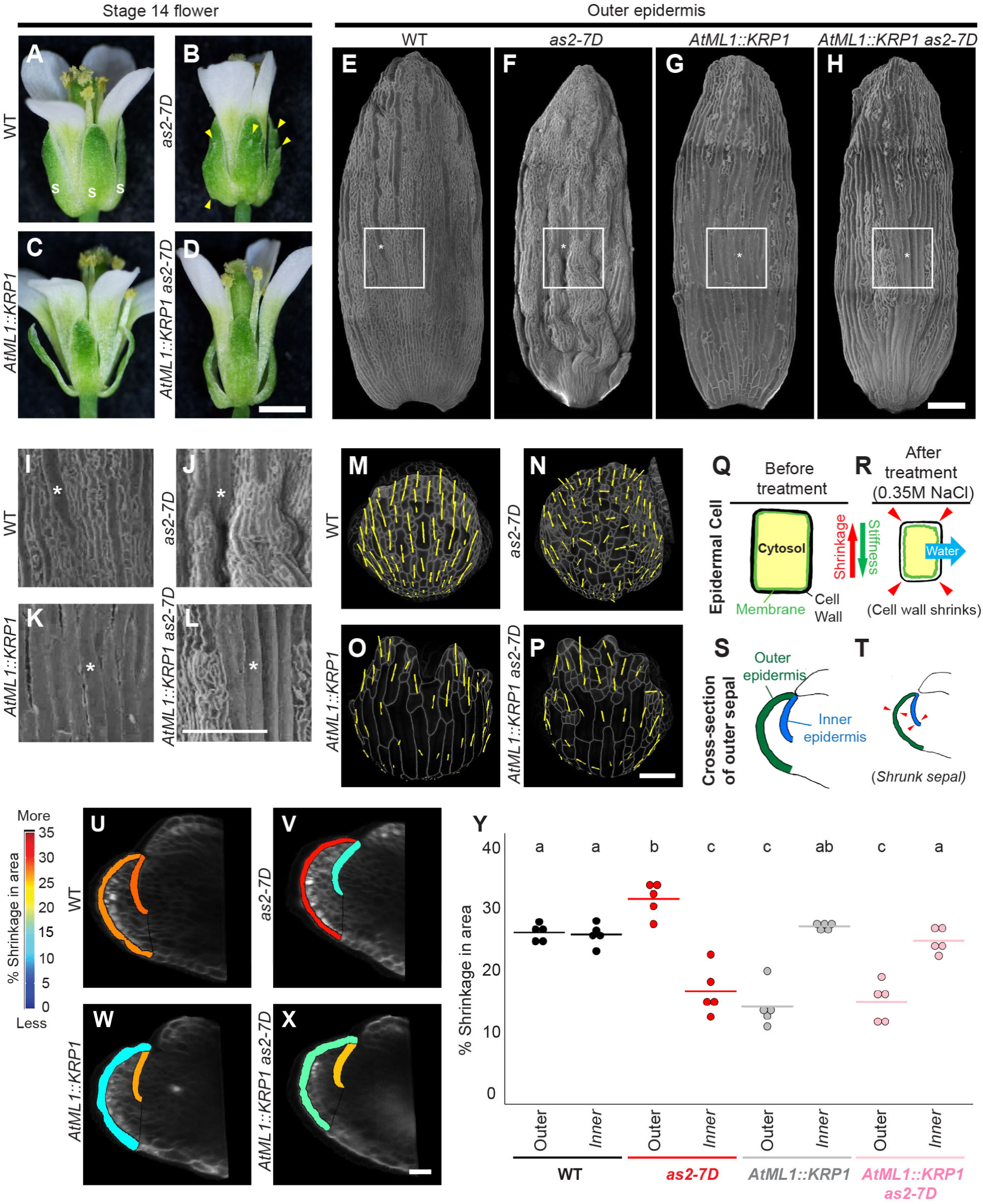
Aligning growth along the longitudinal direction and increasing outer epidermal stiffness is sufficient to rescue buckling in *as2-7D*. (**A** to **D**) Representative stage 14 flowers of (**A**) Wild type (WT), (**B**) *as2-7D* mutant, (**C**) *AtML1::KRP1,* and (**D**) *AtML1::KRP1 as2-7D*. The WT sepals are marked with the letter “S” in white (A). Yellow arrowheads point to outgrowths in *as2-7D* (B). Note the smooth sepal of *AtML1::KRP1 as2-7D* compared to *as2-7D*. Scale bar, 1 cm; *n =* 3 flowers for all genotypes. (**E** to **H**) Confocal images of the outer epidermis of (**E**) WT, (**F**) *as2-7D* mutant, (**G**) *AtML1::KRP1*, and **(H)** *AtML1::KRP1 as2-7D* stage 14 sepals. The sepals were stained with PI (grey, cell walls). Note the increased number of highly elongated giant cells in *AtML1::KRP1* and *AtML1::KRP1 as2-7D* sepal, and the smooth outer epidermis of *AtML1::KRP1 as2-7D* sepal compared to the highly rugose morphology of *as2-7D*. Scale bar, 200 µm; *n =* 3. (**I** to **L**) Magnified insets of the (**I**) WT, (**J**) *as2-7D* mutant, (**K**) *AtML1::KRP1,* and (**L**) *AtML1::KRP1 as2-7D* sepal for the regions marked by white boxes in panels E-H respectively. Asterisks denote a typical giant cell. Scale bar, 200 µm; *n =* 3. (**M** to **P**) The principal direction of growth (PDG_max_) for the outer epidermal cells of the **(M)** WT, **(N)** *as2-7D,* (**O**) *AtML1::KRP1,* and (**P**) *AtML1::KRP1 as2-7D* sepals. Yellow lines mark the PDG_max_ tensors calculated over a 48-hour period, projected onto the later time point. For WT and *as2-7D*, the flower images used are the same as in Figure 2. Segmented cells in grey are superimposed onto the plasma membrane signal (also in grey). Scale bar, 20 µm; *n =* 3. (**Q** and **R**) Schematic diagrams demonstrating the concept of cell wall stiffness assessment by osmotic treatment assays ^55^. (**Q**) A typical plant epidermal cell before osmotic treatment. The cell wall and cell membrane are shown in black and green respectively. (**R**) A cell that has undergone plasmolysis in a hyperosmotic solution (0.35 M NaCl). (**S** and **T**) Anatomical diagrams demonstrate the use of osmotic treatments to estimate epidermal stiffness. Longitudinal sections of an early stage 6 flower **(S)** before and (**T**) after osmotic treatment are shown. The outer (green) and inner (blue) sepal epidermis are marked. Arrowheads (red) demonstrate aerial shrinkage. (**U** to **X**) Heatmaps show the aerial shrinkage (in percentage) of a longitudinal sepal cross-section for the inner and outer epidermis of (**U**) WT, (**V**) *as2-7D* mutant, (**W**) *AtML1::KRP1* and (**X**) *AtML1::KRP1 as2-7D*, post osmotic treatment in 0.35M NaCl solution. Shrinkage was computed as the decrease in area of a comparable cross-sectional region from an early stage 6 flower before and after osmotic treatment. Heatmaps are projected on the post-treatment surface. More shrinkage is represented in red and less in blue. Scale bar, 20 µm. Heatmaps for each genotype are representative of 5 independent replicates (*n =* 5). (**Y**) Graph showing the percentage aerial shrinkage post osmotic treatment. Data across 5 independent replicates each for the WT (black dots), *as2-7D* (red dots), *AtML1::KRP1* (grey dots), and *AtML1::KRP1 as2-7D* (pink dots) for both the outer and inner epidermis are shown. The letters above the plots represent significant differences based on one-way analysis of variance (ANOVA) with Tukey’s honest significant difference (HSD) post hoc correction (*p* < 0.05). Similar letters indicate no significant difference.

Since buckling is a surface-wide phenomenon in our model, we analyzed the differences in whole inner versus outer epidermal growth rates. On an epidermis-wide scale, we found that the outer epidermis of wild type and *as2-7D* grow at the same rate (Fig. 4, A, B and E). This observation is consistent with the growth rate data of individual cell lineages, where the average outer epidermal cell growth rates of wild type and *as2-7D* show no significant difference (Fig. 4, A, B and E and fig. S5E). However, the inner epidermis of *as2-7D* grows significantly slower than wild type, which also aligns with the growth rate data of individual cell lineages on the inner epidermis (Fig. 4, C, D and E and fig. S5E). Our findings indicate that the rate at which the inner epidermis of *as2-7D* grows is insufficient to catch up with the increasing outer epidermal surface area (Fig. 4F).

We then asked whether outer epidermal buckling in *as2-7D* could potentially be attributed to local hypergrowth of the mesophyll cells underneath the outer epidermis. To test this, we processed the sepal images using Convoluted Neural Network (see Materials and methods) ^47^ to enhance the conspicuity of mesophyll cell membranes (fig. S6, A to H). We analyzed the growth rates of the outermost layer of mesophyll cells which contains fewer airspaces in early stages, thus allowing us to lineage track the cells over time. Our findings revealed that the spatial growth patterns of the mesophyll cells of both wild type and *as2-7D* mimic their respective outer epidermal cell growth patterns (fig. S6, I and J). Specifically, the wild-type mesophyll cells exhibit the same basipetal growth gradient as that of the wild-type outer epidermal cells (Fig. 3E and fig. S6I). The mesophyll cells of *as2-7D* exhibit homogeneous cell growth rates, similar to the *as2-7D* outer epidermis (Fig. 3F and fig. S6J). Comparable to the outer epidermal cell growth patterns, we did not find any significant difference in the average growth rate of the mesophyll cells between wild type and *as2-7D* (fig. S6M). Notably, there was no evidence of local hypergrowth in the central region of the sepal blade, where the initial buckling occurs (fig. S6J). To test our model predictions, we then compared the magnitude of growth of the outer and inner epidermal cells relative to the mesophyll (fig. S6N). We found that the average growth rates of the inner and outer epidermal cells significantly exceed that of the mesophyll cells for both wild type and *as2-7D* (fig. S6N), which aligns with the predictions of our model, suggesting that the epidermal layers are under compression.

Since cell growth directions are critical for determining organ shapes ^48–50^, we assessed the principal direction of maximal cell growth (PDG_max_) in the wild type and *as2-7D* mutant. The wild-type cells on both epidermal layers grow in a highly aligned manner along the sepal longitudinal axis (Fig. 3I). In *as2-7D*, although inner epidermal cells grow parallel to the sepal longitudinal axis, the outer epidermal cells exhibit highly misaligned growth, with the majority of PDG_max_ tensors oriented in the transverse direction (Fig. 3J). Similar to the growth directions of the outer epidermal cells, the mesophyll cells of wild type grow longitudinally, while the mesophyll cells of *as2-7D* exhibit misaligned transverse growth (fig. S6, K and L). We quantified the growth directions of the epidermal cells by calculating the deviation angle of PDG_max_ tensors relative to the sepal’s longitudinal axis (Fig. 3K). The heatmaps indicate minimal deviation in cell growth directions of the wild-type epidermal layers and the inner epidermal layer of *as2-7D* (Fig. 3, L and M and fig. S5, K to O). In contrast, the outer layer of *as2-7D* displayed a high degree of deviation in growth directions (Fig. 3M and fig. S5, K to O). Since microtubules are known to align perpendicular to the maximal cell growth direction ^51,52^, we then tested the microtubule orientation pattern in comparable regions of the wild type and *as2-7D* outer epidermis (Fig. 3, N and O). For the wild type, the microtubules show the typical transverse alignment pattern, which is perpendicular to the sepal longitudinal growth direction (Fig. 3N). Conversely, the microtubules are largely misoriented in the *as2-7D,* and in certain instances, mirror the transverse cell growth pattern (yellow arrowheads, Fig. 3O).

Taken together, overgrowth of the *as2-7D* outer epidermis in width compared to the inner epidermis contributes to the buckling of the outer epidermis. Our observations align with the prediction of our model simulations, which indicates that the transverse expansion of the outer epidermis, constrained along the edges by the inner epidermis would favor buckling. Overall, our data suggests that anisotropic growth in an aligned manner on both the epidermal layers is important for maintaining smooth sepal morphology.

Since we hypothesized that the transverse growth orientation of the outer epidermal cells causes the outer epidermis to buckle, we tested whether forcing the cells to grow along the longitudinal direction can rescue the *as2-7D* phenotype. Overexpressing cyclin-dependent kinase (CDK) inhibitor Kip-related protein 1 (KRP1) throughout the epidermis (*pML1::KRP1*) at moderate levels trigger the epidermal cells to skip division and undergo endoreduplication early during development, resulting in an increase in giant cell formation ^53,54^. Giant cells, which are exclusive to the outer epidermis, exhibit the unique property of growing in a highly anisotropic manner ^53^ along the longitudinal sepal axis (Fig. 5, A, C, E and G, fig. S7, A and C, and movie S3). Therefore, we wondered whether increasing giant cell formation can restore directional growth in *as2-7D* and consequently reduce buckling. Indeed, overexpressing *KRP1* in *as2-7D* (*pML1::KRP1 as2-7D*) largely suppresses buckling of the outer sepal epidermis (Fig. 5, A to H, fig. S7, A to D and movie S3). As expected, overexpressing KRP1 increases the number and size of the giant cells, which grow in a highly aligned manner along the longitudinal sepal direction (Fig. 5, E to P and movie S3). The PDG_max_ tensors of *pML1::KRP1 as2-7D* outer epidermal cells are similar to those of *pML1::KRP1*, wherein the transverse growth orientation exhibited by *as2-7D* is largely re-oriented longitudinally (Fig. 5, M to P and fig. S7, E to L). In addition to reduced buckling, the characteristic outgrowths of the *as2-7D* outer epidermis that are formed downstream of the buckles, are also nearly absent in *pML1::KRP1 as2-7D* (Fig. 5, E to H), further suggesting that buckling is required for the formation of the outgrowths. While there is no obvious association between the location of giant cells and small cells and the buckles in the *as2-7D* mutant, it is possible that the other properties caused by overexpression of *KRP1*, such as changes in stiffness (verified below), increased outer epidermal thickness, and reduction of cell division in *pML1::KRP1 as2-7D*, in addition to the growth orientation contributes to the suppression of buckling. Overall, consistent with our hypothesis, our findings suggest that transverse growth of the outer epidermal cells may be a key driver of buckling in *as2-7D*.

### Reduced stiffness of the outer epidermis compared to the inner epidermis is associated with buckling of the outer epidermis in *as2-7D* mutants

The second condition of our 2D analytical model predicts that buckling can be restricted to the outer epidermis if it is softer than the inner epidermis. To test the relative stiffness of the outer and the inner epidermal layers, we performed osmotic treatments ^55,56^ and compared how the epidermal layers of *as2-7D* shrinks relative to wild type. Briefly, when exposed to a hyperosmotic solution, cells undergo plasmolysis and no longer exert turgor pressure on the cell wall, causing the cell wall to contract (Fig. 5, Q and R). The extent of cell wall shrinkage gives a proxy measure for cell wall stiffness, with low shrinkage indicating the wall is stiff (high Young’s modulus) and high shrinkage indicating the wall is soft (low Young’s modulus) ^55^. We exposed early sepals to salt solution and measured the epidermal shrinkage (Fig. 5, S and T, fig. S7, M to P, and see Materials and methods). We found that the wild-type epidermal inner and outer layers exhibit similar shrinkage, which indicates similar stiffness across epidermal layers (Fig. 5, U and Y). In contrast, the outer epidermis of *as2-7D* shrank more than the inner epidermis, indicating that *as2-7D* outer epidermis is softer (Fig. 5, V and Y). This observation is also consistent with the second condition of our analytical model, which proposes that reduced stiffness of the outer epidermis is an essential factor for buckling (Fig. 2, I and J). We then wondered whether increasing giant cell formation affects the stiffness of the outer epidermis. To test this, we performed osmotic treatments on *ML1::KRP1* and *ML1::KRP1 as2-7D*. Indeed, we found increased stiffness of the outer epidermis in both *ML1::KRP1* and *ML1::KRP1 as2-7D* (Fig. 5, W to Y), which aligns with our simulation predictions that increasing outer epidermal stiffness may restore smooth epidermal morphology (Fig. 2K).

Taken together, our results suggest that aligning growth along the longitudinal direction, decreasing cell division, and increasing outer epidermal stiffness suppresses buckling and subsequent outgrowth formation in *as2-7D*. However, we cannot also decouple the possibility that accelerated differentiation resulting from ectopic *KRP1* expression could also potentially contribute as an additional factor to the rescue. Our findings collectively suggest that relative stiffness and growth orientations of the epidermal layers play a critical role in shaping sepal morphology. Specifically, growth along the same direction and comparable stiffness across both epidermal layers are essential criteria for smooth sepal formation, as observed in the wild type.

### Buckles in early *as2-7D* sepals are not caused by an apparent disruption of abaxial-adaxial patterning

We have focused on characterizing how aberrations in growth and mechanical properties lead to buckling on the outer epidermis of *as2-7D* sepals. As described previously, the buckles appear as folds, and these folds are phenotypically very distinct from the outgrowths (which appear as pointed projections later during sepal development). While formation of the folds and its characterization as buckles was previously unreported, the genetic basis of how the outgrowths form on aerial tissues is well established. Waites and Hudson showed nearly 30 years ago how distinct outer (abaxial/lower) and inner (adaxial/upper) tissue identities are established, and more importantly, how the juxtaposition of the outer and inner identities is essential for the lateral growth of the leaf lamina ^57^. In the leaves of *phantastica* (*phan*) mutants of *Antirrhinum majus* (snapdragon), patches of inner tissue on the outer surface generate ectopic ridges that resemble lamina margins, indicating that juxtaposition of outer and inner identities can trigger the initiation of ectopic laminar outgrowths ^57^. Since the time Waites-Hudson proposed how outgrowths are formed, several studies have independently shown that disruption in the gene regulatory network which determines outer-inner identities can generate ectopic outgrowths ^58^.

Broadly, outer identity is specified by *KANADI*, *YABBY*, and *AUXIN RESPONSE FACTOR* (*ARF*) gene families, while inner identity is established by the HD-ZIP class III genes, *ASSYMETRIC LEAVES 1* (*AS1*) and *AS2* (Fukushima and Hasebe 2014). While *kan1* and *kan2* single mutants show mild phenotypic defects, the *kan1 kan2* double mutants exhibit radialized organs with ectopic outgrowths on the outer side ^42,43^. In the *kan1 kan2 kan4* triple mutants, the ectopic outgrowths are observed even in the hypocotyl and outer surface of the cotyledons ^59^. Loss of function double mutants of YABBY genes *FILAMENTOUS FLOWER* (*FIL*) and *YABBY3* (*YAB3*), *fil yab3*, show only mild polarity defects compared to *kan1 kan2*, suggesting partial redundancy or distinct functions ^42^. In line with this, the characteristic folds and outgrowths seen in *kan1 kan2* are largely absent in the quadruple mutants *kan1 kan2 fil yab3*, although the leaves are radialized and severely reduced in size ^60^. The outgrowths in *kan1 kan2* double mutants also exhibit elevated levels of *FIL* expression, indicating that YABBYs are essential for progression of the outgrowths ^60^.

Initiation of leaf primordium occurs at the boundary between the concentric, non-overlapping expression domains of *REVOLUTA* (*REV*) (a member of the HD-ZIP class III gene family) and *KAN1* ^61^. The *WUSCHEL-RELATED HOMEOBOX* (*WOX*) genes *PRESSED FLOWER* (*PRS*)/*WOX3* and *WOX1*, which are expressed in the interface between inner and outer domains, are also required for outgrowth formation^62^. Ectopic *WOX1* expression is sufficient to induce outgrowth formation. While *prs wox1* double mutants exhibit narrow, upward-curved leaves, the ectopic outgrowths are highly suppressed in the *prs wox1 kan1 kan2* quadruple mutants, although the leaves still exhibit a folded morphology ^62^. Similarly, *prs wox1 as2* triple mutants fail to produce outgrowths and instead develop radial filamentous leaves. On the contrary, *ARF2*, *ARF3* and *ARF4*, which restrict the expression domain of *WOX1/PRS*, repress the formation of ectopic outgrowths ^63^. In particular, the *arf2 arf3 arf4* triple mutants show outgrowths on the outer surface of the leaves due to expanded *WOX1/PRS* expression ^63^. Thus, WOX activity is required for the formation of ectopic outer outgrowths.

As stated previously, *AS2* is a key transcription factor that is essential for establishing inner identity^37–40^. Fundamentally, transcription factors cannot drive shape formation without regulating tissue mechanics. Hence, *AS2* must exert its effects on tissue growth and mechanics by regulating downstream genes. It is well known that overexpression of *AS2* cause outgrowth formation on the outer surface of the leaves, and this is accompanied by repression of outer identity genes (*KAN1*, *KAN2*, *FIL*, *YAB3*) ^39^. We have shown that in addition to the outgrowths, *as2-7D* also exhibit folds on the outer sepal epidermis. While there exists substantial evidence for the genetic basis of outgrowth formation, the existence of the epidermal folds, as well as how these folds are formed in the first place was not well characterized before. Our evidence strongly supports our hypothesis that the folds observed in the outer epidermis of *as2-7D* in the initial stages are in fact buckles, formed due to disrupted growth patterns and mechanical properties. However, since *AS2* is known to promote inner epidermal identity ^37–39^, we tested the alternative hypothesis that the initial folds observed on the outer epidermis of *as2-7D* are caused by ectopic juxtaposition of inner and outer identity. Therefore, we looked at the expression pattern of *REV* as an inner identity transcription factor ^64^, and *FIL* as an outer identity transcription factor ^65^. We observe no difference in *REV* and *FIL* expression patterns between wild type and *as2-7D*, and particularly, no ectopic juxtaposition between inner and outer gene expression was formed in the *as2-7D* sepal surface (Fig. 6, A to H). Since REV promotes adaxial identity in conjunction with AS2, we tested whether REV is necessary or sufficient for the *as2-7D* phenotype. Ectopic *AS2* expression in the *rev-6* mutant (*pML1::AS2 rev-6*) leads to folds and outgrowth formation similar to *pML1::AS2* in wild type (Fig. 1, M to P, fig. S1, G to N, Fig. 6, I to L and fig. S8, A to I), indicating REV is not necessary for either the buckles or the outgrowths. Conversely, ectopic expression of *REV* in the wild-type sepals is insufficient to generate buckles or outgrowths (fig. S8, J and K). Further, ectopic *REV* expression in *as2-7D* also does not enhance the phenotype (fig. S8, L and M).

**Fig. 6.**
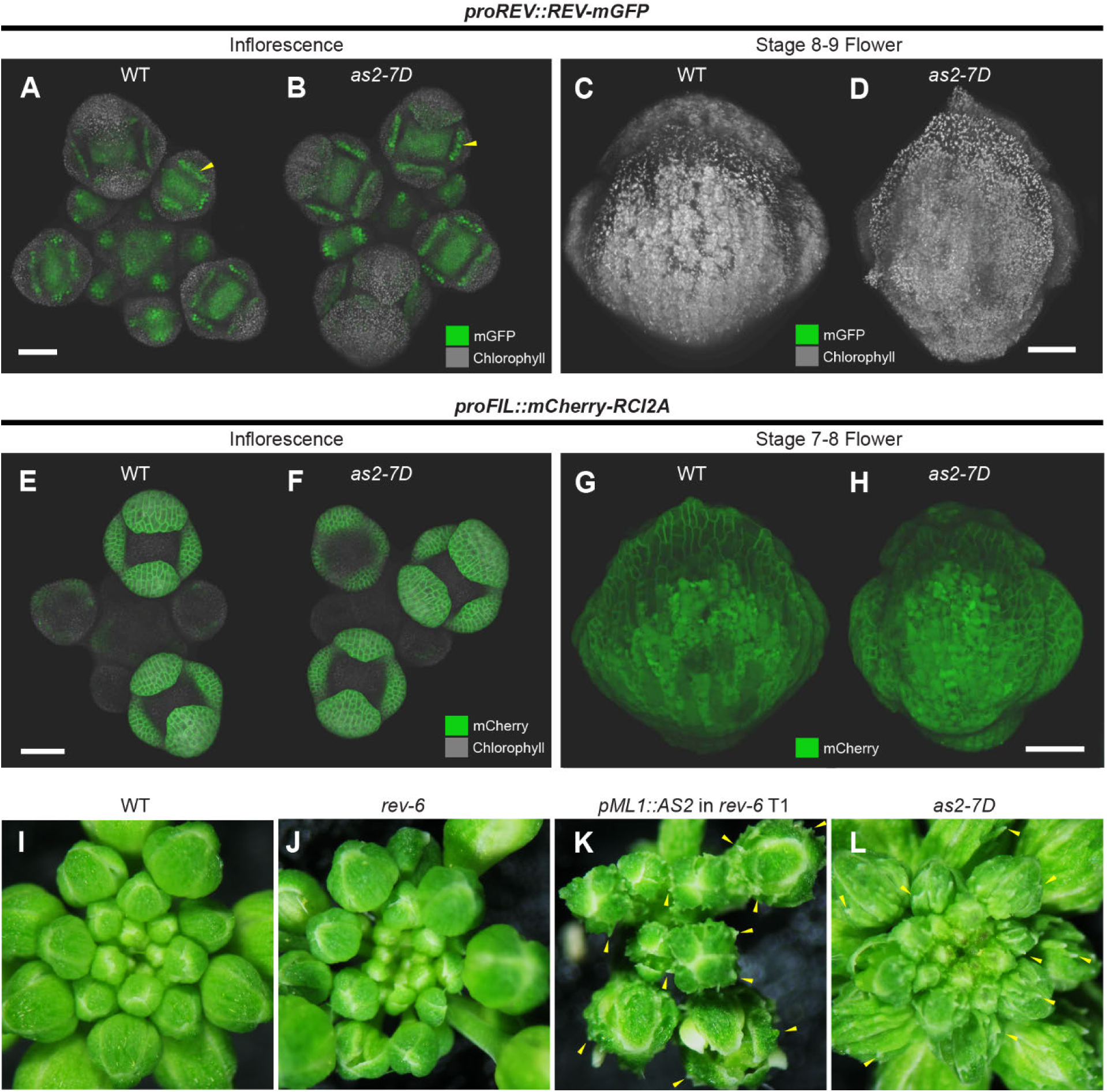
Buckling in *as2-7D* is not caused due to disruption of outer (abaxial) – inner (adaxial) polarity. (**A** to **D**) REVOLUTA (REV) protein reporter (*proREV::REV-mGFP*) shows nuclear localization of REV, restricted exclusively to the inner epidermis of the wild type (WT) and *as2-7D* sepals. (**A** and **B**) Confocal images of inflorescence meristems show adaxial specific REV expression (yellow arrowheads) in the young sepals of (**A**) WT and (**B**) *as2-7D* mutant. (**C** and **D**) Representative middle stage (Stage 8-9) flowers of (**C**) WT and (**D**) *as2-7D* show no detectable REV expression on the outer epidermis of the sepals in the folds or outgrowths. Images of representative T3s (*n* = 3) show mGFP in green and chlorophyll in greyscale. Scalebar, 50 µm. (**E** to **H**) *FILAMENTOUS FLOWER* (*FIL*) promoter reporter (*proFIL::mCherry-RCI2A*) expression pattern is restricted to the outer sepal epidermis and shows no difference between the WT and *as2-7D* mutant. (**E** and **F**) Confocal images of inflorescence meristems show *FIL* expression pattern in the early floral developmental stages of (**E**) WT and (**F**) *as2-7D* mutant. The membrane localized mCherry is shown in green and chlorophyll in greyscale. (**G** and **H**) Representative middle stage (Stage 7-8) flowers of (**G**) WT and (**H**) *as2-7D* mutant showing the *FIL* promoter reporter signal (mCherry, green). Images are representative of three T3s (*n* = 3). Scalebar, 50 µm. (**I** to **L**) Whole inflorescence images of (**I**) WT, (**J**) *rev-6* mutant, (**K**) a representative transgenic (T1) harboring *pML1::AS2* transgene in *rev-6* and (**L**) *as2-7D* mutant. Note the outgrowths (yellow arrows) on the sepal outer epidermis of *pML1::AS2 rev-6* are similar to those of *as2-7D* and *pML1::AS2* in WT (Fig. 1M). Multiple independent T1s for *pML1::AS2* in *rev-6* were assayed (*n* = 19), all of which show similar outgrowth formation on the sepal outer epidermis in the *rev-6* background.

Since abaxial (outer) identity promoting KANADI transcription factors directly repress *AS2*, and the disruption of KANADI binding site leads to ectopic *AS2* expression in *as2-7D* ^41^, we assessed the sepal phenotypes of *kanadi* loss of function mutants (*kan1-2 kan2-1*). Consistent with previous reports ^38,42,43^, the *kan1-2 kan2-1* double mutants display severe floral development and phyllotaxy defects, characterized by irregular initiation and arrangement of sepals, petals and stamens, and conspicuously enlarged carpels (fig. S9, A and B, E to L). In particular, the *kan1-2 kan2-1* sepals from early developmental stages are strongly radialized with little sepal lamina, and they fail to enclose the floral bud (fig. S9, F to L). These phenotypes are distinct from those observed in *as2-7D* mutants and *pML1::AS2* in wild-type transgenics, which show no such defects (Fig. 1 B, M to P and Fig. 2, A and B). While the mature *kan1-2 kan2-1* sepals are curved downwards, indicative of strong adaxialization (predominantly inner identity), we do not observe the strong folds and associated outgrowths on the outer epidermis of *kan1-2 kan2-1* sepals unlike *as2-7D* (Fig. 1 E and F, fig. S9, C and D). However, as reported previously ^37,38,42,43,66^, we do observe the folds and outgrowths on the outer epidermis of the *kan1-2 kan2-*1 leaves, similar to *AS2* constitutive overexpression phenotypes in the leaves (fig. S9, M to O). The positions of outgrowths typically overlap folds (fig. S9O), which suggests a potential link between buckling and ectopic outgrowth in the *kan1-2 kan2-1* mutant as well.

Overall, our investigations indicate that the outer epidermal buckling exhibited by *as2-7D* sepals may not be associated with an obvious ectopic inner-outer polarity boundary, unlike outgrowths. However, our investigations do not rule out the possibility that other polarity-associated genes are disrupted in *as2-7D*.

### Buckling in *as2-7D* generates outgrowths via ectopic PIN convergence driving auxin maxima formation

In addition to the buckles, we observe the emergence of pointed outgrowths at the ends of the buckles in the later stages in *as2-7D* sepals (Fig. 1F). While constitutive *AS2* overexpression has previously been reported to cause outgrowths ^39^, what local signal triggers the formation of these outgrowths at specific sites remains elusive.

The polar localization of PIN-FORMED 1 (PIN1) protein, an auxin efflux transporter, in the plasma membrane on a particular side of the cell determines the directionality of auxin flow. When PIN1 polarity from multiple neighboring cells converges to the same location, auxin accumulates at that site, creating an auxin maximum ^67^. It was shown that PIN1 convergence sites form at the boundary between the *REV* and *KAN1* expression domains, which coincides with the site of leaf initiation^61^. In the context of the outgrowths, Izhaki and colleagues first showed that the site of outgrowth formation in *kan1 kan2 kan4* triple mutants coincide with the site of PIN1 convergence, indicating that outgrowth formation involves PIN1 ^59^. This was also later confirmed by Abley and colleagues, who showed that PIN1 convergence sites form before the outgrowth initiates in *kan1 kan2* mutant leaves ^68^. Localized auxin import is essential for outgrowth formation since Auxin importer genes *AUXIN RESISTANT 1* (*AUX1*) and *LIKE-AUX1* (*LAX1*) are expressed at the tip of the outgrowths in *kan1 kan2* mutants, while the outgrowths were severely reduced in *kan1 kan2 aux1 lax1 lax2 lax3* hextuple mutants ^68^. Auxin biosynthesis genes *YUCCA1* and *YUCCA4* were also expressed in the region surrounding the outgrowths. Moreover, *CUP-SHAPED COTYLEDON2* (*CUC2*) guides the formation of PIN1 convergence sites and localize auxin at the site of outgrowth formation in *kan1 kan2* leaves, indicating the requirement of boundary genes in driving outgrowth formation^68^.

To test whether outgrowths formed on the outer epidermis of *as2-7D* sepals also involve PIN1 convergence, we analyzed PIN1-GFP localization patterns in the wild type and *as2-7D* mutant. In the inflorescence and early-stage flowers, the PIN1-GFP accumulation pattern in *as2-7D* is similar to that of wild type (Fig. 7, A and B). However, in the later stages we observe PIN1-GFP convergence specifically at the sites where the outgrowths form (Fig. 7, C and D, fig. S10, A and B). In contrast, no PIN convergence sites are observed in the middle of wild-type sepals (Fig. 7, C and D, fig. S10, A and B). We also observe that the auxin-responsive promoter DR5 is specifically expressed where outgrowths are formed in *as2-7D* (fig. S10, C and D), which is consistent with the fact that *PIN1* convergence creates local auxin peaks ^69^.

**Fig. 7.**
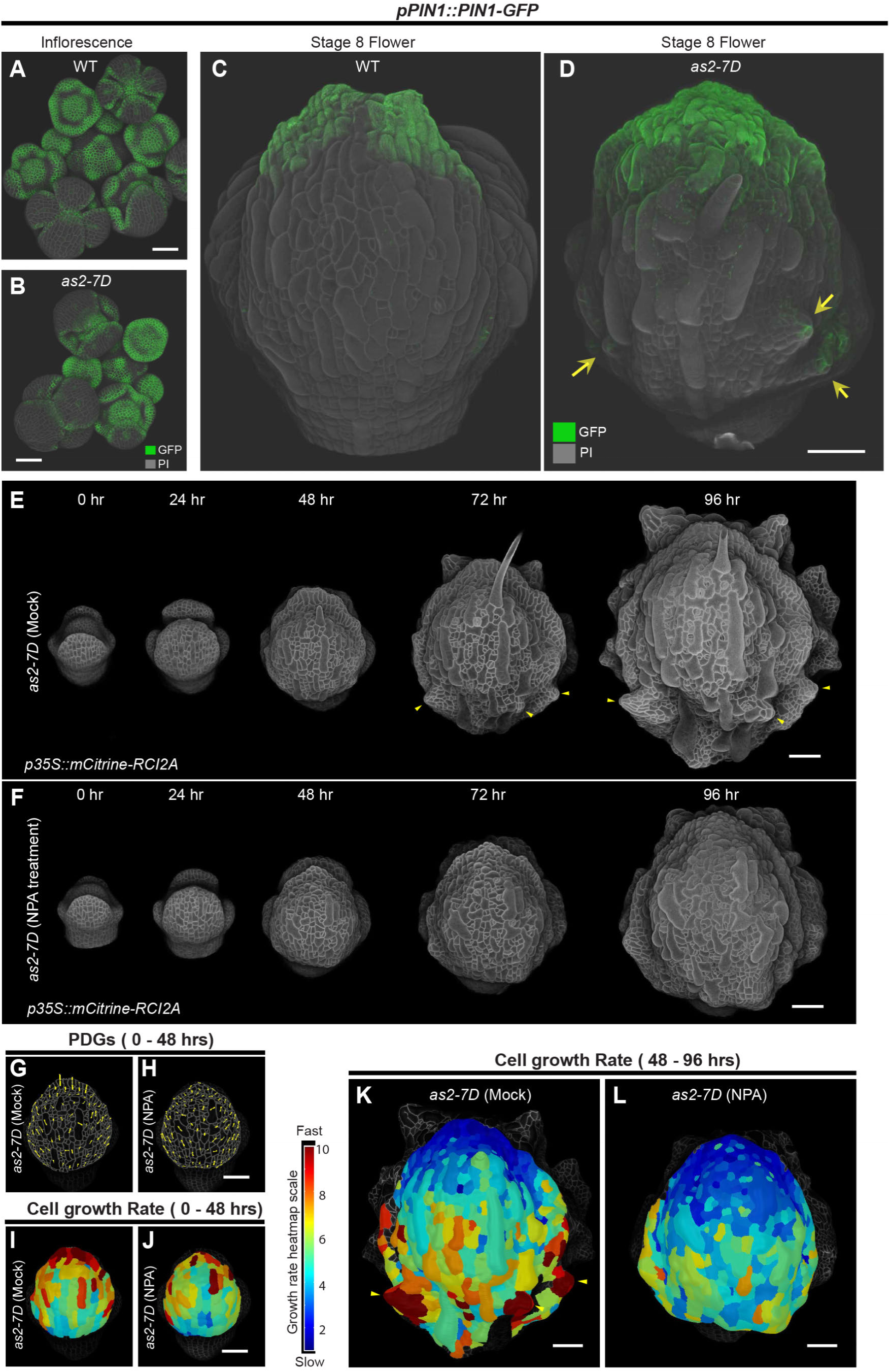
Mechanical buckling drives ectopic PIN convergence creating auxin maxima, causing fast local growth and subsequent outgrowth formation. (**A** to **D**) Confocal stack images showing the expression pattern of auxin efflux carrier protein PIN-FORMED 1 (PIN1) in the wild type (WT) and *as2-7D*. (**A** and **B**) Expression pattern of PIN1 protein reporter (*pPIN1::PIN1-GFP* ^92^) in the inflorescence meristem of (**A**) the WT, (**B**) *as2-7D* mutant. The *PIN1* expression pattern in *as2-7D* is identical to WT in the young floral development stages, wherein PIN1 polarity is directed towards the insipient primordia and growing sepal edges, as described previously ^92^. Scalebar, 50 µm. (**C** and **D**) Expression pattern of *PIN1* in a representative middle stage (stage 8) flower of the (**C**) WT and (**D**) *as2-7D* mutant. Note the expression of *PIN1* at the growing tip of the outgrowths (yellow arrows) in *as2-7D*, whereas no *PIN1* expression is detected in the blade region where the epidermis buckles. In panels A to D, the images are representative of three *pPIN1::PIN1-GFP* and *pPIN1::PIN1-GFP as2-7D* plants (*n* = 3). PI is shown in greyscale and GFP in green. Trichomes were removed from the outer sepal epidermis by image processing. Scale bar, 50 µm. (**E** to **F**) Representative live-imaging series of (**E**) Mock treated *as2-7D* and (**F**) 10µM NPA treated *as2-7D* developing flowers over 96 hours, imaged at every 24-hour interval (representative of *n =* 3 flowers). Note the development of outgrowths from 72-hour time-point onwards (yellow arrowheads) in the mock-treated *as2-7D*, which are absent in NPA treated *as2-7D* flowers. Plants harbor pLH13 (*35S::mCitrine-RCI2A*) cell membrane marker (greyscale). Scale bar, 50 µm. (**G** to **H**) The PDG_max_ tensors (yellow) of the cells on the outer epidermis of (**G**) mock-treated and (**H**) 10µM NPA treated *as2-7D* over the first 48-hour growth period for the representative samples shown in panels E to F. Characteristic misoriented PDG_max_ tensors (relative to the longitudinal sepal axis) on the outer epidermis of *as2-7D* remain unaffected in the NPA-treated samples, thus driving buckling in later stages. Scale bar, 50 µm. (**I** to **L**) Outer epidermal cell growth rates over (**I** and **J**) 0–48-hour growth period and (**K** and **L**) 48–96-hour growth period of the mock and 10µM NPA treated *as2-7D* flowers respectively. Heatmaps are projected onto the later time point. Fast (red) to slow (blue) growth rates are shown. In panel K, note the fast cell growth driving outgrowth formation (yellow arrowheads) which is absent in NPA treated sample (panel L). Scale bar, 50 µm.

The polarity of PIN1 is mechano-sensitive in nature and responds to mechanical stress ^70–72^. In the later stages when *as2-7D* mutants exhibit both buckles and outgrowths, we observe PIN1 convergence specifically in the outgrowths and not the buckles (Fig. 7, C and D, fig. S10, A and B). Since buckling inherently alters mechanical stress distributions ^73,74^, and buckling precedes outgrowth formation, we hypothesized that buckling induced mechanical stress causes PIN1 convergence subsequently leading to outgrowth formation in *as2-7D*. To test our hypothesis, we live imaged growing *pPIN1::PIN1-GFP as2-7D* sepals from initial stages prior to buckling, through the stages where the outer epidermis buckles and outgrowths are evident (Fig. 8).

**Fig. 8.**
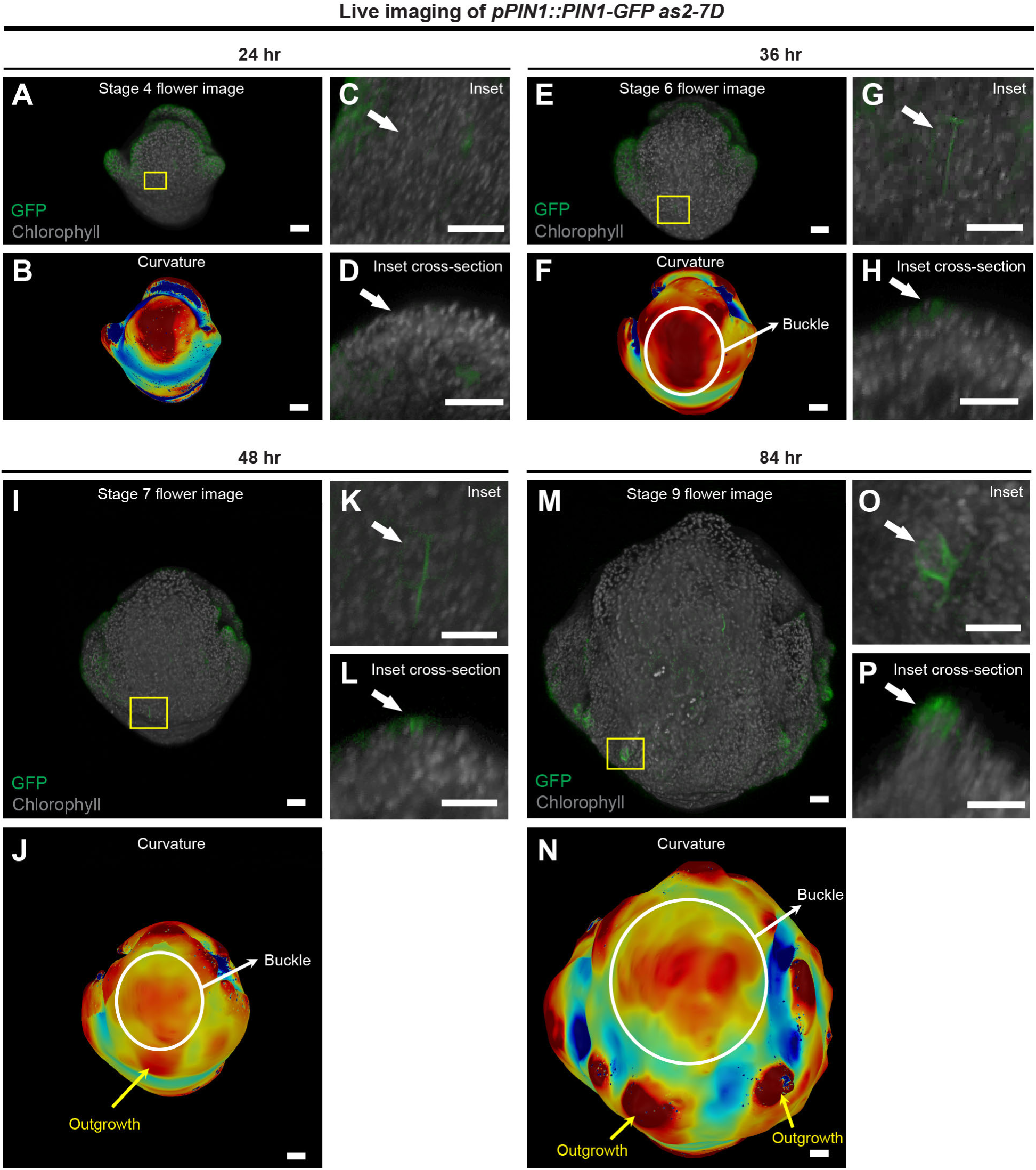
PIN1-GFP converges at the end of a buckle where an outgrowth subsequently forms in *as2-7D* mutants. Confocal live imaging series of a *pPIN1::PIN1-GFP as2-7D* flower, imaged every 12 hours starting from stage 3 (representative of *n = 3*). PIN1-GFP (green) and chlorophyll autofluorescence (gray) are shown. We analyzed the curvature (minimal curvature), in which warm colors denote out-of-plane (bulging) and cool colors represent inward (saddling) deformations. Scalebars, 20 µm. (**A** to **D**) Confocal image of *pPIN1::PIN1-GFP as2-7D* stage 4 flower at 24-hour time point (**A**), and the analysis of its minimal curvature (**B**) is shown. At this stage, the sepal exhibits uniform outward curvature. The inset (**C**) and the cross-section through the inset (**D**) for the region highlighted in panel A (yellow box) depicts the site where an outgrowth will form in the later stages. Note the absence of *PIN1-GFP* expression at this site (white arrow). (**E** to **H**) The *pPIN1::PIN1-GFP as2-7D* flower at the 36-hour time point as it transitions into stages 6 (**E**), and its minimal curvature (**F**) is shown. At this stage, the initial buckle (marked by a white circle) appears as an out-of-plane fold in the center of the sepal blade (deep red on the curvature). No *PIN1* expression is observed at one end of the buckle. The inset (**G**) and the cross-section through the inset (**H**) for the region highlighted in panel **E** (yellow box) is shown. As the outer layer buckles, *PIN1* expression begins to converge in the highlighted region (white arrow), marking the site of the later outgrowth. (**I** to **L**) The *pPIN1::PIN1-GFP as2-7D* flower at the 48-hour time point (**I**), and its minimal curvature (**J**) is shown. At this stage, the outgrowth begins to form (yellow arrow pointing towards the red bulge) at the end of the buckle. The inset (**K**) and its corresponding cross-section (**L**) for the region highlighted in panel E (yellow box) shows the *PIN1* expression associated with the insipient outgrowth (white arrow). (**M** to **P**) The *pPIN1::PIN1-GFP as2-7D* flower at the 84-hour time point as it transitions into stages 9 (**M**), and its minimal curvature (**N**) is shown. The inset (**O**) and the cross-section through the inset (**P**) for the region highlighted in panel **M** (yellow box) where the outgrowth has already formed, is shown. By this stage, *PIN1* expression (white arrow) converges towards the growing tip of the outgrowth.

Initially, when the sepal outer epidermis uniformly curves outwards, no ectopic PIN1 convergence is observed on the sepal blade (Fig. 8, A to D). As the sepal grows, while buckle formation occurs in the middle of the sepal blade, we do not observe PIN1 expression in the region of the buckle (Fig. 8, E and F). Although outgrowth formation is not evident at this stage, we do observe PIN1 convergence at the tip of the buckle, which marks the site of outgrowth initiation (Fig. 8, E to H). In the following time-point, an outgrowth is evident in the region where PIN1 convergence site was formed in the preceding stage (Fig. 8, I to L). As the outgrowth progresses, the site of PIN1 convergence shifts towards the growing tip of the outgrowth, while PIN1 expression continues to be absent in the region where the sepal buckles (Fig. 8, M to P). Collectively, our observations suggest that the tissue buckles first and then PIN1 convergence occurs at the end of a buckle, which precedes the formation of the outgrowths. Our data is consistent with the hypothesis that buckling induced mechanical stress may cause PIN1 to converge and initiate an outgrowth.

We next tested whether PIN-mediated polar auxin transport is required specifically for the formation of outgrowths and not buckles. We treated growing *as2-7D* inflorescences with polar auxin transport inhibitor N-1-naphthylphthalamic acid (NPA) and compared the phenotype to mock-treated samples. While *as2-7D* live-imaged in the presence of NPA exhibited characteristic buckles, the formation of outgrowths was largely attenuated (Fig. 7, E and F, fig. S10, E to H, and movie S4). Buckling continues to occur since the characteristic growth patterns and misdirected sideways PDG_max_ tensors leading to buckling remain unaltered with NPA treatment (Fig. 7, G to J). However, the localized fast growth leading to outgrowth formation was largely reduced in the NPA-treated samples (Fig. 7, K and L). Our results suggest that polar auxin transport is critical for the formation of outgrowth characterized by localized fast growth. Overall, our results are consistent with our hypothesis that mechanical buckling causes ectopic PIN1 convergence, generating local auxin maxima and fast growth, leading to outgrowth formation.

## DISCUSSION

While buckling is critical for generating complex organ shapes required for specialized functional roles ^1–6,16,18^, buckling is not always advantageous, and needs to be circumvented to generate smooth organs like sepals ^75^. Using Arabidopsis sepals as the model, our study suggests that unidirectional growth of both the outer and inner epidermal cells along the sepal longitudinal axis, and similar tissue stiffness across the epidermal layers are key drivers of smooth sepal formation in wild type. Conversely, the outer epidermis of *as2-7D* mutant buckles due to the imbalance of growth and tissue stiffness across the outer and inner epidermal layers.

Overexpression of *AS2* leading to the formation of ectopic outgrowths has been documented by previous studies ^37,39,76^. However, we identify buckling mediated fold formation as an upstream morphological deformity that occurs before the outgrowths are even evident, in the sepals. PIN1 convergence sites form specifically at the tip of buckles, marking the site of outgrowth initiation. Since the Waites Hudson model in 1995 ^57^, the initiation of ectopic outgrowths has been attributed to juxtaposition of abaxial-adaxial identities. We add a second mechanism through which outgrowths can initiate via buckling, which generates auxin maxima via PIN1 convergence. We expect that auxin maxima trigger the gene regulatory networks involved in outgrowth formation, that other studies have extensively characterized.

Although one would intuitively expect *AS2* ectopic expression to affect only the outer epidermis, we find the growth rate and stiffness of the inner epidermis, where *AS2* is normally expressed, is affected as well in *as2-7D*. Changes in inner epidermal growth properties in *as2-7D* could be attributed to the fact that the outer and inner epidermis are connected via the mesophyll and at the sepal margins. Altering growth patterns in the outer epidermis creates mechanical constraints on the inner epidermis, thereby affecting its growth indirectly. Simply put, the outer epidermis of *as2-7D* buckles as the sideways growth of the outer epidermis is limited by the inner epidermis. Conceptually, this resembles trying to fit a cloth inside a box. If the cloth fits perfectly along the edges, it remains smooth; otherwise, the cloth must wrinkle (buckle) to fit inside the box. Our results show how the growth of an epidermal cell layer (or speculatively an animal epithelium ^77^) relative to the tissue to which it is attached, drives organ shape.

While how molecular cues influence tissue mechanics has been relatively well studied ^78–81^, how tissue mechanics inform molecular signals remain elusive. Our study suggests that not only does *AS2* influence tissue mechanics generating buckles, but also that buckles feed back into molecular signals. We show that buckling is associated with convergence of the PIN1-auxin efflux transporter. Our findings support the previous observations from multiple independent studies that mechanical stress affect PIN1 polarity ^70,72,82^. We find an association between the tips of buckles and the establishment of PIN1 polarity convergence sites, causing the formation of auxin maxima to subsequently trigger the initiation of epidermal outgrowths. There is a long-standing debate as to whether primordial initiation in the SAM is also influenced by buckling, a theory originally proposed by Paul Green ^23^. However, PIN1 polarizes before primordia bumps are visible, indicating that PIN1 can converge without buckling ^83^. Our collective evidence suggests that in certain contexts, buckling might be a causative factor, converging PIN1 and generating outgrowths.

### Limitations of the study

First, we used the *as2-7D* mutant as a tool to help us identify how aberrations in cell growth and tissue properties generate buckles, as opposed to smooth sepal surfaces observed in the wild type. However, *as2-7D* being a gain-of-function mutant does not inform much about the native function of *AS2*. We have thus carefully refrained from extrapolating the native role of *AS2* based on our findings.

Second, our numerical simulations are based on a cross-sectional 2D representation of a sepal, since simulating the exact dynamics of a multilayered growing tissue in 3D is extremely difficult. Hence, the Young’s moduli values chosen to achieve outer epidermal buckling in the numerical simulations are not comparable to the differences we observe experimentally. We used the analytical model and numerical simulations as a guide for determining our experimental trajectory, rather than a means to validate our findings.

Third, we used osmotic treatments to assess stiffness of the outer and inner epidermal layers, mainly because the inner epidermis at early stages of sepal development is practically inaccessible via dissection, without inevitably damaging the tissue^84^. As such, alternative methods like AFM or indentation techniques ^85^ which require the tissue to be accessible, are impractical for testing stiffness of the inner epidermis of young sepals. Such methods also typically measure stiffness in highly localized regions, which again does not align with our experimental purpose. Considering these factors, and the fact that osmotic treatments have successfully been used to measure stiffness in multiple studies ^36,55,56,86,87^, we deemed osmotic treatments as a suitable approach.

Fourth, there are currently very limited genetic tools available to orient growth and change tissue mechanical parameters. Although a crude tool, we have chosen to over-expression *KRP1*, since *KRP1* is known to cause anisotropic growth ^53^, and is also unrelated to the core polarity associated genetic network in which *AS2* is involved. However, genetic manipulations are not entirely free from pleotropic phenotypic outcomes. While KRP1 inhibits cell division, the evidence suggests it is not the inhibition of cell division that is suppressing buckling indirectly; the *ML::KRP1 as2-7D* sepals still have dividing cells that could generate buckles and outgrowths, but we do not observe these. Likewise, overexpression of *KRP1* could potentially suppress buckling in *as2-7D* by accelerating differentiation; however, the live imaging shows that the stomatal differentiation is not accelerated in *ML1::KRP1 as2-7D*. Overall, while the data is consistent with our hypothesis, it may not be an absolute proof.

Finally, we acknowledge that we essentially provide negative data where we find no evidence that the buckling is caused due to juxtaposed outer-inner identity. However, as a limitation, there are of course a plethora of genes that *AS2* influences, that we could not test in this study.

## Supporting information

Raw data for making plots

## ACKNOWLEDGEMENTS

We thank Isabella Burda, Frances Clark, Shuyao Kong, Byron Rusnak, Maura Zimmermann, Erich Schwarz, Phoebe Roeder, Scott D. Emr and Stéphane Verger for their valuable comments and suggestions on the manuscript. We thank Olivier Hamant for the *pPDF1::mCitrine-MBD* microtubule marker plasmid. We thank John Paul Alvarez and John Bowman for sharing *kan1-2 kan2-1/+* seeds. We thank Soren Strauss for his guidance with image processing. We thank Mingyuan Zhu for his input on osmotic treatment experiments. We thank Trinh Duy Chi for his input on imaging adult sepals. Finally, we thank Arabidopsis Biological Resource Center for providing seed stocks for this work.

## AUTHOR CONTRIBUTIONS

Conceptualization: AHKR, ASY, LH. Experimentation: ASY, LH, PMK, XH, MH. Imaging: ASY, LH, MH, PMK. Image processing: ASY, LH, PMK, IMB, AMG, RSS. Data analysis: ASY, LH, AK. Analytical modeling: AK, AB. Simulations: AK, MP, AB. Funding acquisition: AHKR, AB, LH. Project administration: AHKR, AB. Supervision: AHKR, AB, LH. Writing – original draft: ASY, LH, AHKR. Writing – review & editing: ASY, LH, PMK, AK, XH, IMB, MH, AMG, RSS, AB and AHKR.

## RESOURCE AVAILABILITY

The raw datasets used for making the graphs are provided in Supplementary Data S1. Primers used for this study are available in Table S1 in Supplementary materials. The transgenic lines for live imaging purposes generated in this study have been deposited with the Arabidopsis Biological Resource Center (ABRC), associated with the following accession numbers: CS73559 (pLH13 L*er*-0), CS73560 (pLH13 *as2-7D*), CS73561 (pLH13 *pML1::KRP1* L*er*-0) and CS73562 (pLH13 *pML1::KRP1 as2-7D*). The meshes and images are publicly available at osf.io (https://osf.io/p5q39/). The R codes used for generating the plots are available at Github (https://github.com/Avilash1993/R_codes_for_plots_as2-7D_sepal_buckling) under GNU General Public License v3.0. Numerical model implementation and the codes for exploration of the parameter space are available under LGPL license at https://gitlab.inria.fr/mosaic/publications/growth_buckling.

## MATERIALS AND METHODS

### Plant Materials

*Arabidopsis thaliana* genotypes used in this study are in Landsberg *erecta* (L*er*) background, which is referred to as the wild type, unless otherwise mentioned. The mutants *rev-6* ^64^ and *as2-5D* mutant (in Col-0 background) ^41^ have been previously described, and were obtained from Arabidopsis Biological Resource Center (ABRC). The *kan1-2 kan2-1* mutant previously described ^93^ was a kind gift from John Alvarez (Monash University). The plants are homozygous for *kan1-2* while *kan2-1* is segregating; the phenotypes were assayed for plants that are homozygous for both *kan1-2* and *kan2-1*. The *as2-7D* mutant in the L*er*-0 background was identified from ethyl methanesulfonate (EMS) based forward genetic screen. Briefly, M2 seeds (second generation) obtained from self-fertilized EMS mutagenized L*er* plants were purchased from Lehle Seeds ^94^. M2 plants were screened under a stereo microscope for sepal defects ^53,95^. The *as2-7D* mutant was isolated based on the lumpy sepal phenotype. A mapping population was generated by crossing the *as2-7D* mutant with a Columbia (Col-0) accession plant. Standard map-based cloning techniques were used to clone the underlying mutation ^96^. The *as2-7D* mutation is a G to A change at base 1,484 upstream of the ATG translational start site of AT1G65620/*AS2*. To reduce background mutations, the *as2-7D* mutant was backcrossed twice to wild-type L*er* plants twice before further analysis. In the backcrosses, we found that *as2-7D* mutant behaves as a dominant allele as the F1s exhibit the *as2-7D* phenotype, although it is not as strong. The *as2-7D* mutation can be PCR-genotyped by amplifying with primer pairs oLH108 and oLH324 at 55°C annealing temperature, followed by digesting the product with EcoRI to produce an 82-bp wild-type product or a 111-bp mutant product. Primers are listed in Supplementary Table S1.

### Plant Growth

Plants were grown in Percival (Percival-Scientific) growth chambers programmed for long day conditions (16-hour light/8-hour dark cycles). Growth temperature (22°C), relative humidity (65%) and fluorescence intensity (∼100 µmol m^-1^s^-1^) were maintained.

For soil-based experiments, plants were grown on soil premixed with Vermaculite (LM111). For stratification, the trays were placed in a 4 °C cold room for three days prior to moving them to the growth chambers. For BASTA (glufosinate ammonium) selection, the soil was autoclaved twice (Program 3-15 min sterilization, 30 min dry, Primus ST-1 Autoclave (Model PSS5-B-MSSD)), and sterilized seeds were planted at a density of 50-100 seeds per pot. For live-imaging and adult-stage phenotyping experiments, one plant was grown per pot (6.5×8.5 inches) to facilitate healthy vegetative growth.

For plate-based experiments, seeds were washed twice with a sterilization solution containing 70% ethanol + 0.1% Triton X-100, replacing the solution after each wash. Two final rounds of washes were provided with 100% ethanol. Seeds were spread on filter paper (Whatman 1, diameter 90 mm) and allowed to dry inside a sterile safety cabinet (Class II, Type A2, Labgard). Dried seeds were evenly distributed on freshly prepared MS plates (1/2 MS basal salt mixture, 0.05% MES w/v, 1% sucrose w/v, pH ∼ 5.8 (adjusted using KOH) containing 0.8% phytoagar. The plates were sealed with 3M micropore tape, wrapped in aluminum foil, and stratified for three days at 4 °C before being moved to the growth chamber.

### Phenotyping whole inflorescences, flowers, and plants

Whole inflorescence and stage 14 flower images were taken either using Canon PowerShot A640 10MP Digital camera attached to Zeiss Stemi 2000-C stereo microscope or Accu-Scope Excelis 4K UHD camera attached to Zeiss Stemi 508 stereo microscope. For inflorescence images, and Stage 14 flowers an optical zoom of 1.0 and 1.6 were used respectively. Whole plant images were captured using iPhone XR with a 26 mm focal length and aperture of f/1.8.

### Schematic diagrams

The schematic diagrams shown in Fig. 1I and Fig. 4 Q to T were hand-drawn using the Notability application in Apple iPad (model A1701).

### Genetic crosses

To generate *pATML1::KRP1 as2-7D*, the *pATML1::KRP1* in L*er* background ^53,54^ were crossed into *as2-7D* mutants. Segregation analysis and confocal microscopy was used to find *pATML1::KRP1 as2-7D* plants. The *pPIN1::PIN1-GFP DR5::VENUS* ^92^ plants and *pPIN1::PIN1-GFP pin1-4* ^92^ plants were crossed into *as2-7D* mutants. The *pPIN1::PIN1-GFP pin1-4* plants (control) exhibit a mild phyllotaxy defect. Genetic segregation analysis and confocal microscopy was used to find *DR5::VENUS as2-7D* and *pPIN1::PIN1-GFP as2-7D* plants.

### Cloning overexpression constructs and reporters

The pLH13 (*p35S::mCitrine-RCI2A,* hygromycin resistance) and pAR169 (*pATML1::RCI2A-mCtrine*, basta resistance) plasma membrane markers have been described previously ^53,97^. The *pPDF1::mCitrine-MBD* microtubule marker ^91^ was a kind gift from Olivier Hamant (RDP institute, ENS de Lyon, France).

For cloning *pML1::AS2*, the AT1G65620/*AS2* coding sequence (600 bps) was amplified from Col-0 cDNA using primers oLH111 and oLH112 by Phusion PCR (NEB) and cloned into pENTR™/D-TOPO™ (Invitrogen™). The *AS2* coding sequence was cloned into the *pATML1* containing gateway vector pAR176 (Basta resistance ^53^) by LR reaction.

For *pML1::REV*, the AT5G60690/*REV* full length coding sequence spanning 2,529 bp was amplified from Col-0 inflorescence cDNA using primer pairs oAS_005 and oAS_006. The resulting PCR product was subcloned into pCR™8TOPO™ vector (Invitrogen™) by TA cloning. LR reaction was used to clone the *REV* coding sequence into pAR176.

To clone *pAS2 ^WT^::GFP-GUS*, the 4.1 kb upstream promoter region and part of the first exon of *AS2* was PCR amplified from L*er*-0 genomic DNA using primer pairs oLH347 and oLH370 and cloned into pENTR™/D-TOPO™ vector (Invitrogen™). The resultant vector was LR recombined into the gateway vector pBGWFS7 ^98^. For cloning *pAS2^as2–7D^::GFP-GUS,* the 4.1 kb upstream promoter region and part of the first exon of *AS2* was PCR amplified from *as2-7D* genomic DNA, and the subsequent cloning steps were similar to cloning *pAS2::GFP-GUS*.

For *proREV::REV-GFP*, the 7.75 kb *proREV::REV* fragment (amplified from L*er*-0) and 1.08 kb mGFP-T35S fragment amplified from *pML1::H2B-mGFP* (pAR180 ^53^) were assembled into the PmeI site of pMOA33 (Kanamycin resistance).

For cloning *proFIL::mCherry-RCI2A*, a 3.92 kb *FIL* promoter fragment was amplified from L*er*-0. This fragment was assembled with a 2.19 kb *omega-mCherry-RCI2A-T35S* fragment amplified from *p35S::mCherry-RCI2A* (pMZ11 ^99^) into the SacI and SpeI site of pMOA34 destination vector (hygromycin plant resistance, ^100^). All the intermediate and final constructs were verified for sequence and directionality by sequencing. Primers are listed in Supplementary Table S1.

### Plant transformation and selection

The binary vector constructs were transformed into *Agrobacterium tumefaciens* GV3101 strain (Rifampicin and Gentamycin resistant) by triparental mating method ^101^. Briefly, two *E.coli* strains, one containing the binary vector and the other containing the helper (conjugative) plasmid, and *Agrobacterium* strain GV3101 were mated together. The GV3101 clones were streaked onto Rifampicin + Gentamycin + binary vector specific antibiotic plates for selection, and positive clones were isolated by colony PCR. The T-DNAs were transformed into plants by Agrobacterium-mediated floral dipping. Transformants were selected either on ½ MS media plates (for Hygromycin and Kanamycin selection) or on soil (for Basta selection). For selecting pLH13 in wild type and *as2-7D*, *proFIL::mCherry-RCI2A* in wild type and *as2-7D* T1s, a final concentration of 25 −30 µg/mL Hygromycin was used. For selecting *proREV::REV-mGFP* in wild type and *as2-7D* T1s, a final concentration of 50 µg/mL Kanamycin was used. Healthy, non-chlorotic plants were transplanted to the soil post two weeks of growth. For selecting *pML1::AS2* in wild type and in *rev-6*, *pML1::REV* in wild type and in *as2-7D*, *pAS2^WT^::GFP-GUS* and *pAS2^as2–7D^::GFP-GUS* in wild type, *pPDF1::mCitrine-MBD* in wild type and in *as2-7D* T1s, BASTA selection was performed on soil grown plants. Seedlings aged 1 to 2 weeks were selected with 100 µg/mL BASTA solution by spraying, and non-chlorotic plants were transplanted to fresh pots after a week. At least twenty independent T1 plants were characterized for each construct. For fluorescent reporter lines, T1 plant was chosen for each line based on fluorescence. Confocal microscopy was used to assess the stability of fluorescence in every generation.

### Flower staging

Floral staging was performed based on Smyth et al ^88^. Briefly, sepal primordia barely arise in Stage 3 and overlie the floral meristem in Stage 4. Sepals enclose the floral bud completely by late stage 6. Flowers beyond stage 6 were staged based on size and morphology ^35^.

### Sepal selected for microscopy

For flower imaging across experiments (live imaging and single time point reporter imaging), we focused primarily on the outer sepal of the flower, unless specified. Briefly, flowers are enclosed within four sepals, wherein the foremost sepal, which faces away from the meristem is denoted as the outer sepal, whereas the sepal that is diametrically opposite to the outer sepal and facing towards the meristem is denoted as the inner sepal. The remaining two sepals positioned on either side are the lateral sepals. For a detailed description, please refer to Roeder et. al ^35^.

### Scanning electron microscopy (SEM)

Scanning electron microscopy was performed as described previously ^53^. Briefly, Stage 14 flowers were fixed in an FAA solution (50% ethanol, 5% acetic acid, and 3.7% formaldehyde) for 4 hours, dehydrated using an ethanol series, and critical point dried. Sepals were dissected, sputter-coated with platinum palladium and then imaged using a LEICA 440 scanning electron microscope.

### Confocal Microscopy

#### Live imaging

For live imaging sepal development, primary inflorescences from six- to eight-week-old plants containing the *p35S::mCitrine-RCI2A* membrane marker (pLH13 in wild type, *as2-7D*, *pML1::KRP1,* and *pML1::KRP1as2-7D* T2s) were excised. Dissection was performed as described previously ^99^, with modifications. Briefly, siliques were removed using surgical scissors (Artman Instruments). Flowers older than stage 7 were dissected using Dumont tweezers (Electron Microscopy Sciences, Style 5) under a stereo microscope (ZEISS Stemi 2000). Dissected inflorescences were inserted upright into small petri-plates containing live imaging media (1/2 MS media supplemented with 1X Gamborg Vitamin mix and 1000-fold diluted plant preservative mixture) with 1.2% w/v phytoagar. The plates were filled with autoclaved water and allowed to stand for two-three hours for the dissected inflorescence to rehydrate. Residual buds were dissected down to Stage 3-4 in water using a needle (BD Micro-Fine^TM^ IV Insulin Syringes). Inflorescences were moved to fresh media plates and transferred to the growth chamber for at least 24 hours for recovery from dissection, prior to imaging. Stage 4 flowers were selected as the 0-hour time point across all live imaging series.

Post recovery, dissected tissues for live imaging were positioned at an angle to ensure the outer sepal of the chosen stage 4 flower directly faces the objective of the confocal microscope. The flowers were submerged in MillQ water containing 0.01% v/v SILWET L-77 surfactant and imaged using Zeiss 710 upright confocal microscope using 20X water dipping objective (Plan-Apochromat 20x/1.0 DIC D=0.17 M27 75mm). The following settings were used: mCitrine (yellow fluorescent protein), excitation laser 514 nm (intensity-2.4-6.0%), emission collection range 519-596 nm and a z-step of 0.415-0.420 µm. For high laser intensity based live imaging of pLH13 in wild type and in *as2-7D* to visualize the adaxial sepal epidermis, imaging settings for mCitrine used: excitation laser 514 nm (intensity-16-18%), emission collection range 519-596 nm and a z-step of 0.244 µm. Post imaging, the plates were returned to the growth chamber, and the process was repeated for each time-point. Inflorescence tissues were transferred to fresh media every 48 hours to prevent fungal contamination.

### NPA treatment assay

Primary inflorescences of six- to eight-week-old pLH13 in *as2-7D* T2s were dissected down to stage 3 and were divided into two groups of 5-6 samples. One group of samples were placed in ½ MS live imaging media plates containing 10 µM NPA. The other group, serving as the control, was placed in media plates containing an equivalent amount of DMSO (20 µL DMSO per 100 ml media). The tissues were dissected 48 hours prior to imaging. Post dissection, the tissues were treated every 12 hours with a 100 µL liquid solution, applied directly above the inflorescence tissue by pipetting. For the 10 µM NPA treatment samples, the liquid solution comprised of 1 µL of 1M NPA+19 µL DMSO+10 µL Silwet in 100 ml water, whereas the control (mock) samples were treated with 20 µL DMSO+10 µL Silwet in 100 ml water. The aim of administering liquid solution was to enhance the effectiveness of the treatment, which was performed every 12 hours throughout the experimental duration. Stage 4 flowers, 48 hours post dissection were selected as the 0-hour time point for imaging mock and 10 µM NPA treated samples. Live imaging was performed as described above, with the samples being moved to fresh mock or 10 µM NPA plates every 48 hours. Imaging settings used: mCitrine (yellow fluorescent protein), excitation laser 514 nm (intensity −2.6-2.8%), emission collection range 519-596 nm, and a z-step of 0.415 µm.

### Propidium Iodide (PI) cell wall staining and imaging

A 5 mg/ml stock solution was prepared for PI staining, with 5 mgs of PI (Sigma) solubilized in 1 mL of MilliQ water. The stock solution was stored in a foil-covered 1.7 ml microcentrifuge tube and refrigerated at 4°C. For staining, stock solution was five-fold diluted with water containing 0.01% v/v Silwet surfactant (PI working solution). Whole inflorescences and flowers below Stage 10 were stained for 6-8 minutes, stage 14 sepals were stained for 12 minutes, and leaves were stained for 25 minutes. Care was taken to ensure that the tissue was fully submerged in the staining solution, and no air bubbles surrounded the tissue. Following staining, the tissue was washed in a fresh solution containing just 0.01% v/v Silwet, prior to imaging. Fresh PI working solution was prepared before every experiment. For imaging PI-stained inflorescences of wild type, *as2-7D* and the transgenics (T1s) of *pML1::AS2* in wild type, *pML1::REV* in wild type, *pML1::REV* in *as2-7D* and the inflorescences, flowers of *kan1-2 kan 2-1* mutants, the following settings were used: excitation 514 nm (laser intensity −18-22%), emission 566-650 nm. For imaging sections of PI-stained *kan1-2 kan 2-1* mutant leaves, samples were excited with 514 nm (laser intensity - 70%) and emission was collected between 566-643 nm range.

### Imaging reporters

For single time point imaging of the reporters, inflorescences were dissected down to stage 3-4. The recovery period was extended to 48 hours to allow ample time for stabilization of the reporter signal. The inflorescences were placed upright in the live imaging media plates with the meristem facing the objective. Prior to taking images of late-stage flowers, the inflorescences were allowed to grow for a few days. For the reporters imaged together with PI, fresh inflorescences were dissected for imaging late-stage flowers. Images were taken with Zeiss 710 upright confocal microscope using 20X water dipping objective.

The following settings specific for each of the reporters were used. For *pAS2 ^WT^::GFP-GUS* and *pAS2 ^as2–7D^::GFP-GUS* images, PI - excitation laser 594 nm (intensity-2%), emission range 599-650 nm and GFP - excitation laser 488 nm (intensity-20%), emission range 493-550 nm, with z-step of 0.240 µm. For *proREV::REV-GFP*, Chlorophyll and GFP were excited with 488 nm laser (intensity-40%), emission range was collected between 647-722 nm for Cholorophyll and 493- 550 nm for GFP. Z-step, 0.240 µm (inflorescences), 0.4 µm (flowers). For *proFIL::mCherry-RCI2A*, Chlorophyll and mCherry were excited with 561 nm laser (intensity-12-15%), emission range was collected between 655-722 nm for Chlorophyll and 578-650 nm for mCherry. Z-step, 0.240 µm. For *pPIN1::PIN1-GFP* single time-point images, PI - excitation laser 594 nm (intensity-2%), emission collection range 599-650 nm and GFP - excitation laser 488 nm (intensity-25-30%), emission collection range 493-550 nm, with z-steps of 0.240-0.5 µm. For *pPIN1::PIN1-GFP as2-7D* live-imaging, GFP – excitation laser 488 nm (intensity – 20-25%), emission collection range 493-538 nm and Chlorophyll – excitation laser 488 nm (intensity – 20-25%), emission collection range 647-722 nm, with z-steps of 0.5 µm. For *pDR5::VENUS*, samples were excited with an argon laser (514 nm), and data were collected in the YFP (520–565 nm) channel. For *pPDF1::mCitrine-MBD,* PI and mCitrine were excited with 514 nm (intensity- 30-32%), and emission range was collected between 603-643 nm for PI and 519-567 nm for mCitrine. Z-step, 0.2 µm.

### Imaging adult sepals

For imaging fully mature sepals, all flowers were dissected off the inflorescence except for a single stage 14 flowers. All the internal floral organs from the stage 14 flower were carefully dissected off, leaving just the outer sepal ^35^ attached to the pedicel. The sepal was wetted in 0.01% v/v Silwet water and mounted between two coverslips filled with 0.01% v/v Silwet water. To prevent the sepal from drying out during imaging, the coverslips were sealed with vacuum grease. Sepals were imaged using 10X objective (C-Apochromat 10X/0.45 W M27). Three-horizontally tiled images were taken with an overlap of 5% and were auto-stitched with a threshold of 0.70 in the ZEISS ZEN software. For imaging fully mature sepals of pLH13 in wild type and in *as2-7D*, mCitrine was excited with 514 nm laser (intensity- 20-22%) and emission was collected at 519-621 nm. For imaging PI-stained sepals of wild types, *as2-7D*, *pML1::KRP1* and *pML1::KRP1 as2-7D*, PI was excited at 514 nm (intensity- 27-30%) and emission was collected at 566-648 nm. For epidermal structure observation, mature flowers harboring pAR169 (*pATML1::RCI2A-mCtrine*) were transverse sectioned using a razor blade. Both mCitrine and chlorophyll were excited using 514 nm laser, and emission was collected at 520-565 nm for mCitrine and 651-700 nm for chlorophyll.

### Osmotic treatments for measuring epidermal stiffness

Osmotic treatments were performed as described previously ^36,55^, with modifications. In brief, primarily inflorescences from pLH13 in wild type, pLH13 in *as2-7D*, pLH13 in *pML1::KRP1* and pLH13 in *pML1::KRP1 as2-7D* six- to eight-week-old plants were dissected down to stage 3 to 4 and placed in ½ MS live imaging media plate. Dissected tissues were allowed to recover for 24 hours in the growth chamber. Early stage 6 buds were precisely chosen for osmotic treatment, since this is the stage right before *as2-7D* outer epidermis starts buckling.

Each bud (corresponding to one replicate) was subjected to osmotic treatment individually and was imaged pre- and post-treatment. Pre-treatment imaging involved staining the bud with a PI working solution for precisely 6 minutes. After washing off the PI stain, the tissue was transferred to a fresh media plate and was oriented to focus on the outer sepal. The tissue was submerged in 0.01% v/v Silwet water, and the pre-treatment image was captured. Imaging settings used for mCitrine – excitation 514 nm (intensity- 20-25%), emission 519-567 nm and PI - excitation 514 nm (intensity- 20-25%), emission 603-643 nm.

For osmotic treatment, the bud was immersed in 0.35M NaCl solution containing 0.01% v/v Silwet for exactly 20 minutes. Post treatment, the bud was re-stained in a PI working solution prepared in 0.35M NaCl (1 part 5 mg/mL PI (solubilized in 0.35M NaCl) and 4 parts 0.35M NaCl solution containing 0.01% v/v Silwet)) for 10 minutes, leading to a cumulative osmotic treatment duration of 30 minutes. This timeframe was found to be optimum for inducing plasmolysis in epidermal cells of early stage 6 flowers, wherein the plasma membrane gets visibly detached from the cell wall at the corners. Re-staining the buds with PI is necessary since NaCl solution erodes off the PI stain. The bud was finally re-positioned in a fresh media plate and submerged in 0.35M NaCl solution. Before image acquisition, the tissue was allowed to stabilize for a few minutes due to its increased fragility post osmotic treatment. The post-treatment image acquisition setting was as follows: mCitrine – excitation 514 nm (intensity- 25-35%), emission 519-567 nm and PI - excitation 514 nm (intensity- 25-35%), emission 603-643 nm. This procedure was iterated for all samples individually.

### Image processing

#### Converting raw images to tiffs, taking screenshots and videos

The raw images, generated either in lsm or czi formats from ZEN imaging software, were converted to tiff files in ImageJ distribution Fiji. For multichannel images, images were split based on channels, which were saved individually. The MorphoGraphX (MGX) ^90^ software was used for image processing and taking the screenshots using the built-in screenshot option in MGX. For taking screenshots of the reporter lines, the reporter image was loaded onto Stack 1, and either chlorophyll or PI was loaded onto Stack 2. Colormaps for Stack 1 and Stack 2 were set to green and gray respectively. Flower images and meristems were rotated in the desired orientation. The voxels not associated with the flower or meristem of interest, were removed using the Voxel Edit tool in MGX. In addition, trichomes from the outer sepal epidermis were removed in a similar manner, for better visualization of the sepal surface. Following this, screenshots were captured. From flower images, trichomes on the outer sepal were removed by image processing for better visualization of the sepal epidermis. The brightness and contrast were uniformly adjusted across samples for any given experiment. Screenshots of MGX generated heatmaps were taken using the default brightness and contrast settings, without any adjustments.

Videos were captured as a series of two-dimensional tiffs files using the built-in “Record movie” option in MGX. The 3D stack images were rotated in the MGX window for the desired duration and time points. For capturing videos involving two genotypes/treatments, the images were loaded onto Stack 1 and Stack 2 and rotated independently. The stack of tiffs generated was loaded onto Fiji and saved as a .avi file without compression. The .avi files were assembled and labeled in Adobe Premiere Pro 2023.

### Generating surfaces for analyzing outer epidermal growth

Making the surfaces and analysis of growth data was performed in MGX as previously described in Zhu et. al. and Hong et. al. ^36,102^, with modifications. The following process was used to generate the surfaces for analyzing cell growth parameters on the outer epidermis of wild type, *as2-7D*, *pML1::KRP1*, *pML1::KRP1 as2-7D*, *as2-7D* mock and 10 µM NPA treatment samples. Briefly, the tiffs were imported onto Stack 1 Main Store in the software. Trichomes on the outer epidermis were removed using the voxel edit tool to get a better surface. Since the outer epidermal morphology of *as2-7D* becomes increasingly irregular in later stages, Level Sets ^103^ was used to extract the structure representing the general flower shape. For this, Level Sets was initialized with an up/down threshold value of 3 to 4. The structure was allowed to evolve (Level Set Evolve) with default parameters, except for “view steps”, which was set to 1 for visualization at more frequent intervals. For performing Edge Detect Angle, the mask was oriented such that the outer sepal directly faces the viewer. The Edge Detect Angle process was then run with default parameters to get a finer structure that closely resembles the image. The holes generated in the structure (which corresponds to places in the image with low membrane marker signal) were closed using Stack/Morphology/Closing - X/Y/Z radius of 3 to 8. A 2.5D surface fitted with a triangular mesh was created from the structure using Marching Cube Surface with a cube size of 3 to 5 µm, followed by smoothing once (Smooth Mesh, passes = 10) and subdividing twice. The membrane signal was projected onto the 2.5D surface with a min. and max. distance range of 1 to 8 µm from the image.

### Generating surfaces for analyzing outer vs inner epidermal growth in wild type and *as2-7D*

To obtain the inner epidermal surface, clipping plane 1 (View Tab/Clip 1), set to a width of 10 to 15 µm, was used to clip the flower image along the longitudinal direction. The voxel editing tool was used to carefully remove the voxels below the inner epidermis of the sepal within the clipping plane region. The clipping plane was then moved to the adjacent region and the same process was repeated, while tilting the plane to ensure voxels from the inner epidermis itself are not removed. Once the voxels below the inner epidermis were removed throughout, the edited tiff was copied back to the main stack and saved. To generate the surface for the inner epidermis, the Z axis of the stack was reversed (Stack/Canvas/Reverse Axes). The stack was blurred using Gaussian blur (X/Y/Z sigma (µm) = 2 to 2.5), and structure was extracted using Edge Detect with a threshold of 2000 to 3000. A triangular mesh surface was created using a cube size of 3 µm, followed by smoothing once (passes = 10) and subdividing twice. Plasma membrane signal from the inner surface was projected onto the mesh surface with a min. and max. distance range of 3 to 8 µm.

To obtain the outer epidermal surface, the processed tiff, prior to reversing the axes, was used. To process the stack, Gaussian blur (X/Y/Z sigma (µm) = 1 to 1.5) followed by Edge detect (threshold of 8000 to 10,000) were used sequentially. Mesh surface was created as described above, and the membrane signal was projected within a 2 to 4 µm distance range from the image.

### Generating surfaces for analyzing growth of outermost mesophyll layer in wild type and *as2-7D*

To enhance the membrane marker signal for the mesophyll cells, we utilized a deep learning approach based on a Convolutional Neural Network (CNN). We used the inbuilt CNN Unet 3D prediction algorithm VijayanUnet.pt ^47^ in MorphoGraphX to precisely predict the cell outlines. The algorithm then uses these predictions for enhancing and de-noising the membrane marker signal and generates a new image based on the predictions.

For both wild type and *as2-7D* mutants harboring pLH13 membrane marker, the CNN algorithm was run on the pre-processed confocal images of the outer sepal, where the parts of the flower not associated with the outer sepal were already removed by voxel editing (see ‘Generating surfaces for analyzing outer vs inner epidermal growth in wild type and *as2-7D*’ section in Materials and Methods). Using the pre-processed sepal images instead of whole flowers for running the CNN algorithm largely reduced computational processing times. Briefly, for wild tpye and *as2-7D* mutants, CNN-generated images of the sepals for each time point were obtained by running Stack/CNN/UNet3D Prediction/VijayanUnet.pt using default parameters. The CNN Unet3D prediction process usually also generates voxels unrelated to the sepals, which were removed using the ‘Voxel edit’ tool. These edited CNN-generated images were then used to run Stack/Mesh Interaction/Annihilate function, to remove the voxels associated with the outer epidermis and the inner layers and obtain the outermost mesophyll layer (layer closest to the outer epidermis). For sepals at 0-hour and 24-hour time points, the outermost mesophyll layer was extracted by setting the minimum distance values for Annihilate function to 5/6 and the maximum distance values to 15. For the 48-hour time points, the minimum and maximum distance value for the Annihilate function was set to 6 and 20 respectively. The resulting images generated post running Annihilate function were used to generate the 2.5D surface for the outermost mesophyll layer. Briefly, for each time-point and genotype, the annihilated image stack was loaded onto Stack 1 and blurred using Gaussian Blur, with X/Y/Z sigma (µm) value of 3. The surface was extracted using Edge Detect threshold of 2000 to 3000. The mesh surface was created using a cube size of 3 µm, followed by smoothing once (passes = 10) and subdividing twice. The membrane signal was projected onto the mesh surface with a minimum and maximum distance range of 7 to 8 µm.

### Lineage tracking

For meshes across all time points, individual cells were manually seeded. Cells were segmented using the watershed algorithm, and unlabeled vertices were fixed by running Cell Mesh/Fix Corner Triangles. Cell lineages were parent tracked to the earlier time-point to analyze cell growth. Parent (lineage) tracking involves transferring the parent labels from cells in the earlier time point onto the daughter cells at the later time point. For this, the meshes corresponding to the earlier (Mesh 1) and later (Mesh 2) time points were opened side by side on the same MGX window. A few daughter cells in Mesh 2 were manually parent labeled, and using these as landmarks, the remaining daughter cells were auto parent labeled (by running Deformation/Morphing/Set Correct Parents followed by Deformation/Mesh 2D/Auto Parent Labelling 2D). Errors in segmentation or parent tracking were manually corrected. Parent tracking was checked for accuracy using Cell Axis/PDG/Check Correspondence, which checks whether the cell junctions are identical across time points. The final parent labels were saved as .csv files. A custom R script (combinelineages.r) was run to automatically correlate the lineage information through multiple time points, followed by manual correction.

### Analyzing cell growth parameters

Meshes corresponding to the earlier (Mesh 1) and later time points (Mesh 2) were loaded side by side in the same MGX window, with parent labels loaded onto Mesh 2. Cell growth rate was calculated based on the total area of the daughter cells in the second time point divided by the area of the parent cell in the earlier time point. Growth rate heatmaps were generated using Heat Map/Heat Map Classic/Heat Map with the Type set to Area. The cell division rate was calculated based on the total number of daughters formed in the second time point for each cell in the earlier time point. Cell division heatmaps were generated using Lineage Tracking/Heat Map Proliferation. The values for growth rate and cell division rate were exported to .csv files.

The Principal Direction of Growth (PDG) uses information of corresponding junctions between the earlier and later time points, to compute the direction of maximum and minimum growth of each individual cell in the earlier time point ^90^. In the figures, the PDGs represent the maximum growth direction (PDG_max_) tensor (yellow lines), the length of which is directly proportional to the magnitude along the direction of maximum growth. The PDGs were calculated for the cells in the earlier time point (using the information of parent labels from the later time point), the PDG_max_ tensors were projected onto the later time point for visualization. For displaying PDG_max_, the values for line width and line scale were set to 5 and 3 respectively.

The Angle of Deviation of the PDG_max_ tensor was calculated relative to the longitudinal axis of the sepal. The sepal longitudinal axis was set using a Bezier line on the earlier time point, since PDGs and deviation angles are calculated on the earlier time point. Briefly, a new Bezier grid (Misc/Measure/New Bezier) was created, which was then collapsed into a Bezier line (Misc/Bezier/Collapse Bezier Points). The supporting points on the Bezier line were placed on the mesh in a manner such that the Bezier line is parallel to the sepal longitudinal direction, and the line shape fits the curvature of the sepal. The custom axis for each cell was set using this Bezier line (Cell Axis/Custom/Create Bezier Line Directions) followed by smoothing (Smooth Custom Directions). The PDGs were loaded onto the mesh, and the angle of deviation of the PDG_max_ from custom X direction (which represents the longitudinal axis) was calculated using Cell Axis/Custom/Custom Direction Angle. The values of the angle of deviation heatmap were exported as .csv files.

### Analyzing osmotic treatment induced epidermal shrinkage

Our osmotic treatment conditions were optimized such that the cell walls shrank, but the cell membrane barely plasmolyzed. Post osmotic treatment, the membrane just visibly pulled away from the cell wall corners, thereby rendering the wall and membrane area almost comparable. Since the PI signal (corresponding to the cell wall) is very weak post-treatment, especially for the inner layers, the PI and mCitrine (membrane marker) images were merged as a single image MGX (Stack/Multistack/Merge Stacks).

For analyzing epidermal stiffness, we assume that the surface tension across genotypes is comparable. The shrinkage in area of the epidermal layers along a comparable longitudinal section was analyzed to infer tissue stiffness. For this, the pre- and post-treatment merged stacks for any given sample were opened side by side in the same MGX window. Using clipping plane 1 (Clip 1), longitudinal cross-section corresponding to the same regions were extracted from the flower images. The tiffs were processed using Levels Sets to extract the global shape. Mesh surfaces were made with a cube size of 3 µm, which was smoothed once (10 passes) and subdivided twice. Signal was projected with a 1.5 to 3 µm distance range from the image. From the cross-section surface, the outer epidermis and inner epidermis were seeded and segmented as single “cells”. Care was taken to ensure that the epidermal region being seeded was comparable between the pre- and post-treatment surfaces. For measuring shrinkage, parent labels from the pre-treatment mesh were transferred onto the post-treatment mesh, following which the shrinkage ratios were quantified (Mesh/Heatmap/Heatmap Classic). The areal shrinkage values were stored as a .csv file, which was converted to percentages for generating heatmaps shown in the figures.

### Computing minimum curvature

In MorphographX, minimal curvature analysis quantifies the least degree of curvature at any given location on the surface. The output is displayed in the form of a heatmap where warm colors represent regions of higher curvature or outward bulging, whereas cool colors represent regions of lower curvature, such as inward deformations (saddles). For calculating the minimal curvature for the sepals from the live imaging series of *pPIN1::PIN1-GFP as2-7D* flowers, we used the images obtained from the chlorophyll autofluorescence channel to construct the mesh surfaces. Briefly, the chlorophyll image corresponding to a given flower (for which the minimal curvature is to be analyzed) was loaded onto Stack 1. The structure representing the general shape of the flower was extracted using Level Sets (Stack/Lyon/Init Level Set), with an up and down threshold values set to 4. The structure was allowed to evolve (Stack/Lyon/Level Set Evolve) with default parameters. The triangular 2.5D mesh was created using Mesh/Creation/Marching Cube Surface with 5 µm cube size, followed by one round of smoothing (Mesh/Structure/Smooth Mesh, passes = 5) and subdividing (Mesh/Structure/subdivide). Minimal curvature was then calculated using Mesh/Signal/Project Mesh Curvature, selecting Type = Minimal and setting the neighborhood radius value to 50 µm. For visualizing the resulting heatmap, the transfer function was set to “Jet”.

### Analytical model for a growing trilayer

#### System and assumptions

We consider a growing elastic system made of three layers, corresponding to the inner epidermis (denoted i), mesophyll (m), and outer epidermis (o) of a lateral organ. We consider that the three layers may grow at different (specified or target) rates and have different elastic moduli. To simplify the formulae, we assume that the epidermal layers have similar thicknesses. We also assume that the Poisson ratios of all layers are equal (value *v*). Finally, we neglect the bending modulus of each layer considering their small thickness. Note, these three assumptions do not affect the qualitative behavior of the system.

We assumed *E_i_*, *E_m_*, and *E_o_* be the values of elastic modulus, *ℎ*,*ℎ_m_*, and *ℎ* the values of thickness,and *g_i_*,*g_m_*, and *g_o_* as the values of strain associated with target (specified) growth rates for the inner epidermis, the mesophyll, and the outer epidermis, respectively.

The differences in growth rates between the layers may lead to changes in system geometry. As the three layers are bound to each other, they have approximately the same curvature *c* (corresponding to the radius of curvature measured at the middle of the mesophyll), while the strain in the inner and outer layers can be deduced from the strain in the middle layer *E* using curvatures and assuming that the reference state is flat. From kinematics of the system, the elastic (residual) strains in the three layers are given by

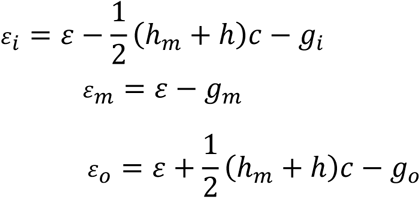

The elastic energy of the system takes the form

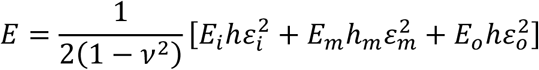

Minimizing the elastic energy with respect to strain and curvature yields their values at equilibrium. In particular, using the notation

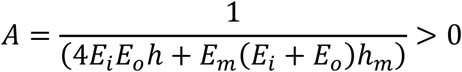

we obtain the equilibrium values for the strain

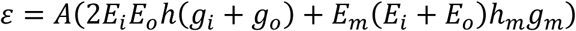

and the curvature

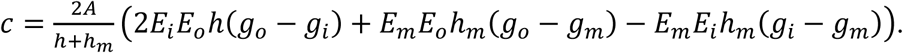

This curvature is in general non-zero, the system curves with the abaxial side on the convex side as soon as the abaxial growth rate is sufficiently larger than the adaxial growth rate.

To determine whether buckling of the abaxial epidermis may occur, we compute the elastic (residual) strain in this layer at equilibrium:

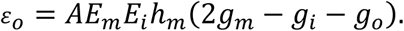

*ε_o_* can become compressive (<0) only if 2*g_m_* − *g_i_* − *g_o_* < 0. When it falls beyond a threshold ^104^, this leads to buckling of the abaxial side, with patterns as described ^105^, depending on the anisotropy of the residual strain.

Interestingly, the elastic strain of the adaxial epidermis,

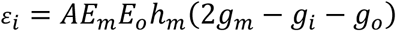

always has the same sign as that of the abaxial epidermis. This means that either the two epidermal layers are in compression, or the two epidermal layers are in tension. A situation where one epidermis would be in tension and the other would be in compression does not correspond to mechanical equilibrium. Nevertheless, the amount of compression may differ between the two epidermal layers: the ratio of the elastic deformations is related to the ratio of Young’s moduli

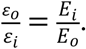

Accordingly, it is possible that only one of the two epidermal layers reaches the buckling threshold. For instance, a much smaller Young’s modulus for the outer epidermis compared to the inner epidermis would lead to buckling of the outer epidermis exclusively.

Finally, we mention that the elastic strain in the mesophyll has an opposite sign

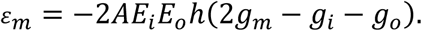

This means that as soon as the epidermis is under compression, the mesophyll is under tension and vice versa.

### Numerical model for a growing layered sepal

We again considered a growing elastic system comprising of three layers (inner epidermis (i), mesophyll (m), and outer epidermis (o)). To better represent the geometry of the transversal section of a sepal, we also considered two lateral semi-circles (the edges) to connect the inner and outer epidermis (Fig. 2, I to K). These semi-circles are denoted “end”, while the region enclosed by the epidermal layers and the edges denote the mesophyll of the trilayer. This geometry is characterized by three parameters: the epidermal layer thickness *ℎ*, the mesophyll thickness *ℎ_m_* and the width of the trilayer part *L*.

We adopt the framework of morpho-elasticity ^106^, assuming that the residual stresses are solely created by a local growth deformation tensor *F_g_*, and the observed deformation gradient can be decomposed multiplicatively on the elastic and the growth deformation tensors. In this framework we consider a quadratic energy density in the Green-Lagrange strain tensor *E_e_*,

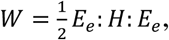

where *H*, the elasticity tensor of rank four, represents the mechanical properties of the material. The proposed model assumes all regions of the geometry to be isotropic materials. Therefore, the elasticity tensor field *H*, is characterized by independent parameters: the Young’s moduli of the four regions (*E_i_*, *E_m_*, *E_o_* and *E_end_*), and the common Poisson ratio *v*. An equation for the time evolution of the growth tensor *F_g_* completes the model,

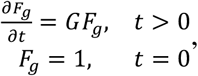

with *G* the growth rate tensor. We consider here that the growth rate has both an imposed target (or specified) component *G_T_*, as well as a component that stands for an influence of the elastic strain on growth,

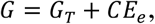

with *C* being the extensibility (assumed to be scalar). We assume that the target growth is unidirectional along the trilayer, so that the target growth rates *y_i_*, *y_m_*, *y_o_* and *y_end_* for the inner epidermis, the mesophyll, the outer epidermis, and the edge respectively are the only independent parameters encoded in the target growth rate tensor *G_T_*. As for the extensibility *C*, we have four additional parameters *C_i_*, *C_m_*, *C_o_* and *C_end_* corresponding to the four regions.

In comparison with the analytical model presented above, all geometrical and mechanical parameters are the same, while the growth parameters *g_i_*,*g_m_*, and *g_o_* can be seen as the relative growth cumulated until time *t*, which can be approximated as when *C* = 0.

The model of the growing layered sepal was solved using the Finite Element Method and implemented with the FEniCS computing platform ^107^ with its Python interface, with a special use of the BVPy Python library ^108^.

We simulated the deformation of the defined geometry when the outer epidermis is growing (*y_o_* ≠ 0), while the other regions do not (*y_i_* = *y_m_* = *y_end_* = 0). This is one of the simplest situations, when the mean growth rate of the two epidermal layers is higher than the growth rate of the mesophyll, and according to the predictions of the analytical model the epidermal layers are under compression. This first condition for buckling being satisfied, we studied whether buckling indeed occurs depending on the moduli of the mesophyll and epidermal layers (fig. S3). To validate the approach, we also compared the equilibrium curvature predicted by the analytical model to the mean midline curvature of the deformed sepal in the numerical model (see fig. S4).

For all our simulations we considered fixed values for the length *L =* 10, and thickness 2*ℎ* + *ℎ_m_* = 5 of the sepal, the target growth rate of the outer epidermis *y_o_* = 0.1, and the Poisson ratio *v* = 0.49. The value of the extensibility is chosen low for all regions but relatively higher for the mesophyll region, *C_o_* = *C_i_* = *C_end_* = 0.01 and *C_m_* = 0.1. The Young’s modulus of the mesophyll is chosen as the lowest among all regions, *E_m_* = 1, while the Young’s modulus of the edge epidermis is a high value, *E_end_* = 10^6^. As for the Young moduli of the outer and inner epidermis, we performed an exploration of the parameter space defined by these two extreme values, (*E_o_, E_i_*) ∈ [1, 10^6^] × [1, 10^6^]. We present the results of this analysis for different values of the epidermis thickness, ℎ ∈ {0.1,0.2,0.3,0.4} (fig. S3 and S4).

### Graph plotting

Graphs were generated in R (version 4.2.2 (2022-10-31 ucrt)) using ggplot, raw data for which is provided in Data S1.

## SUPPLEMENTARY FIGURE TITLES AND LEGENDS

**Fig. S1.**
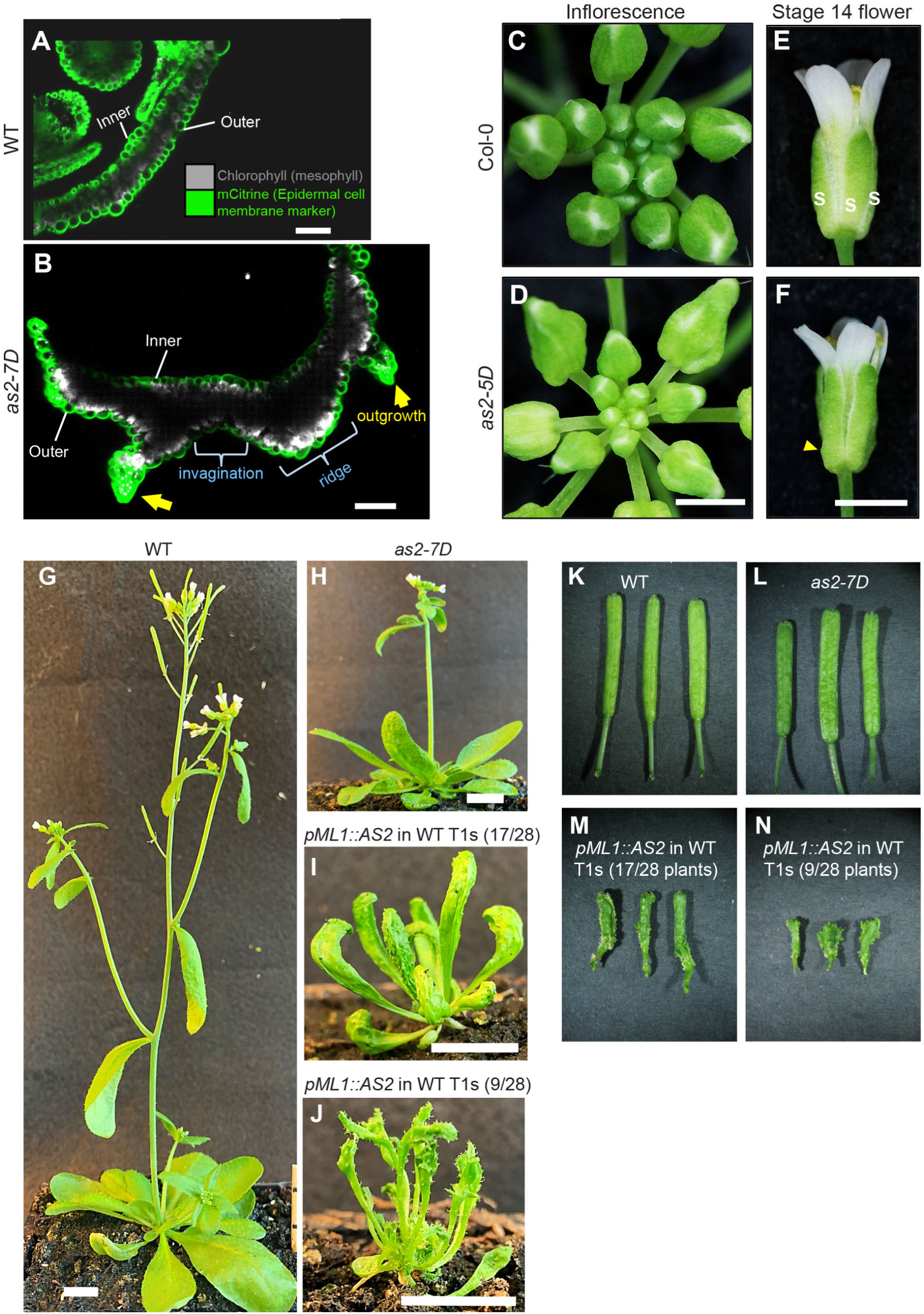
*AS2* overexpression causes characteristic ridges, invaginations, and outgrowths on the outer epidermis, associated with Main Fig. 1. (**A** and **B**) Representative optical cross-sections of Stage 11 flowers of (**A**) the wild type (WT) and (**B**) *as2-7D* mutant. Images are representative of T2s harboring pAR169 (*pATML1::RCI2A-mCitrine*), an epidermal layer specific plasma membrane marker. Green and grey represent epidermal cells and chlorophyll containing mesophyll cells respectively. WT sepals (A) have a smooth shape, while the *as2-7D* sepal (B) outer epidermis exhibit outward protrusions (ridges), inward indents (invaginations) and pointed epidermal extensions (outgrowths). Note that the outgrowths lack mesophyll cells (grey) underneath whereas mesophylls occupy the space beneath ridges and invaginations. (**C** and **D**) Whole inflorescence images of (**C**) Col-0 (control) and (**D**) *as2-5D* (in Col-0, isolated by Wu et. al ^41^). Images are representative of *n =* 3 inflorescences. Scalebar, 1 cm. (**E** and **F**) Stage 14 flowers of (**E**) Col-0 and (**F**) *as2-5D* mutant, representative of *n =* 3 flowers. Yellow arrowhead points toward an outgrowth. Note the mild nature of outgrowth formation in *as2-5D* mutant as compared to *as2-7D* (Fig. 1). Scalebar, 1 cm. (**G** to **J**) Representative six-week-old adult plants of (**G**) WT, (**H**) *as2-7D* and (**I-J**) *pML1::AS2* in WT T1s. The transgenics show strong (17/28 plants, panel I) to extreme (9/28 plants, panel J) developmental defects, categorized qualitatively by the extent of leaf deformity. The transgenics exhibit outgrowths on the lower (abaxial) surface of the leaves as well, in addition to severely stunted growth. Scalebar, 1 cm. (**K** to **N**) Fully formed siliques of (**K**) WT, (**L**) *as2-7D*, (**M-N**) *pML1::AS2* in WT T1s. Note the outgrowths on the silique epidermis. In panels M and N, siliques are representative of strong and extreme categories, described for panels I and J respectively. Three siliques on each panel correspond to three independent plants. Total number of independent transgenics (T1s) assessed *n =* 28.

**Fig. S2.**
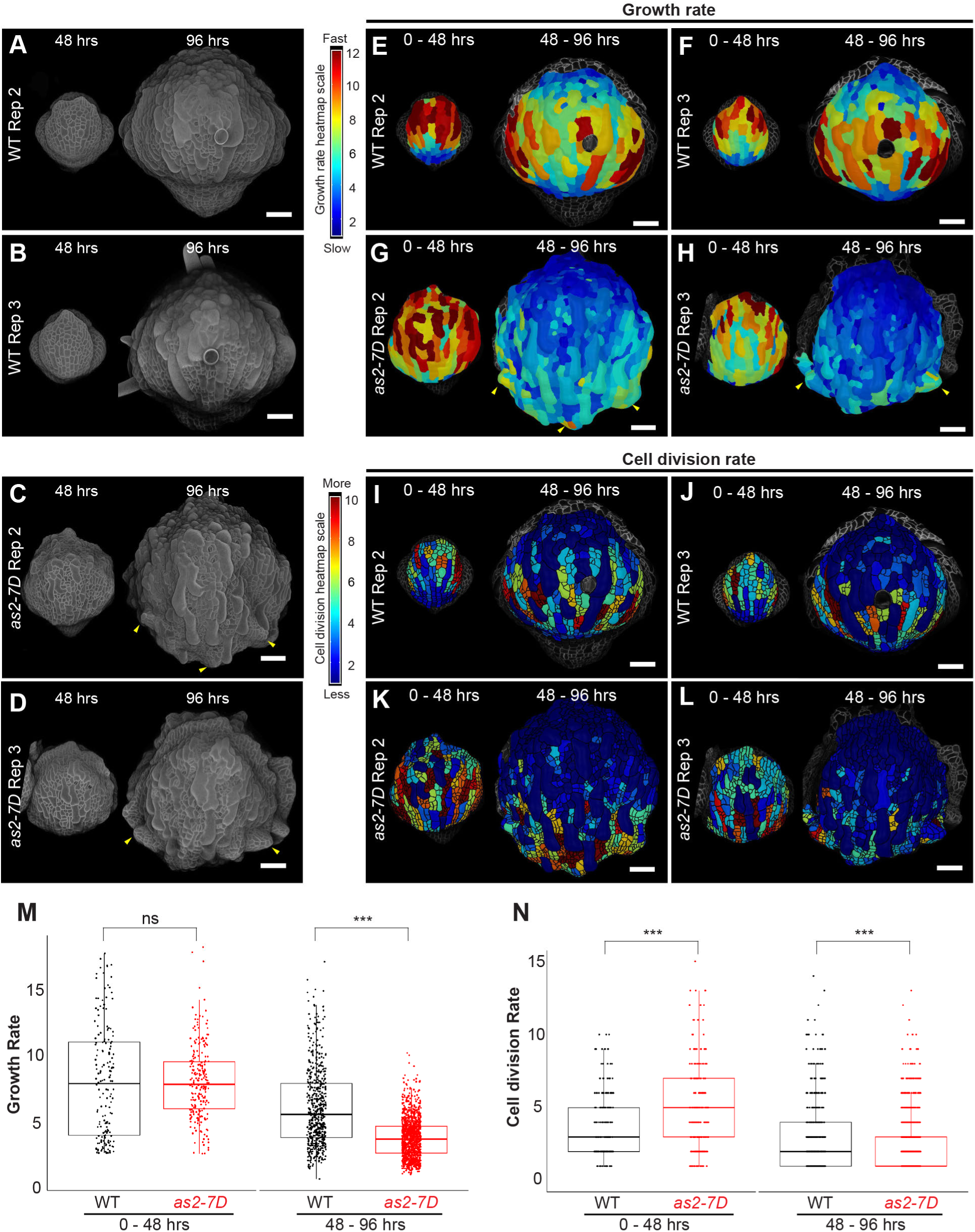
The outer epidermis of *as2-7D* exhibits loss of stereotypical growth patterns, associated with Main Fig. 2. (**A** to **D**) Confocal stack images of (**A**) Wild type (WT) replicate 2, (**B**) WT replicate 3, (**C**) *as2-7D* replicate 2 and (**D**) *as2-7D* replicate 3 developing flowers, transformed with the pLH13 (*35S::mCitrine-RCI2A*) membrane marker. Images correspond to the 48-hour (left) and 96-hour (right) time points from WT and *as2-7D* live imaging experiment (*n =* 3). Yellow arrowheads show outgrowths. Trichomes were removed from the outer sepal epidermis by image processing. Scalebar, 50 µm. (**E** to **H**) Cell area growth rates of (**E**) WT replicate 2, (**F**) WT replicate 3, (**G**) *as2-7D* replicate 2 and (**H**) *as2-7D* replicate 3. The heatmap is projected onto the later time point. Red to blue gradient shows fast to slow growth. Yellow arrowheads point towards the outgrowth which exhibit localized fast growth. Scalebar, 50 µm. (**I** to **L**) The cell division rates of (**I**) WT replicate 2, (**J**) WT replicate 3, (**K**) *as2-7D* replicate 2 and (**L**) *as2-7D* replicate 3. Red to blue gradient shows more to less division. Cell division over 0-48 hours is shown on the left and over 48-96 hours on the right. Scalebar, 50 µm. (**M** and **N**) Graphical representation of (**M**) cell growth rates and (**N**) cell division rates across all three independent biological replicates of the WT (black dots) and *as2-7D* mutant (red dots). Dot represents the growth rate (M) and division rate (N) of a given cell lineage. Dot plots are superimposed with box plots, where the central line indicates the median. Left and right panels display data for 0-48 and 48-96 hours, respectively. Note the reduced spatial variability in the growth rates of *as2-7D* (panel M). Lines above data points indicate statistical comparisons. The *p-values* for Bonferroni corrected pairwise t-test (two-tailed) are shown. *p-values*: ***<0.0001, **<0.001, *<0.05, ns = non-significant. Exact *p-values* for growth rates: WT vs *as2-7D* for 0-48 hours, 0.9884; WT vs *as2-7D* for 48-96 hours, 1.565172 × 10^-93^. Exact *p-values* for cell division rates: WT vs *as2-7D* for 0-48 hours, 7.800489 × 10^-05^; WT vs *as2-7D* for 48 - 96 hours, 9.47882 × 10^-07^.

**Fig. S3.**
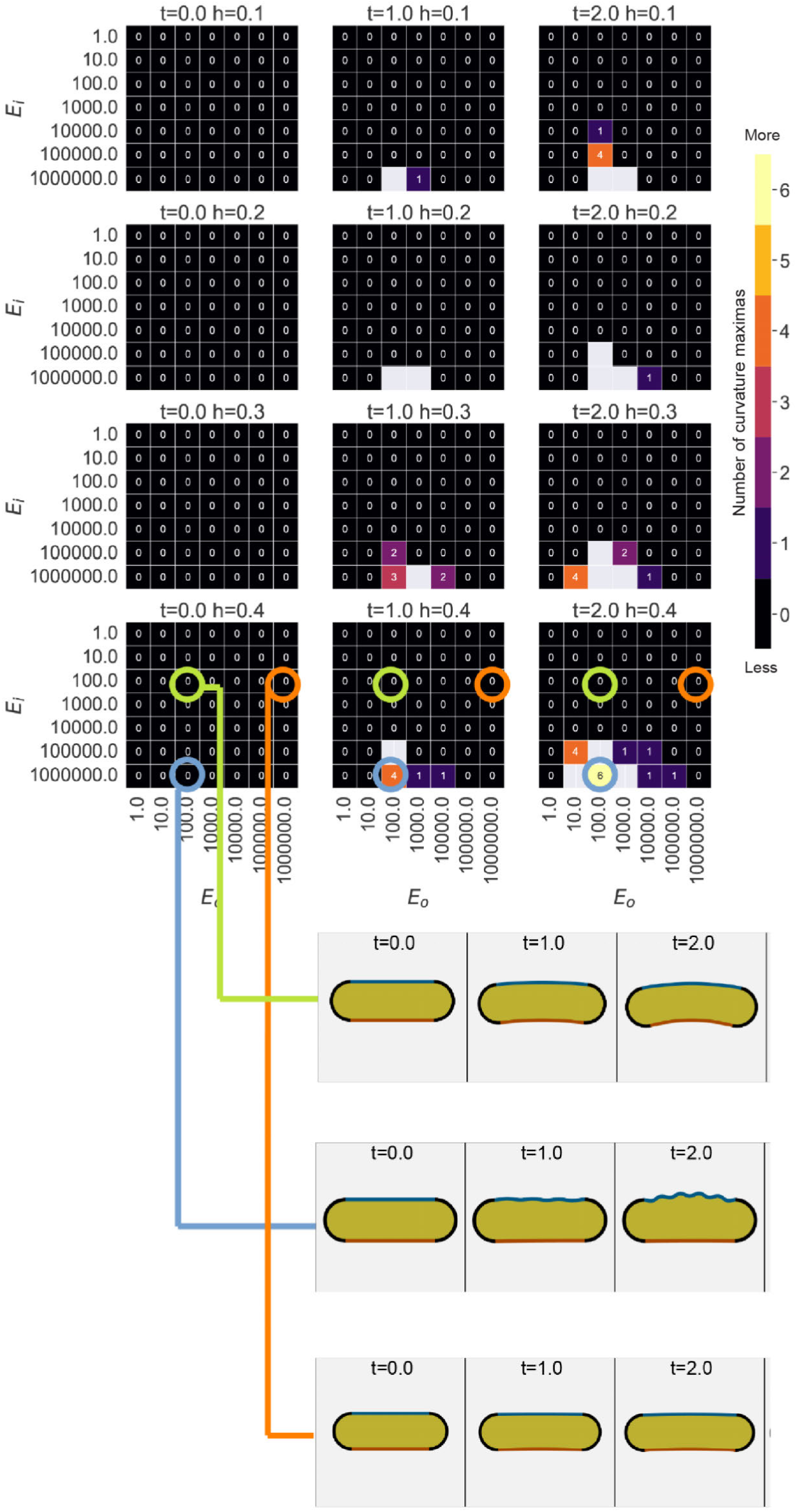
Numerical simulations for a sepal cross-section with the outer epidermis growing in width, showing that buckling is determined by the relative Young’s moduli of the outer and inner epidermis, associated with Main Fig. 2. Parameter space exploration for the numerical model of the growing sepal with three layers, corresponding to the outer epidermis (blue), mesophyll (yellow) and the inner epidermis (orange) as shown in Fig. 2, I to K. As a proxy for detecting whether outer epidermis buckles or not, we monitor the number of curvature maxima on the surface of the outer epidermis. A value for maxima higher than one indicates the occurrence of a buckling event. Block rows in the grid correspond to different epidermal thickness values ℎ ∈ {0.1,0.2,0.3,0.4}, while block columns correspond to different time points during growth. In each block the Young’s moduli of the outer and inner epidermis are variable, (*E_o_, E_i_*) ∈ [1, 10^6^] × [1, 10^6^]. All other parameters are kept constant: *L* = 10, 2*ℎ* + *ℎ_m_* = 5, *y_o_* = 0.1, *y_i_* = *y_m_* = *y_end_* = 0, *v* = 0.49, *C_o_* = *C_i_* = *C_end_* = 0.01, *C_m_* = 0.1, *E_m_* = 1, *E_end_* = 10^6^. Holes in the grid are simulations for which the numerical solver diverged for the given parameter set and time point.

**Fig. S4.**
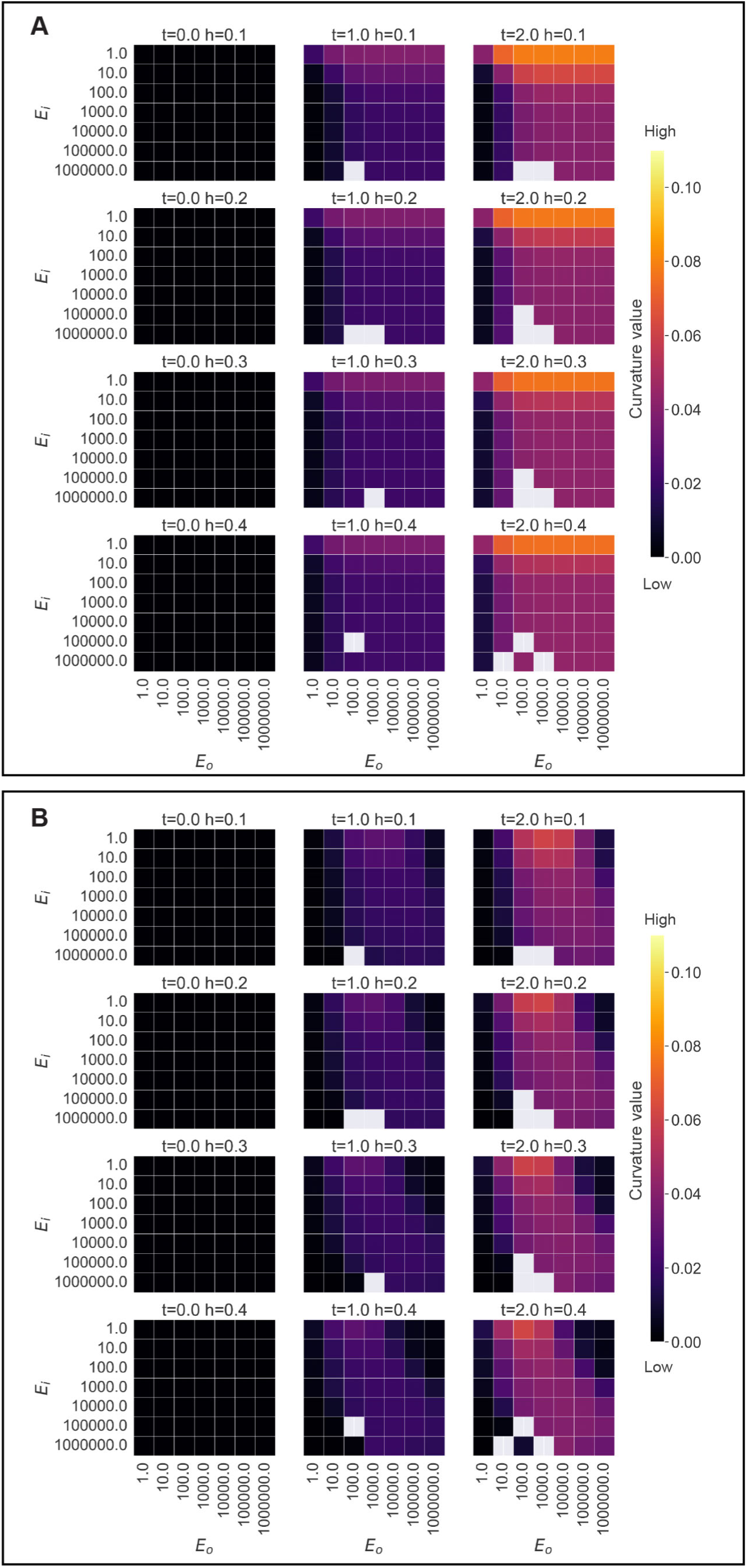
Cross validation of the analytical and numerical models for the growing layered sepal, associated with Main Fig. 2. (**A**) Equilibrium curvature value predicted by the analytical tri-layer model. (**B**) Mean midline curvature of the deformed sepal in the numerical model. (**A** and **B**) Parameter values and the organization of the grids are the same as described in fig. S3. The curvature values in the numerical model deviate from the analytical model prediction primarily when the inner epidermis is much softer than the outer epidermis and the edges. We attribute this deviation to the difference in geometry between the two models, namely the presence of a rigid edge in the numerical model, that connects the inner and outer epidermal layers.

**Fig. S5.**
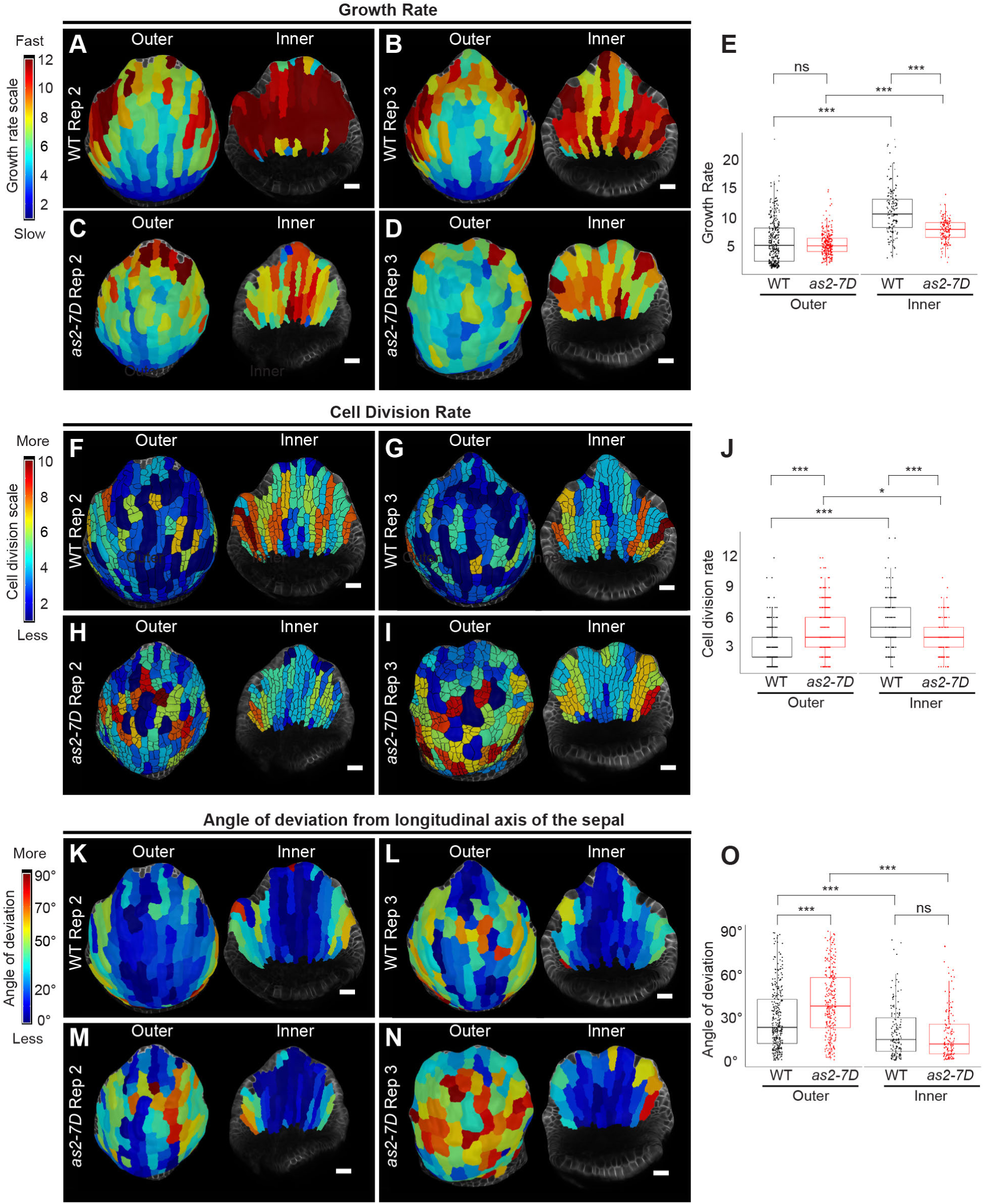
Slower growth on the inner epidermis together with misdirected outer epidermal growth trigger buckling in *as2-7D*, associated with Main Fig. 3. (**A** to **D**) Growth rate heatmaps for (**A**) Wild type (WT) replicate 2, (**B**) WT replicate 3, (**C**) *as2-7D* replicate 2 and (**D**) *as2-7D* replicate 3 developing sepals. Red to blue heatmap gradient shows fast to slow growth. Outer (left) and inner (right) epidermal growth rates over 48 hours is projected as a heatmap onto the later time point. Scalebar, 20 µm. (**E**) Graph shows cell growth rates over a 48-hour period for the WT (black) and *as2-7D* (red) mutant (*n =* 3 biological replicates). Each dot represents the growth rate for a given cell lineage. Dot plots are overlayed with boxplots, where the central line represents the median. The *p-values* for Bonferroni corrected pairwise t-test (two-tailed) are shown. *p-values*: ***<0.0001, **<0.001, *<0.05, ns = non-significant. Exact *p-*values for epidermal cell growth rate: WT outer vs inner, 5.751359 × 10^-35^; *as2-7D* outer vs inner, 3.039502 × 10^-25^; WT vs *as2-7D* outer, 0.1011; WT vs *as2-7D* inner, 8.496122 × 10^-16^. (**F** to **I**) Cell division heatmaps for (**F**) WT replicate 2, (**G**) WT replicate 3, (**H**) *as2-7D* replicate 2 and **(I)** *as2-7D* replicate 3. Red to blue color gradient shows more to less cell division. The data is presented in the same format as described for panels A-D. Scalebar, 20 µm. (**J**) Graph shows cell division rates for the WT and *as2-7D* mutant (*n =* 3 biological replicates) Dots represent the number of cell divisions for each cell lineage. Data presentation and statistical comparisons are same as described above for panel E. Exact *p*-values for epidermal cell division rates: WT outer vs inner, 4.08502 × 10^-28^; *as2-7D* outer vs inner, 0.004; WT vs *as2-7D* outer, 3.677026 × 10^-23^; WT vs *as2-7D* inner, 1.865456 × 10^-06^. (**K** to **N**) Heatmap representing the angle of deviation of PDG_max_ tensors from the sepal longitudinal axis for (**K**) WT replicate 2, (**L**) WT replicate 3, (**M**) *as2-7D* replicate 2 and (**N**) *as2-7D* replicate 3. Scalebar represents a deviation of 90° in red and 0° (no deviation) in blue. Scalebar, 20 µm. (**O**) Graph showing the angle of deviation for the WT and *as2-7D* mutant (*n =* 3 biological replicates). Each dot represents the angle of deviation for a given cell lineage. Data representation and statistical comparisons are shown in the same format as described for panels E and J. Exact *p-values* for epidermal angle of deviation: WT outer vs inner, 3.793354 × 10^-05^; *as2-7D* outer vs inner, 1.572509 × 10^-20^; WT vs *as2-7D* outer, 8.052118 × 10^-10^; WT vs *as2-7D* inner, 0.349.

**Fig. S6.**
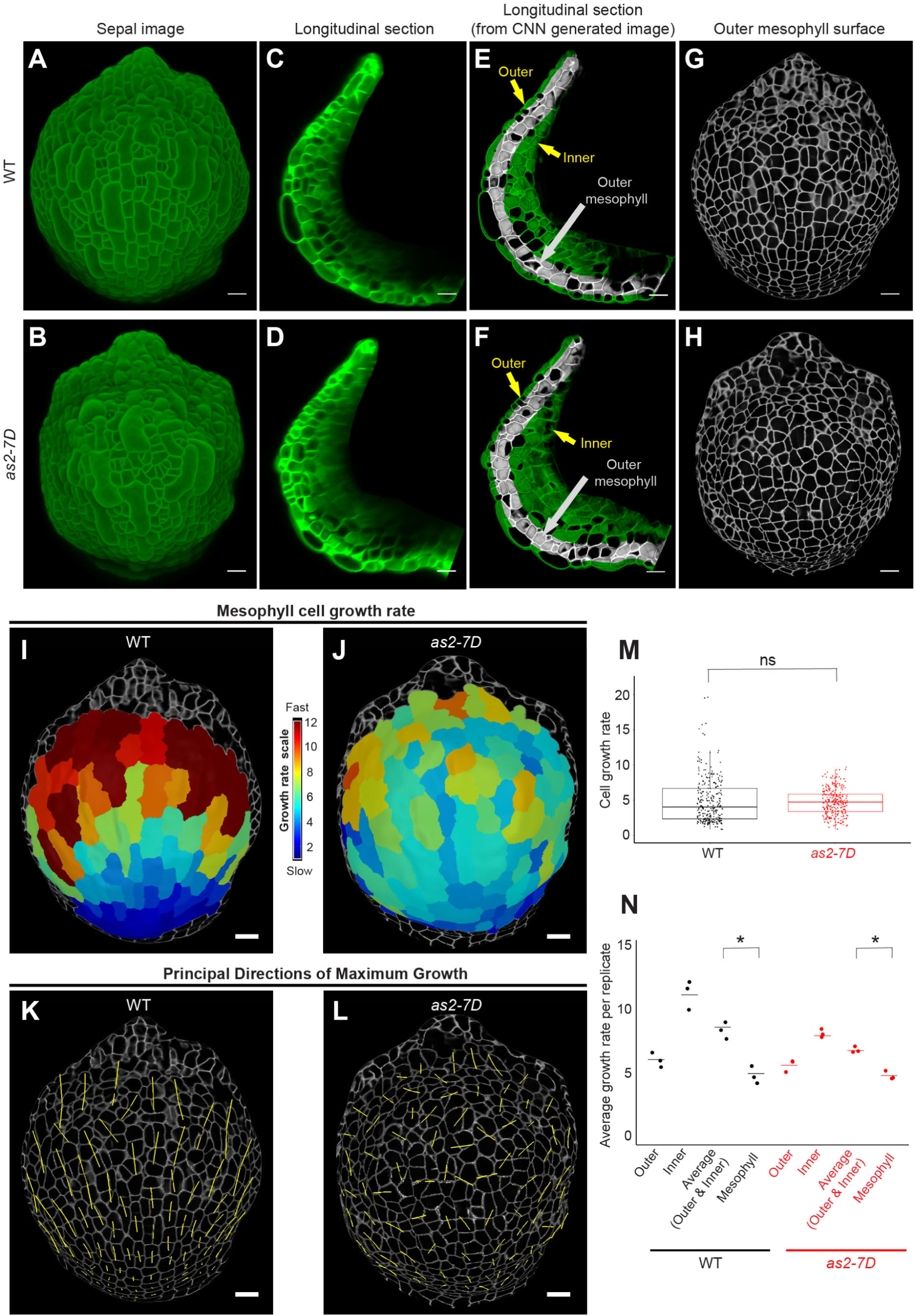
Growth patterns of the mesophyll cells mimic that of the outer epidermis for both wild type (WT) and *as2-7D*, associated with Main Fig. 3. (**A** to **B**) Processed confocal images of (**A**) the wild-type (WT) and (**B**) *as2-7D* mutant stage 7 sepals, harboring the pLH13 membrane marker, at the 48-hour time point (representative of *n = 3*). Scalebars, 20 µm. (**C** to **D**) View of the longitudinal cross section through the center of the stage 7 sepal blade for (**C**) the WT and (**D**) *as2-7D* mutant (representative of *n = 3*). Scalebars, 20 µm. (**E** to **F)** Longitudinal sections of stage 7 sepal images of (**E**) the WT and (**F**) *as2-7D* mutant processed with CNN (Unet display/Vijayan.Unet algorithm, see Materials and Methods), to enhance the cell outlines (representative of *n = 3*). The gray areas mark the mesophyll layer closest to the outer epidermis (outer mesophyll), generated after running (Mesh Interaction/Annihilate) process in MorphographX to remove the outer epidermis. The annihilated image (gray) was overlayed on the CNN generated image (green) for comparison. The outer and inner epidermal layers are shown (yellow arrows). Scalebars, 20 µm. (**G** to **H**) Plasma membrane signal from the outer mesophyll layer of (**G**) to WT and (**H**) the *as2-7D* mutant projected onto the mesh surface generated using MorphographX. Scalebars, 20 µm. (**I** to **J**) Growth rates of the mesophyll cells over a 48-hour period for (**I**) the WT and (**J**) *as2-7D* mutant are shown (representative of *n = 3*). Red to blue heatmap gradient shows fast to slow growth. Scalebars, 20 µm. (**K** to **L**) Principal directions of maximal growth (PDGmax, yellow) of the mesophyll cells of (**K**) WT and (**L**) *as2-7D* mutant are shown. Segmented cells in grey are superimposed on the plasma membrane signal (also in grey). Note that both the growth rates (panels I to J) and growth directions of the mesophyll cells are similar to those of the outer epidermis, as shown in Main Figure 3. Scalebars, 20 µm. (**M**) Graph shows the cell growth rates of the mesophylls cells over a 48-hour period for the WT (black) and *as2-7D* (red) mutant (*n = 3* biological replicates). Each dot represents the growth rate for a given cell lineage. The dot plots are overlayed with box plots, where the central line represents the median. The p-value from Tukey’s test (two-tailed) is shown. *p-values*: ns = non-significant (*p> 0.05*). (**N**) Graph shows the average cell growth rates of the outer epidermis (Outer), inner epidermis (inner), average of the outer and inner epidermis and the mesophyll cells of the WT (black) and *as2-7D* (red) mutant. Each dot represents the average growth rate across all cell lineages in each replicate (*n = 3* biological replicates). The lines represent the mean growth rate across three biological replicates. The p-value from Tukey’s test (two-tailed) is shown. *p-values*: *<0.05, ns = non-significant. Exact p-values for WT average (Outer & Inner) vs Mesophyll, 0.0036; WT average (Outer & Inner) vs Mesophyll, 0.0036.

**Fig. S7.**
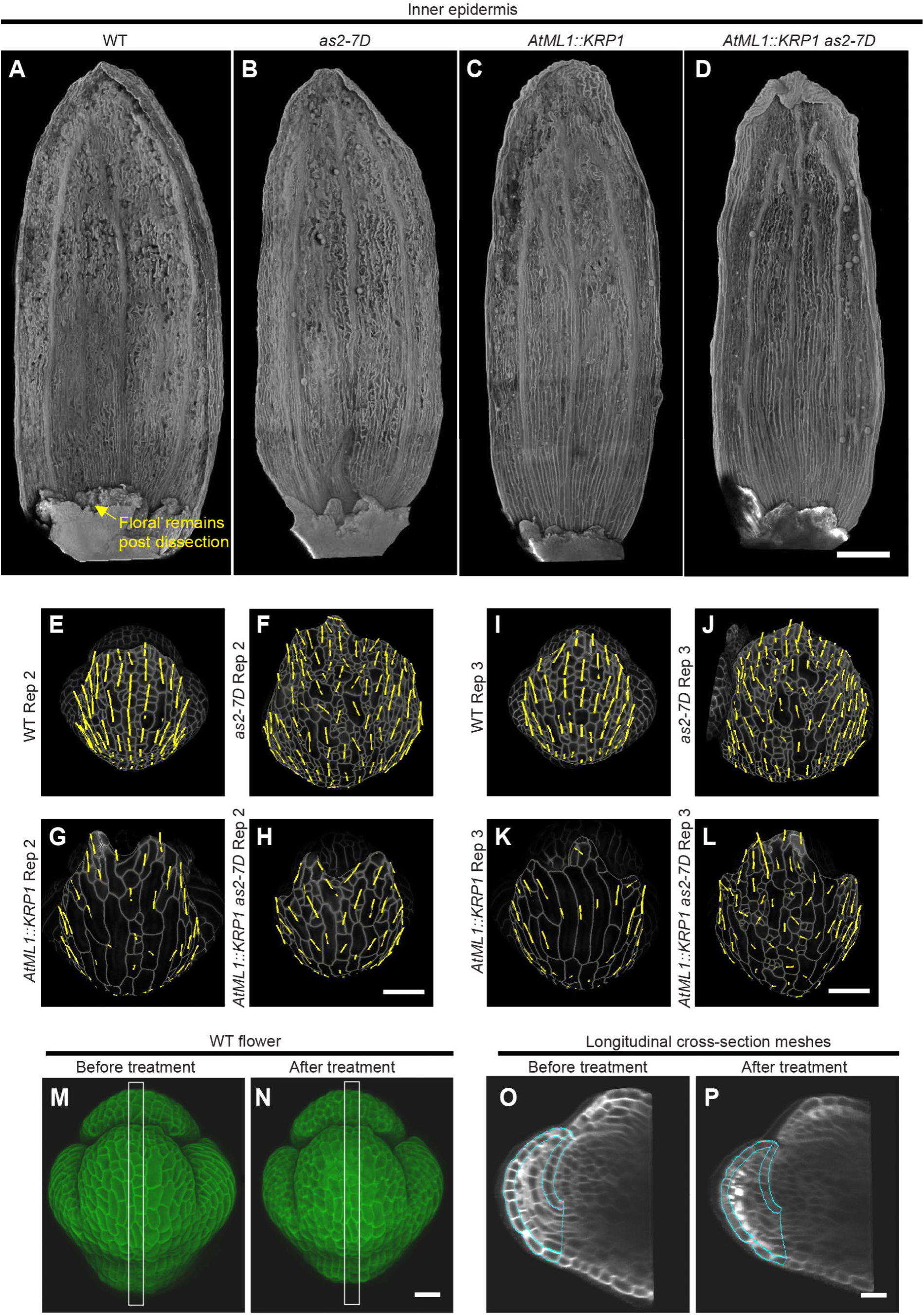
Analysis of cell growth direction and stiffness due to *KRP1* overexpression in wild type (WT) and *as2-7D*, associated with Main Fig. 5. (**A** to **D**) Confocal images of the inner epidermis of (**A**) Wild type (WT), (**B**) *as2-7D* mutant, (**C**) *AtML1::KRP1* and (**D**) *AtML1::KRP1 as2-7D* stage 14 sepals. The sepals were stained with PI. Note the smooth morphology of the inner epidermis across all genotypes. Scalebar, 200 µm, images representative of *n =* 3 adult sepals. (**E** to **L**) Principal direction of growth (PDG_max_) for the outer epidermal cells of (**E**) WT Replicate 2, (**F**) *as2-7D* mutant Replicate 2, (**G**) *AtML1::KRP1* Replicate 2, (**H**) *AtML1::KRP1 as2-7D* Replicate 2, (**I**) WT Replicate 3, (**J**) *as2-7D* mutant Replicate 3, (**K**) *AtML1::KRP1* Replicate 3 and (**L**) *AtML1::KRP1 as2-7D* Replicate 3 sepals. PDG_max_ tensors (yellow) calculated over a 48-hour live imaging period starting from stage 4, projected on a later time point surface. Segmented cells in grey are superimposed onto the plasma membrane signal (also in grey). Scalebar, 20 µm, *n =* 3. (**M** and **N**) Representative confocal images of WT (*n = 5*) early stage 6 flower (**M**) before and (**N**) after osmotic treatment in 0.35M NaCl. The PI (cell wall stain) and membrane marker (pLH13, *35S::mCitrine-RCI2A*) signals have been merged together and shown. The highlighted region of the outer sepal (white rectangle) indicates the comparable longitudinal section extracted for analyzing epidermal shrinkage. (**O** and **P**) MorphoGraphX generated surface of the longitudinal section (**O**) before treatment and (**P**) after osmotic treatment, extracted from flower images shown in panels M and N respectively. Comparable regions of the outer and inner sepal epidermis were segmented (outlined in cyan) to calculate the percentage areal shrinkage.

**Fig. S8.**
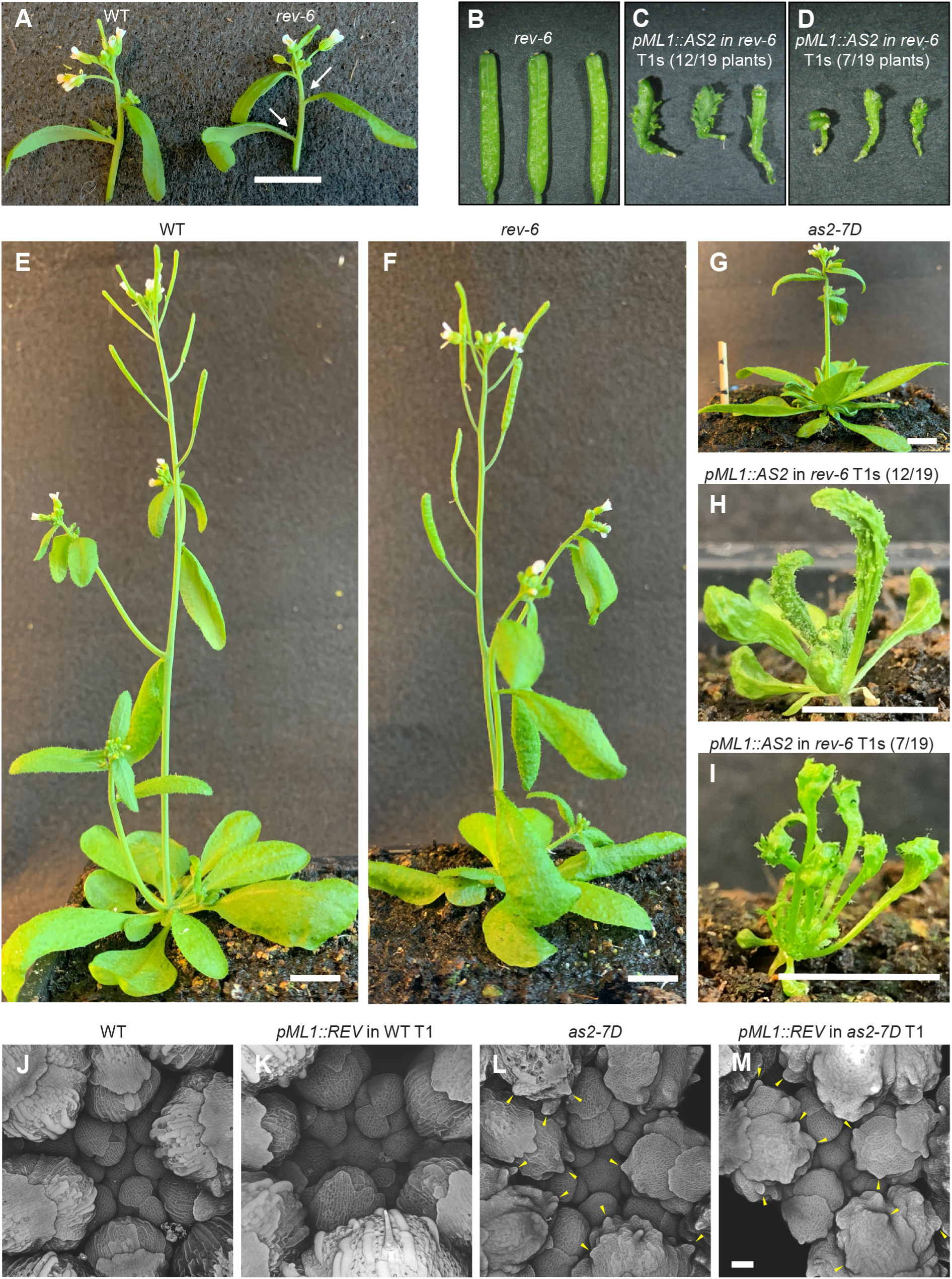
Ectopic *AS2* expression mediated buckles and subsequent outgrowth formation occur independent of *REVOLUTA*, associated with Main Fig. 6. (**A**) Representation of the previously characterized meristem initiation defect phenotype of *rev-6* mutant. Images show an excised lateral inflorescence meristem of the WT (left) and *rev-6* mutant (right). Majority of the cauline leaves axils (white arrows) in the *rev-6* mutant are bare, as they fail to produce axillary meristem, as described by Otsuga et. al. ^64^. Scalebar, 1cm. (**B** to **D**) Fully developed siliques of (**B**) *rev-6* and (**C** and **D**) *pML1::AS2* in rev-6 T1s. No of independent T1s assessed *n =* 19. The three siliques correspond to three independent plants. The numbers represent the total number of plants showing strong (12/19 plants, panel C) to extreme (7/19 plants, panel D) phenotypes, categorized based on the extent of leaf deformity in the adult vegetative stage. (**E** to **I**) Representative six-week-old adult plants of (**E**) WT, (**F**) *rev-6*, (**G**) *as2-7D* and (**H** and **I**) *pML1::AS2* in *rev-6* T1s. Total number of T1s assessed *n =* 19. The transgenics show strong (12/19 plants, panel H) to extreme (7/19 plants, panel I) defects in leaf development, characterized by outgrowth formation on the abaxial epidermis and stunted growth. Scalebar, 1 cm. (**J** to **M**) Confocal images of PI-stained (greyscale) inflorescences of (**J**) WT, (**K**) representative *pML1::REV* in WT T1, (**L**) *as2-7D* mutant, and (**M**) representative *pML1::REV* in *as2-7D* T1. Note that the transgenics ectopically expressing *REV* looks like their respective backgrounds in either the WT (panels J and K) or *as2-7D* (panels L and M). Yellow arrowheads point to the outgrowths. For each *pML1::REV* transgenics in WT and in *as2-7D* backgrounds, number of independent T1s assessed, *n =* 28. Scalebar, 50 µm.

**Fig. S9.**
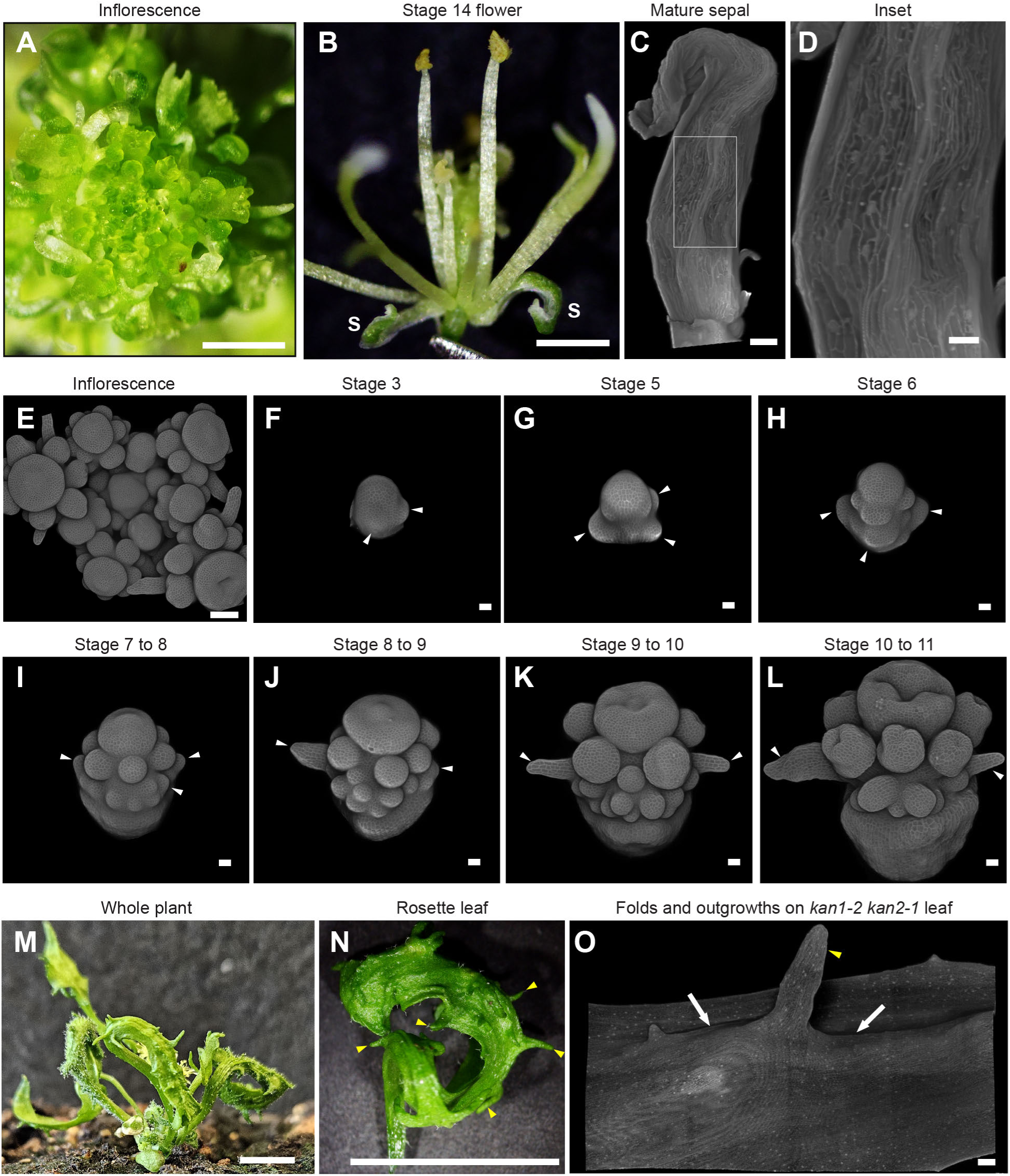
KANADI loss-of-function mutant (*kan1-2 kan2-1*) exhibits floral development defects which are distinct from *as2-7D*, associated with Main Fig. 6. (**A**) Whole inflorescence images of *kan1-2 kan2-1*. The image is representative of *n =* 3 inflorescences. Scalebar, 1 mm. (**B**) Stage 14 flower of *kan1-2 kan2-1*. The image is representative of *n =* 3 inflorescences. Scalebar, 1 mm. (**C**) Confocal image of PI-stained (greyscale) *kan1-2 kan2-1* mature sepal showing the outer (abaxial) epidermis. The image is representative of *n =* 3 sepals. Scalebar, 100 µm. Note the twisted and curled morphology of the sepal. (**D**) Inset of *kan1-2 kan2-1* mature sepal outlined in panel (C). Note the folds on the abaxial epidermis of *kan1-2 kan2-1* sepal, which are less severe as compared to *as2-7D*. Scalebar, 50 µm. (**E**) Confocal image of PI-stained (greyscale) *kan1-2 kan2-1* inflorescence. The image is representative of *n =* 3 inflorescences. (**F** to **L**) Confocal images of PI-stained (greyscale) *kan1-2 kan2-1* flowers of different developmental stages (staged based on floral organ development in accordance with Smyth et al., 1990 ^88^. The sepals are shown by white arrows. All the floral organs are morphologically deformed. Note that the sepals fail to enclose the floral meristem from initial stage onwards and exhibit a narrow blade development in the later stages. Parts of the inflorescence tissue not associated with the flowers were removed by image processing. Scalebar, 20 µm. (**M**) Representative five-week-old adult *kan1-2 kan2-1* plant. The image is representative of *n =* 3 plants. Scalebar, 1 cm. (**N**) Representative rosette leaf of *kan1-2 kan2-1*. Yellow arrowheads show the outgrowths. The image is representative of *n =* 3 leaves. Scalebar, 1 cm. (**O**) Confocal image showing a region of the abaxial epidermis of kan*1-2 kan2-1* leaf (PI-stained, grayscale). Yellow arrows show the outgrowth, while the white arrows denote the folds on the leaf. Note that the outgrowth forms in the region where the folds form. Scalebar, 200 µm.

**Fig. S10.**
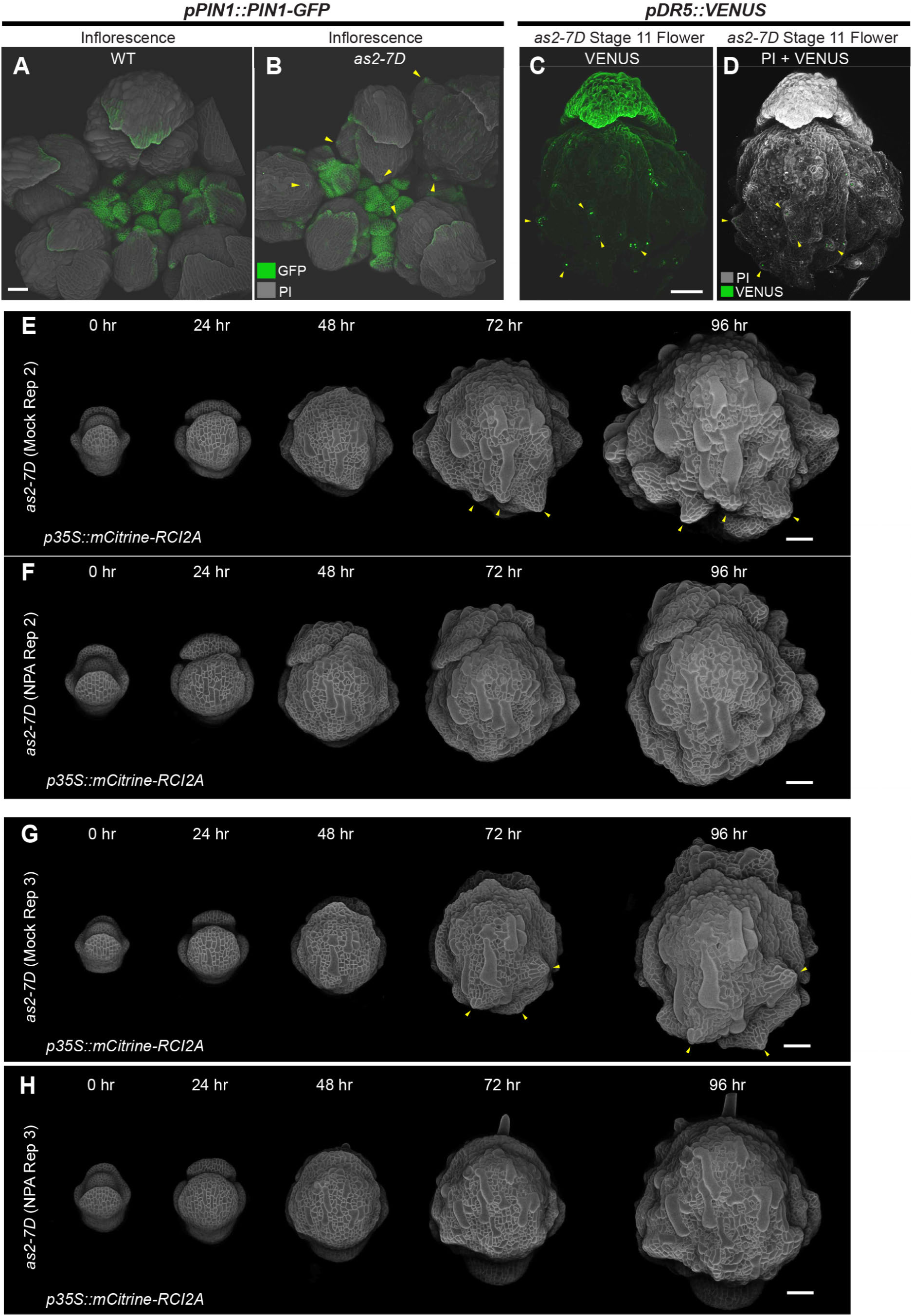
Outgrowths on *as2-7D* outer epidermis are formed due to buckling mediated PIN1 convergence and subsequent auxin maxima formation, associated with Main Fig. 7. (**A** and **B**) Confocal images show the expression pattern of PIN1 protein reporter (*pPIN1::PIN1-GFP*) in **(A)** wild-type (WT) and (**B**) *as2-7D* inflorescence meristems. Note the expression of *PIN1* directed towards the emerging outgrowths on the outer epidermis of *as2-7D* older flowers. Scalebar, 50 µm, *n =* 3. (**C** and **D**) Expression pattern of auxin response reporter (*pDR5::VENUS*) in stage 11 flower of *as2-7D* mutant. Shown is (**C**) VENUS signal by itself (green) and (**D**) VENUS signal merged with PI (grey). Note the presence of DR5 signal specifically in the outgrowths (yellow arrowheads). Scalebar, 100 µm, *n =* 3. (**E** to **H**) Representative live-imaging series showing the second replicate of (**E**) Mock treated, (**F**) 10µM NPA treated developing flowers, and third replicate of (**G**) Mock treated, (**H**) 10µM NPA treated developing flowers of *as2-7D* mutant. Images taken at every 24-hour interval over 96-hour period. Note the development of outgrowths from 72-hour time-point onwards (yellow arrowheads) in the mock treated *as2-7D* samples (panels E and G), which are absent in NPA treated *as2-7D* flowers (panels F and H). The *as2-7D* flowers harbor pLH13 (*35S::mCitrine-RCI2A*) cell membrane marker (greyscale). Trichomes were removed from the outer sepal epidermis by image processing. Scalebar, 50 µm.

**Supplementary Table S1:** List of primers used in this study.

**Supplementary Data S1:** Raw data for making the plots

**Supplementary Movie S1:** Confocal images of the outer and inner epidermis of adult sepals of the wild type (WT, left) and *as2-7D* mutant (right). Genotypes harbor pLH13 (*p35S::mCitrine-RCI2A*) plasma membrane marker. Scalebar, 200 µm.

**Supplementary Movie S2:** Confocal images of developing wild-type (WT, left) and *as2-7D* mutant (right) flowers imaged every 24 hours, over 96-hour time period. Genotypes harbor pLH13 (*p35S::mCitrine-RCI2A*) plasma membrane marker. Trichomes were removed from the outer sepal epidermis by image processing for better visualization of sepal morphology. Scalebar, 50 µm.

**Supplementary Movie S3:** Confocal images show the outer epidermis of adult sepals of wild type (WT, top left), *as2-7D* (top right), *pML1::KRP1* (bottom left) and *pML1::KRP1 as2-7D* (bottom right). Note the largely rescued sepal smoothness in *pML1::KRP1 as2-7D* compared to *as2-7D*, due to an increase in the longitudinally aligned giant cells. Sepals were stained with Propidium-iodide, which stains the cell walls. Scalebar, 200 µm.

**Supplementary Movie S4:** Confocal images of developing *as2-7D* mutant flowers grown in the presence of mock (left) or 10µM NPA (right). Images were captured every 24 hours over the 96-hour time-period. The *as2-7D* mutant harbors pLH13 (*p35S::mCitrine-RCI2A*) plasma membrane marker. Trichomes were removed from the outer sepal epidermis by image processing. Note the reduction in outgrowth formation in 10µM NPA-treated samples despite buckling. Scalebar, 50 µm.

## REFERENCES

1. Richman, D.P., Stewart, R.M., Hutchinson, J.W., and Caviness, V.S., Jr. (1975). Mechanical model of brain convolutional development. Science 189, 18–21. 10.1126/science.1135626.

2. Hannezo, E., Prost, J., and Joanny, J.F. (2011). Instabilities of monolayered epithelia: shape and structure of villi and crypts. Phys Rev Lett 107, 078104. 10.1103/PhysRevLett.107.078104.

3. Savin, T., Kurpios, N.A., Shyer, A.E., Florescu, P., Liang, H., Mahadevan, L., and Tabin, C.J. (2011). On the growth and form of the gut. Nature 476, 57–62. 10.1038/nature10277.

4. Shyer, A.E., Tallinen, T., Nerurkar, N.L., Wei, Z., Gil, E.S., Kaplan, D.L., Tabin, C.J., and Mahadevan, L. (2013). Villification: how the gut gets its villi. Science 342, 212–218. 10.1126/science.1238842.

5. Antoniou Kourounioti, R.L., Band, L.R., Fozard, J.A., Hampstead, A., Lovrics, A., Moyroud, E., Vignolini, S., King, J.R., Jensen, O.E., and Glover, B.J. (2013). Buckling as an origin of ordered cuticular patterns in flower petals. Journal of The Royal Society Interface 10, 20120847. doi:10.1098/rsif.2012.0847.

6. Moyroud, E., Airoldi, C.A., Ferria, J., Giorio, C., Steimer, S.S., Rudall, P.J., Prychid, C.J., Halliwell, S., Walker, J.F., Robinson, S., et al. (2022). Cuticle chemistry drives the development of diffraction gratings on the surface of Hibiscus trionum petals. Current Biology 32, 5323–5334.e5326. 10.1016/j.cub.2022.10.065.

7. Waddington, C.H. (1939). Preliminary Notes on the Development of the Wings in Normal and Mutant Strains of Drosophila. Proc Natl Acad Sci U S A 25, 299–307. 10.1073/pnas.25.7.299.

8. Thomson, J.A. (1917). On Growth and Form. Nature 100, 21–22. 10.1038/100021a0.

9. Tsukaya, H. (2005). Leaf shape: genetic controls and environmental factors. Int J Dev Biol 49, 547–555. 10.1387/ijdb.041921ht.

10. Tsukaya, H. (2006). Mechanism of leaf-shape determination. Annu Rev Plant Biol 57, 477–496. 10.1146/annurev.arplant.57.032905.105320.

11. Boudaoud, A. (2010). An introduction to the mechanics of morphogenesis for plant biologists. Trends Plant Sci 15, 353–360. 10.1016/j.tplants.2010.04.002.

12. Li, B., Cao, Y.-P., Feng, X.-Q., and Gao, H. (2012). Mechanics of morphological instabilities and surface wrinkling in soft materials: a review. Soft Matter 8, 5728–5745. 10.1039/C2SM00011C.

13. Nelson, C.M. (2016). On Buckling Morphogenesis. J Biomech Eng 138, 021005. 10.1115/1.4032128.

14. Borowska-Wykręt, D., and Kwiatkowska, D. (2018). Folding, Wrinkling, and Buckling in Plant Cell Walls. In Plant Biomechanics: From Structure to Function at Multiple Scales, A. Geitmann, and J. Gril, eds. (Springer International Publishing), pp. 209–233. 10.1007/978-3-319-79099-2_10.

15. Shyer, A.E., Rodrigues, A.R., Schroeder, G.G., Kassianidou, E., Kumar, S., and Harland, R.M. (2017). Emergent cellular self-organization and mechanosensation initiate follicle pattern in the avian skin. Science 357, 811–815. 10.1126/science.aai7868.

16. Varner, V.D., Gleghorn, J.P., Miller, E., Radisky, D.C., and Nelson, C.M. (2015). Mechanically patterning the embryonic airway epithelium. Proceedings of the National Academy of Sciences 112, 9230–9235. doi:10.1073/pnas.1504102112.

17. Maroudas-Sacks, Y., and Keren, K. (2021). Mechanical Patterning in Animal Morphogenesis. Annual Review of Cell and Developmental Biology 37, 469–493. 10.1146/annurev-cellbio-120319-030931.

18. Silverberg, J.L., Noar, R.D., Packer, M.S., Harrison, M.J., Henley, C.L., Cohen, I., and Gerbode, S.J. (2012). 3D imaging and mechanical modeling of helical buckling in Medicago truncatula plant roots. Proc Natl Acad Sci U S A 109, 16794–16799. 10.1073/pnas.1209287109.

19. Yan, J., Wang, B., Zhou, Y., and Hao, S. (2018). Resistance from agar medium impacts the helical growth of Arabidopsis primary roots. Journal of the Mechanical Behavior of Biomedical Materials 85, 43–50. 10.1016/j.jmbbm.2018.05.018.

20. Sachse, R., Westermeier, A., Mylo, M., Nadasdi, J., Bischoff, M., Speck, T., and Poppinga, S. (2020). Snapping mechanics of the Venus flytrap (Dionaea muscipula). Proc Natl Acad Sci U S A 117, 16035–16042. 10.1073/pnas.2002707117.

21. Forterre, Y., Skotheim, J.M., Dumais, J., and Mahadevan, L. (2005). How the Venus flytrap snaps. Nature 433, 421–425. 10.1038/nature03185.

22. Airoldi, C.A., Lugo, C.A., Wightman, R., Glover, B.J., and Robinson, S. (2021). Mechanical buckling can pattern the light-diffracting cuticle of Hibiscus trionum. Cell Reports 36, 109715. 10.1016/j.celrep.2021.109715.

23. Green, P.B. (1992). Pattern Formation in Shoots: A Likely Role for Minimal Energy Configurations of the Tunica. International Journal of Plant Sciences 153, S59–S75. 10.1086/297064.

24. Dumais, J., and Steele, C.R. (2000). New Evidence for the Role of Mechanical Forces in the Shoot Apical Meristem. Journal of Plant Growth Regulation 19, 7–18. 10.1007/s003440000003.

25. Rambaud-Lavigne, L., Chatterjee, A., Bovio, S., Battu, V., Lavigne, Q., Gundiah, N., Boudaoud, A., and Das, P. (2024). Heterogeneous identity, stiffness and growth characterise the shoot apex of Arabidopsis stem cell mutants. Development 151. 10.1242/dev.202810.

26. Żebrowski, J. (1999). Dynamic behaviour of inflorescence-bearing Triticale and Triticum stems. Planta 207, 410–417.

27. Fournier, M., Dlouhá, J., Jaouen, G., and Almeras, T. (2013). Integrative biomechanics for tree ecology: beyond wood density and strength. Journal of Experimental Botany 64, 4793–4815. 10.1093/jxb/ert279.

28. Chapotin, S.M., Razanameharizaka, J.H., and Holbrook, N.M. (2006). A biomechanical perspective on the role of large stem volume and high water content in baobab trees (Adansonia spp.; Bombacaceae). Am J Bot 93, 1251–1264. 10.3732/ajb.93.9.1251.

29. Méndez-Alonzo, R., Moctezuma, C., Ordoñez, V.R., Angeles, G., Martínez, A.J., and López-Portillo, J. (2015). Root biomechanics in Rhizophora mangle: anatomy, morphology and ecology of mangrove’s flying buttresses. Annals of Botany 115, 833–840. 10.1093/aob/mcv002.

30. Huang, C., Wang, Z., Quinn, D., Suresh, S., and Hsia, K.J. (2018). Differential growth and shape formation in plant organs. Proceedings of the National Academy of Sciences 115, 12359–12364. doi:10.1073/pnas.1811296115.

31. Derr, J., Bastien, R., Couturier, É., and Douady, S. (2018). Fluttering of growing leaves as a way to reach flatness: experimental evidence on *Persea americana*. Journal of The Royal Society Interface 15, 20170595. doi:10.1098/rsif.2017.0595.

32. Wang, T., Fu, C., Potier-Ferry, M., and Xu, F. (2024). Morphomechanics of growing curled petals and leaves. Journal of the Mechanics and Physics of Solids 184, 105534. 10.1016/j.jmps.2023.105534.

33. Whitewoods, C.D. (2021). Riddled with holes: Understanding air space formation in plant leaves. PLoS Biol 19, e3001475. 10.1371/journal.pbio.3001475.

34. Evans, J.R. (2013). Improving photosynthesis. Plant Physiol 162, 1780–1793. 10.1104/pp.113.219006.

35. Roeder, A.H.K. (2021). Arabidopsis sepals: A model system for the emergent process of morphogenesis. Quantitative Plant Biology 2, e14, e14. 10.1017/qpb.2021.12.

36. Hong, L., Dumond, M., Tsugawa, S., Sapala, A., Routier-Kierzkowska, A.L., Zhou, Y., Chen, C., Kiss, A., Zhu, M., Hamant, O., et al. (2016). Variable Cell Growth Yields Reproducible OrganDevelopment through Spatiotemporal Averaging. Dev Cell 38, 15–32. 10.1016/j.devcel.2016.06.016.

37. Iwakawa, H., Ueno, Y., Semiarti, E., Onouchi, H., Kojima, S., Tsukaya, H., Hasebe, M., Soma, T., Ikezaki, M., Machida, C., and Machida, Y. (2002). The ASYMMETRIC LEAVES2 gene of Arabidopsis thaliana, required for formation of a symmetric flat leaf lamina, encodes a member of a novel family of proteins characterized by cysteine repeats and a leucine zipper. Plant Cell Physiol 43, 467–478. 10.1093/pcp/pcf077.

38. Bowman, J.L., Eshed, Y., and Baum, S.F. (2002). Establishment of polarity in angiosperm lateral organs. Trends Genet 18, 134–141. 10.1016/s0168-9525(01)02601-4.

39. Lin, W.C., Shuai, B., and Springer, P.S. (2003). The Arabidopsis LATERAL ORGAN BOUNDARIES-domain gene ASYMMETRIC LEAVES2 functions in the repression of KNOX gene expression and in adaxial-abaxial patterning. Plant Cell 15, 2241–2252. 10.1105/tpc.014969.

40. Scacchi, E., Paszkiewicz, G., Thi Nguyen, K., Meda, S., Burian, A., de Back, W., and Timmermans, M.C.P. (2024). A diffusible small-RNA-based Turing system dynamically coordinates organ polarity. Nature Plants 10, 412–422. 10.1038/s41477-024-01634-x.

41. Wu, G., Lin, W.C., Huang, T., Poethig, R.S., Springer, P.S., and Kerstetter, R.A. (2008). KANADI1 regulates adaxial-abaxial polarity in Arabidopsis by directly repressing the transcription of ASYMMETRIC LEAVES2. Proc Natl Acad Sci U S A 105, 16392–16397. 10.1073/pnas.0803997105.

42. Eshed, Y., Baum, S.F., Perea, J.V., and Bowman, J.L. (2001). Establishment of polarity in lateral organs of plants. Curr Biol 11, 1251–1260. 10.1016/s0960-9822(01)00392-x.

43. Kerstetter, R.A., Bollman, K., Taylor, R.A., Bomblies, K., and Poethig, R.S. (2001). KANADI regulates organ polarity in Arabidopsis. Nature 411, 706–709. 10.1038/35079629.

44. Lin, X., Gu, D., Zhao, H., Peng, Y., Zhang, G., Yuan, T., Li, M., Wang, Z., Wang, X., and Cui, S. (2018). LFR is functionally associated with AS2 to mediate leaf development in Arabidopsis. Plant J. 10.1111/tpj.13973.

45. Hervieux, N., Dumond, M., Sapala, A., Routier-Kierzkowska, A.L., Kierzkowski, D., Roeder, A.H., Smith, R.S., Boudaoud, A., and Hamant, O. (2016). A Mechanical Feedback Restricts Sepal Growth and Shape in Arabidopsis. Curr Biol. 10.1016/j.cub.2016.03.004.

46. Onoda, Y., Schieving, F., and Anten, N.P. (2015). A novel method of measuring leaf epidermis and mesophyll stiffness shows the ubiquitous nature of the sandwich structure of leaf laminas in broad-leaved angiosperm species. J Exp Bot 66, 2487–2499. 10.1093/jxb/erv024.

47. Vijayan, A., Tofanelli, R., Strauss, S., Cerrone, L., Wolny, A., Strohmeier, J., Kreshuk, A., Hamprecht, F.A., Smith, R.S., and Schneitz, K. (2021). A digital 3D reference atlas reveals cellular growth patterns shaping the Arabidopsis ovule. eLife 10, e63262. 10.7554/eLife.63262.

48. Mollier, C., Skrzydeł, J., Borowska-Wykręt, D., Majda, M., Bayle, V., Battu, V., Totozafy, J.C., Dulski, M., Fruleux, A., Wrzalik, R., et al. (2023). Spatial consistency of cell growth direction during organ morphogenesis requires CELLULOSE SYNTHASE INTERACTIVE1. Cell Rep 42, 112689. 10.1016/j.celrep.2023.112689.

49. Harmansa, S., Erlich, A., Eloy, C., Zurlo, G., and Lecuit, T. (2023). Growth anisotropy of the extracellular matrix shapes a developing organ. Nature Communications 14, 1220. 10.1038/s41467-023-36739-y.

50. Hong, L., Dumond, M., Zhu, M., Tsugawa, S., Li, C.-B., Boudaoud, A., Hamant, O., and Roeder, A.H.K. (2018). Heterogeneity and Robustness in Plant Morphogenesis: From Cells to Organs. Annual Review of Plant Biology 69, 469–495. 10.1146/annurev-arplant-042817-040517.

51. Sylvester, A.W., Williams, M.H., and Green, P.B. (1989). Orientation of cortical microtubules correlates with cell shape and division direction. Protoplasma 153, 91–103. 10.1007/BF01322469.

52. Hamant, O., Heisler, M.G., Jönsson, H., Krupinski, P., Uyttewaal, M., Bokov, P., Corson, F., Sahlin, P., Boudaoud, A., Meyerowitz, E.M., et al. (2008). Developmental patterning by mechanical signals in Arabidopsis. Science 322, 1650–1655. 10.1126/science.1165594.

53. Roeder, A.H., Chickarmane, V., Cunha, A., Obara, B., Manjunath, B.S., and Meyerowitz, E.M. (2010). Variability in the control of cell division underlies sepal epidermal patterning in Arabidopsis thaliana. PLoS Biol 8, e1000367. 10.1371/journal.pbio.1000367.

54. Bemis, S.M., and Torii, K.U. (2007). Autonomy of cell proliferation and developmental programs during Arabidopsis aboveground organ morphogenesis. Dev Biol 304, 367–381. 10.1016/j.ydbio.2006.12.049.

55. Sapala, A., and Smith, R.S. (2020). Osmotic Treatment for Quantifying Cell Wall Elasticity in the Sepal of Arabidopsis thaliana. Methods Mol Biol 2094, 101–112. 10.1007/978-1-0716-0183-9_11.

56. Kierzkowski, D., Nakayama, N., Routier-Kierzkowska, A.L., Weber, A., Bayer, E., Schorderet, M., Reinhardt, D., Kuhlemeier, C., and Smith, R.S. (2012). Elastic domains regulate growth and organogenesis in the plant shoot apical meristem. Science 335, 1096–1099. 10.1126/science.1213100.

57. Waites, R., and Hudson, A. (1995). phantastica: a gene required for dorsoventrality of leaves in Antirrhinum majus. Development 121, 2143–2154. 10.1242/dev.121.7.2143.

58. Fukushima, K., and Hasebe, M. (2014). Adaxial-abaxial polarity: the developmental basis of leaf shape diversity. Genesis 52, 1–18. 10.1002/dvg.22728.

59. Izhaki, A., and Bowman, J.L. (2007). KANADI and Class III HD-Zip Gene Families Regulate Embryo Patterning and Modulate Auxin Flow during Embryogenesis in Arabidopsis. The Plant Cell 19, 495–508. 10.1105/tpc.106.047472.

60. Eshed, Y., Izhaki, A., Baum, S.F., Floyd, S.K., and Bowman, J.L. (2004). Asymmetric leaf development and blade expansion in Arabidopsis are mediated by KANADI and YABBY activities. Development 131, 2997–3006. 10.1242/dev.01186.

61. Caggiano, M.P., Yu, X., Bhatia, N., Larsson, A., Ram, H., Ohno, C.K., Sappl, P., Meyerowitz, E.M., Jönsson, H., and Heisler, M.G. (2017). Cell type boundaries organize plant development. eLife 6, e27421. 10.7554/eLife.27421.

62. Nakata, M., and Okada, K. (2012). The three-domain model: a new model for the early development of leaves in Arabidopsis thaliana. Plant Signal Behav 7, 1423–1427. 10.4161/psb.21959.

63. Guan, C., Wu, B., Yu, T., Wang, Q., Krogan, N.T., Liu, X., and Jiao, Y. (2017). Spatial Auxin Signaling Controls Leaf Flattening in Arabidopsis. Current Biology 27, 2940–2950.e2944. 10.1016/j.cub.2017.08.042.

64. Otsuga, D., DeGuzman, B., Prigge, M.J., Drews, G.N., and Clark, S.E. (2001). REVOLUTA regulates meristem initiation at lateral positions. Plant J 25, 223–236. 10.1046/j.1365-313x.2001.00959.x.

65. Watanabe, K., and Okada, K. (2003). Two discrete cis elements control the Abaxial side-specific expression of the FILAMENTOUS FLOWER gene in Arabidopsis. Plant Cell 15, 2592–2602. 10.1105/tpc.015214.

66. Semiarti, E., Ueno, Y., Tsukaya, H., Iwakawa, H., Machida, C., and Machida, Y. (2001). The ASYMMETRIC LEAVES2 gene of Arabidopsis thaliana regulates formation of a symmetric lamina, establishment of venation and repression of meristem-related homeobox genes in leaves. Development 128, 1771–1783. 10.1242/dev.128.10.1771.

67. Gälweiler, L., Guan, C., Müller, A., Wisman, E., Mendgen, K., Yephremov, A., and Palme, K. (1998). Regulation of polar auxin transport by AtPIN1 in Arabidopsis vascular tissue. Science 282, 2226–2230. 10.1126/science.282.5397.2226.

68. Abley, K., Sauret-Güeto, S., Marée, A.F., and Coen, E. (2016). Formation of polarity convergences underlying shoot outgrowths. Elife 5. 10.7554/eLife.18165.

69. Reinhardt, D., Pesce, E.-R., Stieger, P., Mandel, T., Baltensperger, K., Bennett, M., Traas, J., Friml, J., and Kuhlemeier, C. (2003). Regulation of phyllotaxis by polar auxin transport. Nature 426, 255–260. 10.1038/nature02081.

70. Heisler, M.G., Hamant, O., Krupinski, P., Uyttewaal, M., Ohno, C., Jönsson, H., Traas, J., and Meyerowitz, E.M. (2010). Alignment between PIN1 polarity and microtubule orientation in the shoot apical meristem reveals a tight coupling between morphogenesis and auxin transport. PLoS Biol 8, e1000516. 10.1371/journal.pbio.1000516.

71. Hamant, O., Meyerowitz, E.M., and Traas, J. (2011). Is cell polarity under mechanical control in plants? Plant Signal Behav 6, 137–139. 10.4161/psb.6.1.14269.

72. Nakayama, N., Smith, Richard S., Mandel, T., Robinson, S., Kimura, S., Boudaoud, A., and Kuhlemeier, C. (2012). Mechanical Regulation of Auxin-Mediated Growth. Current Biology 22, 1468–1476. 10.1016/j.cub.2012.06.050.

73. Kabir, A.M.R., Inoue, D., Afrin, T., Mayama, H., Sada, K., and Kakugo, A. (2015). Buckling of Microtubules on a 2D Elastic Medium. Scientific Reports 5, 17222. 10.1038/srep17222.

74. Sanyal, A., and Han, H.C. (2015). Artery buckling affects the mechanical stress in atherosclerotic plaques. Biomed Eng Online 14 *Suppl 1*, S4. 10.1186/1475-925x-14-s1-s4.

75. Roeder, A.H.K. (2021). Arabidopsis sepals: A model system for the emergent process of morphogenesis. Quant Plant Biol 2. 10.1017/qpb.2021.12.

76. Xu, L., Xu, Y., Dong, A., Sun, Y., Pi, L., Xu, Y., and Huang, H. (2003). Novel as1 and as2 defects in leaf adaxial-abaxial polarity reveal the requirement for ASYMMETRIC LEAVES1 and 2 and ERECTA functions in specifying leaf adaxial identity. Development 130, 4097–4107. 10.1242/dev.00622.

77. Friesen, S., and Hariharan, I.K. (2023). Coordinated growth of linked epithelia is mediated by the Hippo pathway. bioRxiv, 2023.2002.2026.530099. 10.1101/2023.02.26.530099.

78. Kelly-Bellow, R., Lee, K., Kennaway, R., Barclay, J.E., Whibley, A., Bushell, C., Spooner, J., Yu, M., Brett, P., Kular, B., et al. (2023). Brassinosteroid coordinates cell layer interactions in plants via cell wall and tissue mechanics. Science 380, 1275–1281. doi:10.1126/science.adf0752.

79. Du, J., Kirui, A., Huang, S., Wang, L., Barnes, W.J., Kiemle, S.N., Zheng, Y., Rui, Y., Ruan, M., Qi, S., et al. (2020). Mutations in the Pectin Methyltransferase QUASIMODO2 Influence Cellulose Biosynthesis and Wall Integrity in Arabidopsis. The Plant Cell 32, 3576–3597. 10.1105/tpc.20.00252.

80. Qi, J., Wu, B., Feng, S., Lü, S., Guan, C., Zhang, X., Qiu, D., Hu, Y., Zhou, Y., Li, C., et al. (2017). Mechanical regulation of organ asymmetry in leaves. Nat Plants 3, 724–733. 10.1038/s41477-017-0008-6.

81. Enugutti, B., Kirchhelle, C., and Schneitz, K. (2013). On the genetic control of planar growth during tissue morphogenesis in plants. Protoplasma 250, 651–661. 10.1007/s00709-012-0452-0.

82. Braybrook, S.A., and Peaucelle, A. (2013). Mechano-chemical aspects of organ formation in Arabidopsis thaliana: the relationship between auxin and pectin. PLoS One 8, e57813. 10.1371/journal.pone.0057813.

83. Kuhlemeier, C. (2007). Phyllotaxis. Trends Plant Sci 12, 143–150. 10.1016/j.tplants.2007.03.004.

84. Yadav, A.S., and Roeder, A.H.K. (2024). An optimized live imaging and growth analysis approach for Arabidopsis Sepals. bioRxiv, 2024.2001.2022.576735. 10.1101/2024.01.22.576735.

85. Routier-Kierzkowska, A.L., and Smith, R.S. (2013). Measuring the mechanics of morphogenesis. Curr Opin Plant Biol 16, 25–32. 10.1016/j.pbi.2012.11.002.

86. Liu, S., Strauss, S., Adibi, M., Mosca, G., Yoshida, S., Dello Ioio, R., Runions, A., Andersen, T.G., Grossmann, G., Huijser, P., et al. (2022). Cytokinin promotes growth cessation in the Arabidopsis root. Current Biology 32, 1974–1985.e1973. 10.1016/j.cub.2022.03.019.

87. Natonik-Białoń, S., Borowska-Wykręt, D., Mosca, G., Grelowski, M., Wrzalik, R., Smith, R.S., and Kwiatkowska, D. (2020). Deformation of a cell monolayer due to osmotic treatment: a case study of onion scale epidermis. Botany 98, 21–36. 10.1139/cjb-2019-0027.

88. Smyth, D.R., Bowman, J.L., and Meyerowitz, E.M. (1990). Early flower development in Arabidopsis. The Plant Cell 2, 755–767. 10.1105/tpc.2.8.755.

89. Robinson, D.O., Coate, J.E., Singh, A., Hong, L., Bush, M., Doyle, J.J., and Roeder, A.H.K. (2018). Ploidy and Size at Multiple Scales in the Arabidopsis Sepal. The Plant Cell 30, 2308–2329. 10.1105/tpc.18.00344.

90. Strauss, S., Runions, A., Lane, B., Eschweiler, D., Bajpai, N., Trozzi, N., Routier-Kierzkowska, A.-L., Yoshida, S., Rodrigues da Silveira, S., Vijayan, A., et al. (2022). Using positional information to provide context for biological image analysis with MorphoGraphX 2.0. eLife 11, e72601. 10.7554/eLife.72601.

91. Armezzani, A., Abad, U., Ali, O., Andres Robin, A., Vachez, L., Larrieu, A., Mellerowicz, E.J., Taconnat, L., Battu, V., Stanislas, T., et al. (2018). Transcriptional induction of cell wall remodelling genes is coupled to microtubule-driven growth isotropy at the shoot apex in Arabidopsis. Development 145. 10.1242/dev.162255.

92. Heisler, M.G., Ohno, C., Das, P., Sieber, P., Reddy, G.V., Long, J.A., and Meyerowitz, E.M. (2005). Patterns of Auxin Transport and Gene Expression during Primordium Development Revealed by Live Imaging of the Arabidopsis Inflorescence Meristem. Current Biology 15, 1899–1911. 10.1016/j.cub.2005.09.052.

93. Eshed, Y., Baum, S.F., Perea, J.V., and Bowman, J.L. (2001). Establishment of polarity in lateral organs of plants. Current Biology 11, 1251–1260. 10.1016/S0960-9822(01)00392-X.

94. Hong, L., Brown, J., Segerson, N.A., Rose, J.K., and Roeder, A.H. (2017). CUTIN SYNTHASE 2 Maintains Progressively Developing Cuticular Ridges in Arabidopsis Sepals. Mol Plant 10, 560–574. 10.1016/j.molp.2017.01.002.

95. Roeder, A.H., Cunha, A., Ohno, C.K., and Meyerowitz, E.M. (2012). Cell cycle regulates cell type in the Arabidopsis sepal. Development 139, 4416–4427. 10.1242/dev.082925.

96. Lukowitz, W., Gillmor, C.S., and Scheible, W.-R.d. (2000). Positional Cloning in Arabidopsis. Why It Feels Good to Have a Genome Initiative Working for You1. Plant Physiology 123, 795–806. 10.1104/pp.123.3.795.

97. Robinson, D.O., Coate, J.E., Singh, A., Hong, L., Bush, M., Doyle, J.J., and Roeder, A.H.K. (2018). Ploidy and Size at Multiple Scales in the Arabidopsis Sepal. Plant Cell 30, 2308–2329. 10.1105/tpc.18.00344.

98. Karimi, M., Inze, D., and Depicker, A. (2002). GATEWAY vectors for Agrobacterium-mediated plant transformation. Trends Plant Sci 7, 193–195.

99. Zhu, M., Chen, W., Mirabet, V., Hong, L., Bovio, S., Strauss, S., Schwarz, E.M., Tsugawa, S., Wang, Z., Smith, R.S., et al. (2020). Robust organ size requires robust timing of initiation orchestrated by focused auxin and cytokinin signalling. Nat Plants 6, 686–698. 10.1038/s41477-020-0666-7.

100. Barrell, P.J., and Conner, A.J. (2006). Minimal T-DNA vectors suitable for agricultural deployment of transgenic plants. BioTechniques 41, 708–710. 10.2144/000112306.

101. Wise, A.A., Liu, Z., and Binns, A.N. (2006). Three methods for the introduction of foreign DNA into Agrobacterium. Methods Mol Biol 343, 43–53. 10.1385/1-59745-130-4:43.

102. Zhu, M., and Roeder, A.H. (2020). Plants are better engineers: the complexity of plant organ morphogenesis. Curr Opin Genet Dev 63, 16–23. 10.1016/j.gde.2020.02.008.

103. Kiss, A., Moreau, T., Mirabet, V., Calugaru, C.I., Boudaoud, A., and Das, P. (2017). Segmentation of 3D images of plant tissues at multiple scales using the level set method. Plant Methods 13, 114. 10.1186/s13007-017-0264-5.

104. Huang, Z.Y., Hong, W., and Suo, Z. (2005). Nonlinear analyses of wrinkles in a film bonded to a compliant substrate. Journal of the Mechanics and Physics of Solids 53, 2101–2118. 10.1016/j.jmps.2005.03.007.

105. Audoly, B., and Boudaoud, A. (2008). Buckling of a stiff film bound to a compliant substrate—Part I:: Formulation, linear stability of cylindrical patterns, secondary bifurcations. Journal of the Mechanics and Physics of Solids 56, 2401–2421. 10.1016/j.jmps.2008.03.003.

106. Goriely, A. (2016). Morpho-elasticity: The mechanics and mathematics of biological growth (Springer New York).

107. Alnæs, M., Blechta, J., Hake, J., Johansson, A., Kehlet, B., Logg, A., Richardson, C., Ring, J., Rognes, M., and Wells, G. (2015). The FEniCS project version 1.5. 3. 10.11588/ans.2015.100.20553.

108. Gacon, F., Godin, C., and Ali, O. (2021). BVPy: A FEniCS-based Python package to ease the expression and study of boundary value problems in Biology. Journal of Open Source Software 6, 1–6. 10.21105/joss.02831.

